# Odor preference maps to cohesive transcriptional domains in the olfactory bulb

**DOI:** 10.64898/2026.06.11.731560

**Authors:** Oded Mayseless, Joaquin Navajas Acedo, Yinan Wan, Vincent Hahaut, Simone Picelli, Rainer W. Friedrich, Alexander F. Schier

**Affiliations:** Biozentrum, University of Basel, Basel, Switzerland; Institute of Molecular and Clinical Ophthalmology Basel (IOB), Basel, Switzerland; Friedrich Miescher Institute for Biomedical Research, Basel, Switzerland; University of Basel, 4003 Basel

## Abstract

Animals classify odors along behaviorally relevant dimensions, yet how the olfactory bulb, the first stage of olfactory processing, is organized and how this organization relates to odor classification remains unclear. Here, we identify cohesive transcriptional domains within the larval zebrafish olfactory bulb whose differential functional recruitment reflects odor preference. Using a flow-based two-choice assay, we characterized behavioral preferences across chemically diverse odors. Mapping odor-evoked neuronal activity patterns revealed that similarly preferred odors converge onto shared spatial activation patterns in the olfactory bulb. Whole-mount multiplexed spatial transcriptomics revealed that projection neuron and interneuron subtypes form cohesive spatial domains. Mapping odor-activated neurons onto these domains identified preference-biased recruitment of specific neuronal subtypes. Targeted ablation of a TH^+^ interneuron population associated with attraction selectively impaired attraction while sparing aversion. Together, these findings reveal that the olfactory bulb is organized into spatially cohesive, transcriptionally defined domains and that selective recruitment of specific subtypes is linked to odor preference.

**Key findings:**

- Chemically diverse odorants elicit behaviors reflecting olfactory preference.
- Transcriptionally defined neuronal subtypes form cohesive spatial domains in the olfactory bulb.
- Odorants of opposing preference recruit distinct transcriptional domains in the olfactory bulb.
- A TH^+^ interneuron population is selectively required for odor attraction but not aversion.

## Introduction

Olfaction is among the most ancient sensory modalities, enabling animals to detect food, predators, and conspecifics ^1–3^. Across species, animals classify odors along multiple dimensions, including valence, where some odorants are distinguished as attractive or aversive even in the absence of prior experience. Cross-cultural agreement in odor perception ^4^ is consistent with the existence of conserved neural substrates for odor valence. However, how value is assigned to odors remains incompletely understood.

At the anatomical level, the overall organization of the olfactory system, from the periphery to the olfactory bulb (OB), shows striking similarities from insects to vertebrates ^5–11^. In many species, olfactory sensory neurons (OSNs) converge onto discrete glomeruli in the OB, generating a combinatorial sensory map. This map is loosely related to chemical features of odorants, resulting in a partially chemotopic organization of sensory input ^3,12–23^. Within the OB, interactions between projection neurons (PNs) and local interneurons (INs) transform odor-evoked activity patterns and transmit odor information to downstream brain regions ^24–27^. How the cellular and spatial organization of the OB relate to behaviorally relevant odor classification remains unclear.

Chemotopy alone cannot explain how odors drive innate attractive or aversive behaviors. Although certain receptors and glomeruli are preferentially activated by innately attractive or aversive stimuli ^28–35^, most ecologically relevant odorants activate overlapping groups of receptor types rather than isolated channels ^36–38^. Moreover, OB output can diverge substantially from sensory input ^27,39,40^, suggesting that early olfactory circuits reshape odor representations beyond chemical identity. In insects, antennal lobe output has been linked to behaviorally relevant dimensions of odor space, providing evidence that early olfactory circuits can transform sensory input before higher-order processing ^41–44^. In vertebrates, population dynamics have been shown to represent behaviorally relevant dimensions of odor space in higher olfactory areas. For example, in zebrafish, odor-evoked population activity in a subregion of the pallium predicts behavioral preference ^45^. In mammals, the posterolateral cortical amygdala is topographically organized into domains associated with opposing innate behavioral responses, although its odor-evoked activity does not appear to encode valence in a simple global manner ^46–49^. Together, these observations suggest that early olfactory processing may structure odor representations along behaviorally relevant dimensions beyond chemical identity, raising the question of whether such organization is already reflected in the olfactory bulb.

The cellular architecture of the OB provides a potential structural substrate for this reorganization. In vertebrates including zebrafish, the OB displays a conserved concentric laminar organization ^23,50–53^. Distinct subclasses of OSNs converge onto reproducible glomerular territories, partitioning the bulb into stereotyped sensory neuron-defined domains ^23,54^. These domains are overlaid onto an outer layer of glutamatergic PNs, an intermediate layer of GABAergic INs, and a medial proliferative zone at the core ^55–60^. This anatomical organization may therefore provide a structural scaffold for the reorganization of sensory representations, raising the possibility that the spatial distribution of distinct cell types within the OB contributes to this process.

Recent advances in single-cell genomics and spatial transcriptomics have revealed extensive cellular diversity within the vertebrate OB ^61–66^. In zebrafish, initial studies have identified heterogeneity within both PN and IN populations. PNs comprise genetically distinct groups defined by differential transgenic marker expression and by the glomerular territories they innervate ^67^, whereas GABAergic INs can be subdivided into multiple classes based on marker expression and morphology ^26,68^. However, a unified view linking transcriptional identity, spatial position, odor responsiveness, and behavioral output is still lacking. The larval zebrafish OB is well suited to address this gap. Five days postfertilization (dpf), larvae exhibit stereotyped, odor-driven behaviors ^69,70^, including attraction to bile acids and kin odor as well as avoidance of nucleotides and salinity ^71–76^, providing a tractable system in which behavioral output can be linked to circuit organization and function.

Here, we use a systematic, multimodal approach to examine the organizational principles of the larval zebrafish OB, focusing on odor preference as one possible behaviorally relevant axis of organization. By integrating a flow-based two-choice behavioral assay, CaMPARI2-based population activity mapping, single-cell RNA sequencing, and whole-mount multiplexed spatial transcriptomics, we generate an atlas linking the transcriptional identity, spatial position, and functional recruitment of OB neurons. We find that the OB exhibits domain-like organization, with transcriptionally distinct PN and IN subtypes arranged into cohesive domains that are differentially recruited by odors of opposing behavioral preference.

## Results

### A two-choice olfactory assay identifies chemically diverse odors that elicit preference-specific navigation strategies

To identify a comprehensive set of odorants that drive innate attraction or aversion in larval zebrafish, we developed a two-choice, flow-based assay inspired by earlier paradigms in zebrafish and other fish species ^74,77–79^. This approach uses controlled low-flow delivery rather than stagnant water diffusion ^71,73,80^, and more closely mimics the gentle flow (2–12 cm/s) that zebrafish experience in riverbeds ^81–83^. In our configuration (Fig. 1A), two parallel low-flow streams (∼3 mm/s) created adjacent, stable zones containing either odorant or control solution (Fig. S1A). Individual larvae explored this environment freely (Fig. 1A) and exhibited no inherent side bias in the absence of odorant (Fig. 1A; Fig. S1B-C; p = 0.55).

**Figure 1.**
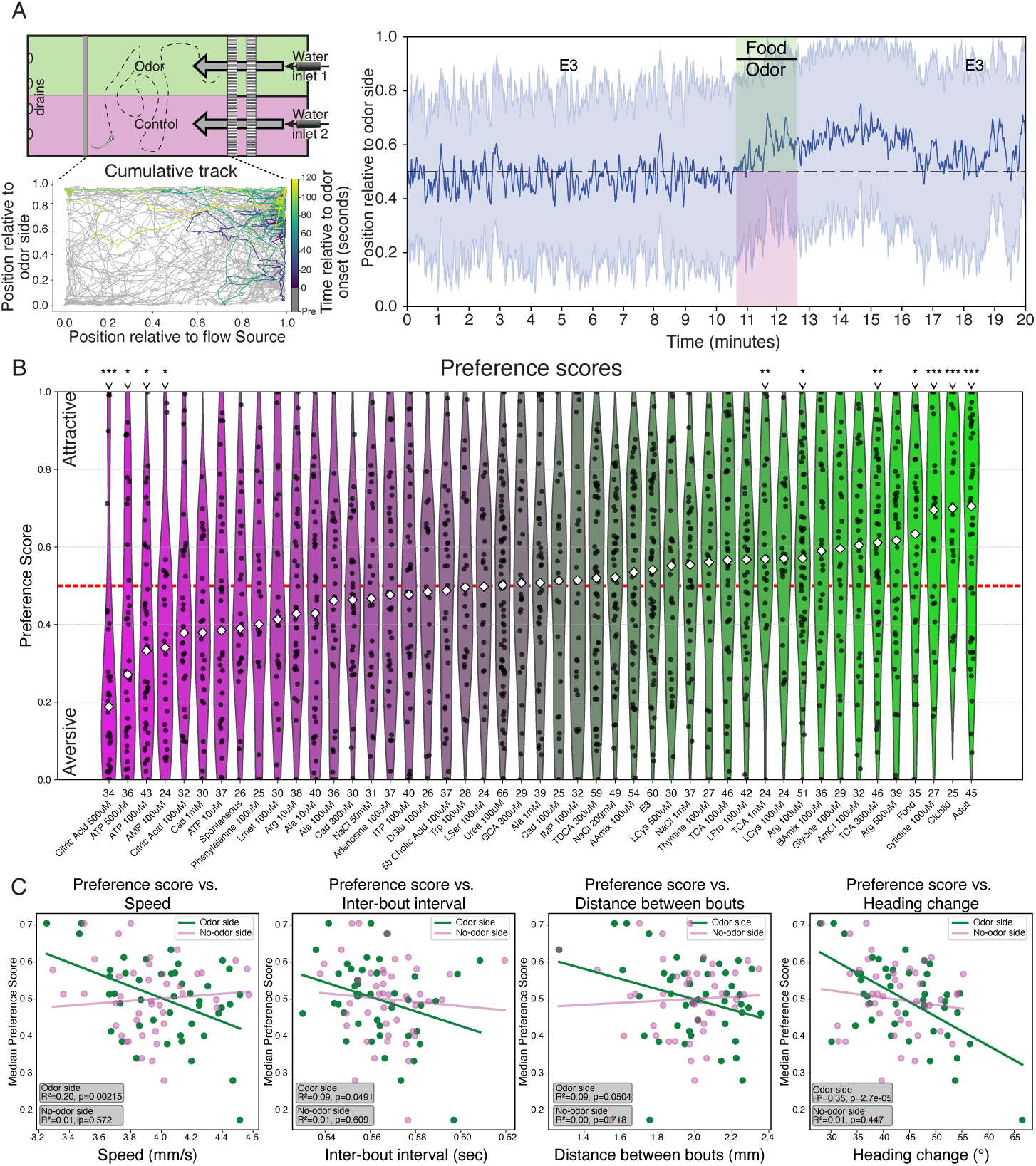
Flow-based two-choice assay reveals odor preferences and preference-associated locomotor patterns. **(A)** Flow-based two-choice assay. **Top left:** schematic of the chamber showing two adjacent laminar streams under constant low flow (∼3 mm/s), with odorant delivered on one side (green) and control solution on the other (magenta). **Bottom left:** cumulative trajectory of a larva exposed to food odor. The pre-odor period is shown in gray, illustrating unbiased exploration, whereas the odor-response period is color-coded with a viridis scale to indicate relative time from odor onset. **Right:** population trace of normalized position relative to the odorant side for food odor, shown as mean ± s.d. Trials consisted of a 10-min habituation period, 2-min odorant delivery, and 8-min washout. No side bias was observed during baseline. **(B)** Odor preference scores across tested odorants. Each point represents one larva. Preference score was defined as the fraction of time spent on the odorant side during odorant delivery. Violins are ordered by median preference score, indicated by white diamonds, and colored from aversive, magenta, to attractive, green. Odorants tested at multiple concentrations are indicated by the concentration shown next to the odorant name. The number of animals per condition is shown below each violin. The red dashed line marks chance-level preference, 0.5. Asterisks indicate significant differences from the E3 control (*p < 0.05, **p < 0.01, ***p < 0.001), assessed using Mann–Whitney U tests or independent t-tests with Bonferroni correction for multiple comparisons. **(C)** Odor preference is associated with odorant-side-specific changes in locomotor strategy. For each locomotor parameter, per-odorant median values were plotted against preference score separately for the odorant side, green, and the control side, magenta. Linear fits are shown. On the odorant side, increasing attraction was associated with reduced swim speed, longer inter-bout intervals, shorter inter-bout distances, and smaller heading changes. These relationships were not observed on the control side, indicating that the locomotor changes were odorant-side-specific rather than reflecting global changes in activity.

Using this setup, we assayed the odor preference of over 1,600 larvae to 46 odorant conditions spanning multiple chemical classes, some at multiple concentrations (Fig. 1B; Supplementary Table 1; Fig. S2). Preference was quantified as the proportion of time spent in the odorant zone during odor delivery. We identified several significantly attractive odorants (e.g., adult zebrafish-conditioned E3, cichlid-conditioned E3, 100 μM cytidine, food extract, 300 μM and 1 mM taurocholic acid (TCA), and 100 μM arginine) and several significantly aversive odorants (e.g., 500 μM citric acid, 500 μM and 100 μM ATP, and 100 μM AMP).

Odors of opposing preference elicited distinct locomotor patterns. On the odor side, attractive odors were associated with slower movement (speed, R^2^ = 0.2, p = 0.002), fewer and smaller swim bouts (inter-bout interval, R^2^ = 0.09, p = 0.05; distance between bouts, R^2^ = 0.09, p = 0.05), and smaller turns (heading change, R^2^ = 0.35, p = 2.7×10^-5^; Fig. 1C). In contrast, aversive odors evoked a distinct pattern, with faster swimming, more frequent turning, and greater distances traveled. On the non-odor side, these relationships disappeared (all metrics R^2^ < 0.01, p > 0.44). Together, these findings indicate that different odor classes promote distinct behaviors, with attractive odors promoting localized exploration and aversive odors driving dispersal and erratic search.

Behavioral responses to odorants depended on both concentration and stimulus timing (Fig. S1D-E). Aversive odors such as ATP, cadaverine, and citric acid evoked progressively stronger avoidance at higher concentrations (Fig. S1D). In contrast, the attractive bile acid TCA exhibited a non-monotonic pattern, with preference tending to increase at moderate concentrations and to decline at the highest concentration. Arginine even shifted preference with dose, switching from mild aversion at low concentrations to mild attraction at higher concentrations (Fig. S1D). Responses also varied with stimulus timing (Fig. S1E). Several responses were strongest after odor offset, with attraction to TCA, food extract, and adult-conditioned E3 peaking during the post-odor period, and aversion to phenylalanine, ITP, IMP, adenosine, and L-cysteine emerging predominantly during odor washout (Fig. S1E). These delayed or secondary phases likely reflect sensitivity to declining gradients or multi-phase odor dynamics, consistent with observations in other species ^84,85^.

Beyond these dynamic features, odor preference was also shaped by chemical structure. Grouping odorants by class (e.g., amino acids, bile acids, nucleotides) explained ∼37% of the variance in preference and exceeded chance expectations under permutation of odor-group labels (p = 0.019; permutation p = 0.047; excluding complex odors - see Supplementary Table 1 for chemical class assignments). Amino acids clustered near neutral and generally elicited mild responses, whereas nucleotides/nucleosides such as ATP and cytidine drove the strongest innate aversion or attraction among single compounds tested.

Together, these results identify salient odors and show that larval zebrafish deploy preference-specific locomotion programs, with attractive odors promoting localized exploration and aversive odors driving dispersal.

### Chemically diverse odorants with similar preferences converge onto shared spatial activity patterns in the olfactory bulb

To ask whether odor preference is reflected in the spatial organization of OB activity, we mapped whole-bulb responses using CaMPARI2, a photoconvertible Ca^2+^ sensor expressed pan-neuronally under the HuC (*elavl3*) promoter (Fig. 2A; Fig. S3A) ^86^. This approach irreversibly labels neurons active during a defined time window, allowing a comprehensive assessment of the spatial patterns of neuronal response to chemically diverse odorants in freely swimming animals.

**Figure 2.**
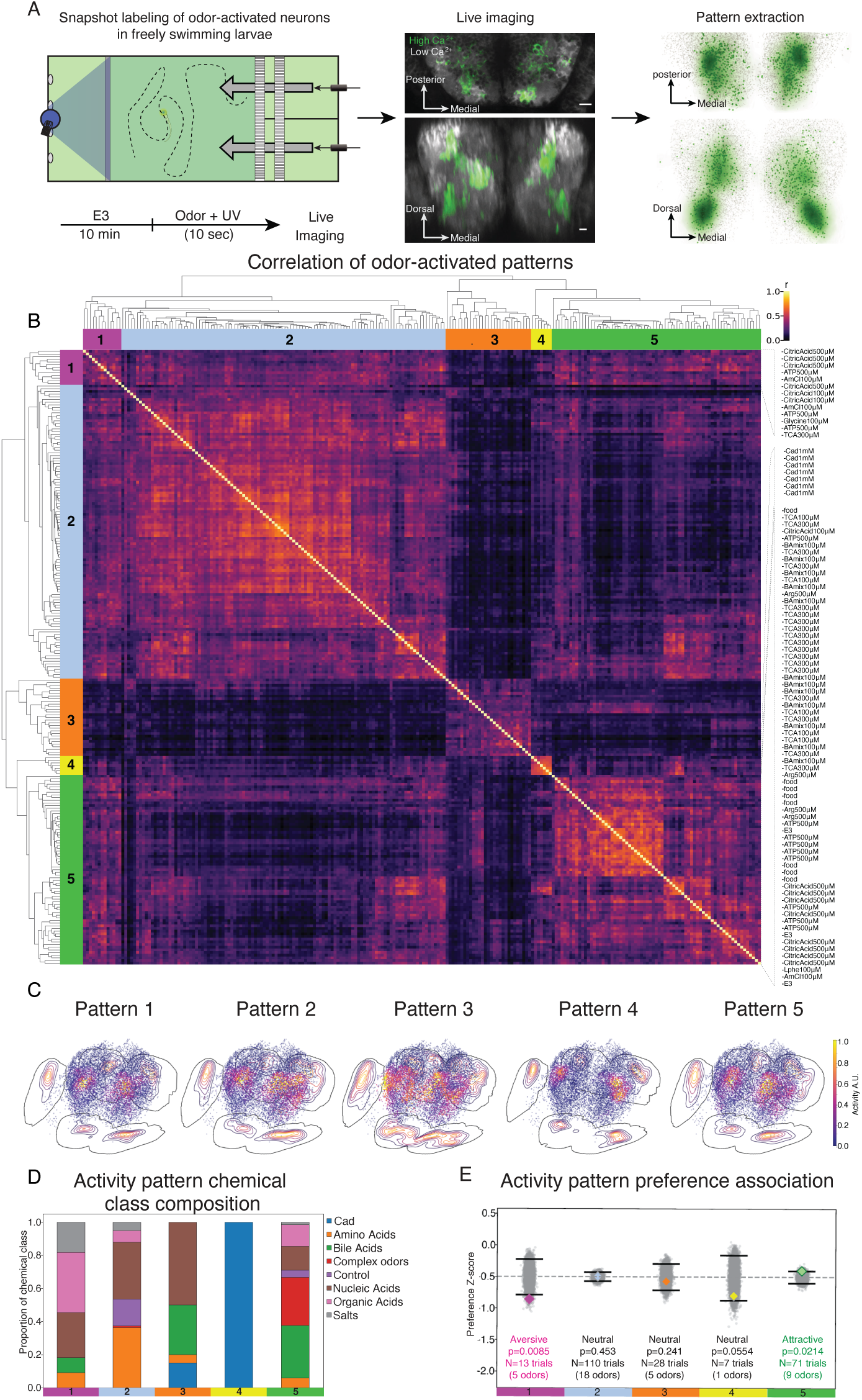
Chemically diverse odorants evoke preference-associated spatial activity patterns in the olfactory bulb. **(A)** CaMPARI2-based snapshot labeling of odorant-activated neurons in freely swimming larvae. Larvae were exposed to an odorant followed by brief UV illumination, irreversibly photoconverting CaMPARI2 in active neurons. Whole-OB volumes were then imaged and registered to a common reference. Center: example substack maximum-intensity projections of native CaMPARI2 signal, gray, and photoconverted CaMPARI2 signal, green, shown in horizontal, top, and coronal, bottom, views. Right: maximum-intensity projections of representative activity-pattern KDEs associated with aversive, magenta, and attractive, green, odorants, shown in horizontal (top) and coronal (bottom) views. **(B)** Pairwise similarity matrix of odorant-evoked activity patterns across 230 larvae exposed to 22 odorants. Each matrix entry shows the Pearson correlation between two 3D OB activity maps. Hierarchical clustering identified five reproducible spatial activity patterns, indicated by color-coded bars. Activity maps from larvae exposed to the same odorant and concentration preferentially co-clustered, indicating reproducible pattern structure across individuals. A complete list of all larval OB activity maps used for this analysis is provided in Supplementary Table 2. **(C)** Three-dimensional visualization of identified odorant-evoked activity patterns using a common anatomical reference cloud. The reference cloud was generated by sampling neurons from all odorant conditions and larvae within ∼30 µm of the olfactory bulb and was reused for each pattern. For each activity pattern, every reference neuron was assigned the value of the corresponding pattern KDE at its spatial coordinate, and dots were colored and opacity-scaled by this normalized KDE value. Thus, the density and intensity of the visualization reflect pattern-specific spatial enrichment, while the underlying anatomical sampling remains identical across patterns. Colored isodensity contours show the corresponding pattern KDE. All views are from a skewed frontal perspective (anterior, lower left; dorsal, upper right). **(D)** Chemical class composition of each spatial activity pattern. Bars show the proportion of odorants from each chemical class assigned to each pattern, with odorants tested at multiple concentrations collapsed by majority-cluster assignment. Some patterns were dominated by a single chemical class, such as cadaverine in Pattern 4, whereas others contained chemically diverse odorants. **(E)** Association between spatial activity patterns and behavioral preference. For each pattern, the mean preference score of member odorants, diamond, was compared with a null distribution generated by shuffling odorant–preference score pairings, gray; 10,000 iterations. Z-scores indicate deviation from chance. Pattern 1 was biased toward aversive odorants (p = 0.0085, FDR q = 0.051), whereas Pattern 5 was biased toward attractive odorants (p = 0.0214, FDR q = 0.064), suggesting that chemically diverse odorants can converge onto shared spatial activity patterns associated with behavioral preference.

To map spatial activity patterns across a chemically diverse odorant panel, we exposed 230 larvae to 22 odorants covering a broad range of preference scores. Larvae were individually placed in the behavior arena, flooded with the odorant, and briefly illuminated with UV light to generate a snapshot of odor response. Forebrain neurons were segmented and activity values extracted at single-cell resolution, yielding robust, odor-specific activity maps (Fig. 2A; Fig. S4; Movie S1).

CaMPARI2 activity patterns recapitulated odor-specific spatial domains previously described by calcium imaging ^19,22,67,87^. For example, bile acids such as TCA activated medio-dorsal OB regions, whereas amino acids such as alanine engaged ventro-lateral territories (Fig. S4). To quantitatively compare responses across odorants, we represented the active OB neurons in each larva as a 3D kernel density estimate (KDE), with each cell weighted by its CaMPARI2 fluorescence intensity. KDE volumes were then compared pairwise across all 230 activity maps using voxel-wise Pearson correlation, yielding an odorant–odorant similarity matrix. Hierarchical clustering of this matrix revealed five partially overlapping spatial activity patterns (Fig. 2B-C; Movie S2; Supplementary Table 2; mean silhouette = 0.11). Larvae exposed to the same odorant at the same concentration tended to cluster together, supporting the robustness of activity maps across individuals. To visualize these activity patterns, we combined the activity maps from larvae assigned to the same cluster while balancing the contribution of each odorant. The resulting pattern-level maps were shown as maximum-intensity projections and as three-dimensional scatter plots, representing the distribution of each pattern (Fig. 2C; Fig. S3D-E; Movie S2).

At the odorant level, chemical class was associated with spatial activity pattern assignment regardless of concentration (Fig. 2D). For instance, pattern 4 was exclusively associated with the amine cadaverine, and amino acids were enriched in pattern 2. Other patterns contained chemically diverse odorants, indicating that chemical similarity alone does not fully account for OB spatial response maps. Activity patterns showed preference-associated biases (Fig. 2E). Averaging preference scores within each pattern revealed biases consistent with behavioral preference: pattern 1 toward aversive odorants (p = 0.0085; FDR q = 0.051) and pattern 5 toward attractive odorants (p = 0.021; FDR q = 0.064). Notably, both patterns 1 and 5 contained odorants from multiple chemical groups (five of eight for pattern 1, seven of eight for pattern 5), demonstrating that chemically diverse odors converge on shared spatial ensembles linked to preference.

The identified activity patterns corresponded to distinct anatomical domains. As expected from its characteristic localized activation, cadaverine responses (pattern 4) were restricted to the dorso-lateral OB ^30^. By contrast, the aversive-associated pattern 1 engaged medial and dorso-lateral regions, while the attractive-associated pattern 5 recruited medial and ventro-lateral regions (Fig. 2C-D; Fig. S3D-E; Movie S2), consistent with a coarse dorsoventral topography associated with odor preference in the OB ^7^.

To test whether preference-associated spatial domains generalize across independent datasets and imaging modalities, we compared CaMPARI2-derived odor maps with an independent brain-wide calcium imaging dataset obtained using two-photon microscopy in larval zebrafish ^88^. In that study, neuronal responses to attractive and aversive odorants were measured across the brain, and regressors associated with odor preference were extracted. Both datasets were registered to the mapzebrain reference space ^89^, enabling voxel-wise comparison within the OB. Comparable odorants shared across datasets, such as food extract and cadaverine, showed moderate to strong spatial correspondence between CaMPARI2 odor maps and the corresponding two-photon activity maps (average Pearson’s R ∼0.42–0.66, Fig. S3B), despite differences in imaging readout. This cross-modal correspondence indicated that, despite distinct readouts, the two datasets captured overlapping OB activity structure suitable for comparing preference-associated spatial domains.

We next asked whether preference-associated spatial structure in the OB was consistent across datasets. For each odorant, we correlated its CaMPARI2 activity map separately with the positive- and negative-preference maps defined in the two-photon dataset. We then quantified this spatial preference bias as ΔR, defined as the correlation with the two-photon positive-preference map minus the correlation with the two-photon negative-preference map. Positive ΔR values therefore indicate greater similarity to independently defined positive-preference domains, whereas negative ΔR values indicate greater similarity to negative-preference domains. Across odorants, ΔR correlated with behavioral preference measured in our two-choice assay (Fig. S3C; Spearman ρ = 0.46, p < 0.05), suggesting that preference-associated spatial organization in the OB is partially reproducible across independent datasets and imaging modalities.

Spatial activity in the larval OB therefore reflects both odorant chemistry and behavioral preference. The convergence of chemically diverse odorants with similar preferences onto shared, anatomically structured patterns, suggests that OB population activity is organized in part by behaviorally relevant odor features.

### Single-cell and spatial transcriptomics resolve transcriptionally distinct olfactory bulb subtypes

To connect odor-evoked activity patterns to neuronal subtypes within the larval OB, we generated a single-cell RNA sequencing (scRNA-seq) reference atlas from GFP-enriched telencephalic cells isolated from *-8kb:cldnb:lynEGFP* transgenic larvae ^90,91^ (Fig. 3A; Fig. S5A). We filtered this dataset for postmitotic neurons using canonical neuronal, maturation, and progenitor markers, yielding >2,000 high-quality cells that segregated into 26 distinct transcriptional clusters (Fig. 3B–D’; Fig. S5A,B). Cluster identities could be predicted with high accuracy (∼75%) using only combinatorial transcription factor expression, a hallmark of neuronal subtype identity ^92,93^ (Fig. S5C). The most informative transcription factors included markers characteristic of projection neurons (PNs), such as *eomesa, tbr1a, tbr1b* ^94^ and interneurons (INs), such as *dlx5a, dlx2a* ^56,95,96^. By contrast, a classifier trained exclusively on ribosomal genes (Fig. S5D) performed substantially worse (∼23% accuracy), underscoring the specificity of transcription factor-based classification ^97^.

**Figure 3.**
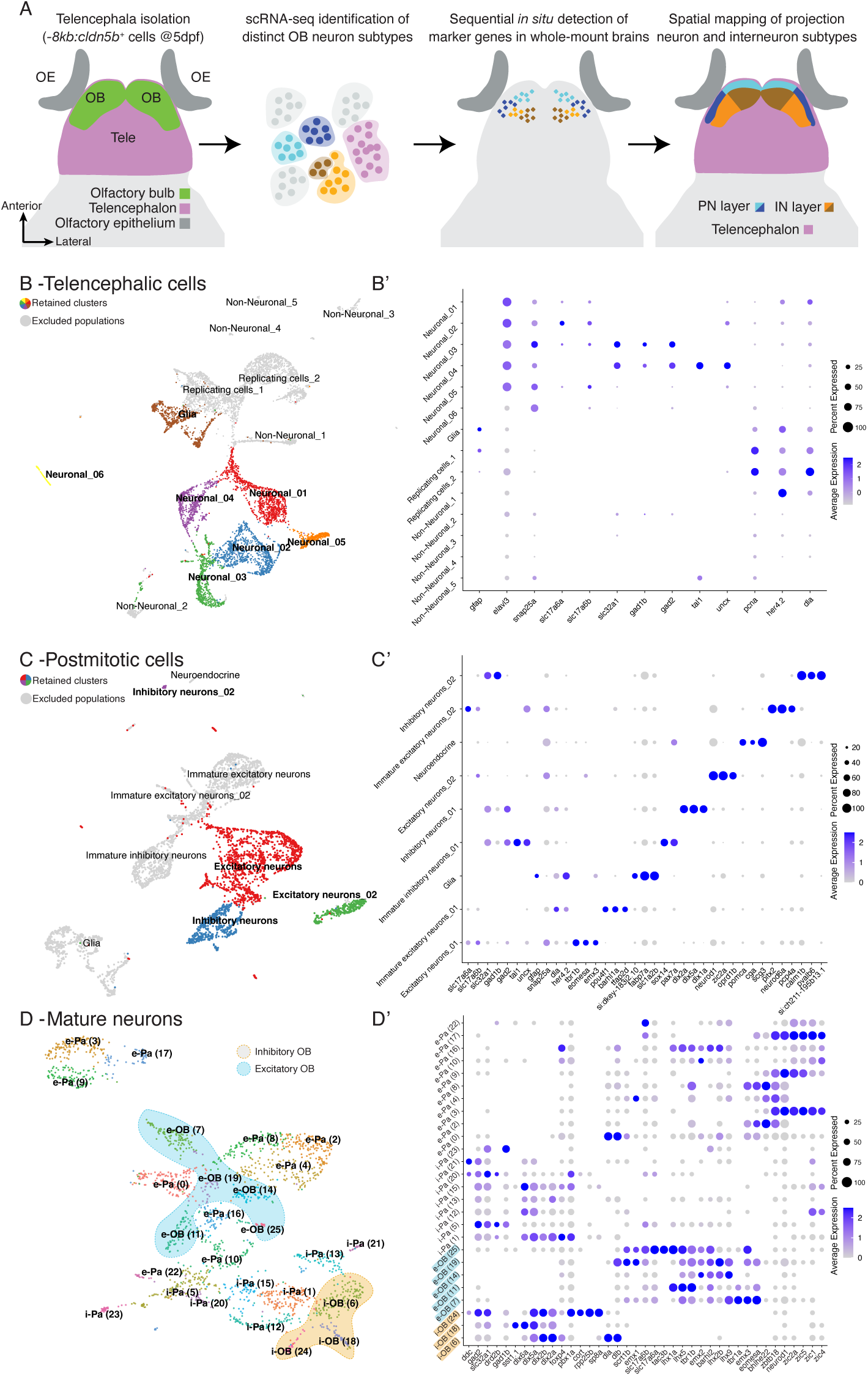
Single-cell transcriptomics identifies transcriptionally distinct neuronal subtypes in the zebrafish telencephalon. **(A)** Experimental workflow for linking telencephalic neuronal subtypes to spatial domains. Larval telencephala, 5 dpf, were dissected, dissociated, and profiled by scRNA-seq. Marker genes were then mapped in whole-mount brains by sequential *in situ* hybridization, seqFISH, to relate transcriptional subtypes to their anatomical distributions. **(B)** UMAP of telencephalic cells after initial quality control. Colored clusters indicate postmitotic populations retained for downstream analysis, whereas gray clusters indicate non-neuronal, proliferative, or immature populations that were excluded. Each dot represents one cell. **(B′)** Dot plot of broad neuronal, non-neuronal, proliferative, and immature markers used to define the postmitotic neuronal subset. Dot size indicates the fraction of cells expressing each gene; color indicates scaled average expression. **(C)** UMAP of the postmitotic neuronal subset. Colored clusters represent mature neuronal populations retained for downstream analysis, whereas excluded populations are shown in gray. **(C′)** Dot plot of excitatory and inhibitory markers used to assign broad neuronal identities, together with the top three variable genes per cluster. Dot size indicates the fraction of cells expressing each gene; color indicates scaled average expression. **(D)** UMAP of mature neurons enriched for olfactory bulb identities. Subtypes are annotated as excitatory, “e”, or inhibitory, “i”, and classified as pallial, “Pa”, or olfactory bulb-associated, “OB”, based on marker-gene expression and imputed spatial distributions, see Fig. 4J–K and Movie S4. Shaded outlines mark spatially validated OB subtypes, with excitatory projection neuron, PN, subtypes in blue and inhibitory interneuron, IN, subtypes in orange. **(D′)** Dot plot of subtype-defining marker genes. Dot size indicates the fraction of cells expressing each gene; color indicates scaled average expression.

To annotate neuronal subtypes as OB-associated, we examined the expression pattern of the canonical transcription factors highlighted by a random forest classifier. We found that many of these markers are shared between bulb-associated and pallial spatial domains. For example, the conserved PN markers (*tbr1b, eomesa, slc17a6a*) and IN markers (*gad1b, dlx5a, dlx2a, slc32a1*) are co-expressed across multiple subtypes and regions (Fig. 3D’; Fig. S5E; mapzebrain ^89^). This observation echoes recent reports that transcriptionally similar neuronal types can be distributed in distinct topographic locations ^98^ and with prior anatomical descriptions of the dorsal subpallium ^99^.

To resolve the anatomical context of transcriptionally defined subtypes, we first asked how many genes are required to reliably classify these subtypes within the full single-cell dataset. Random forest classifiers trained on subtype labels showed that reliable subtype assignment required a combinatorial set of 32 informative genes (Fig. S5F), motivating a multiplexed spatial mapping strategy. We therefore adapted the weMERFISH approach ^100^ into a sequential hybridization workflow (seqFISH) for whole-mount imaging of intact larval brains. A curated 68-gene probe panel, comprising highly variable genes, transcription factors, and known markers (Supplementary Table 3; ^61,93,101–104)^, was sequentially imaged at single-molecule resolution in two brains (Fig. 4A-D; Figs. S6-7; Movie S3). Reproducibility between brains was high (log-log correlation R = 0.93; Fig. 4E), demonstrating reliable gene detection across samples.

**Figure 4.**
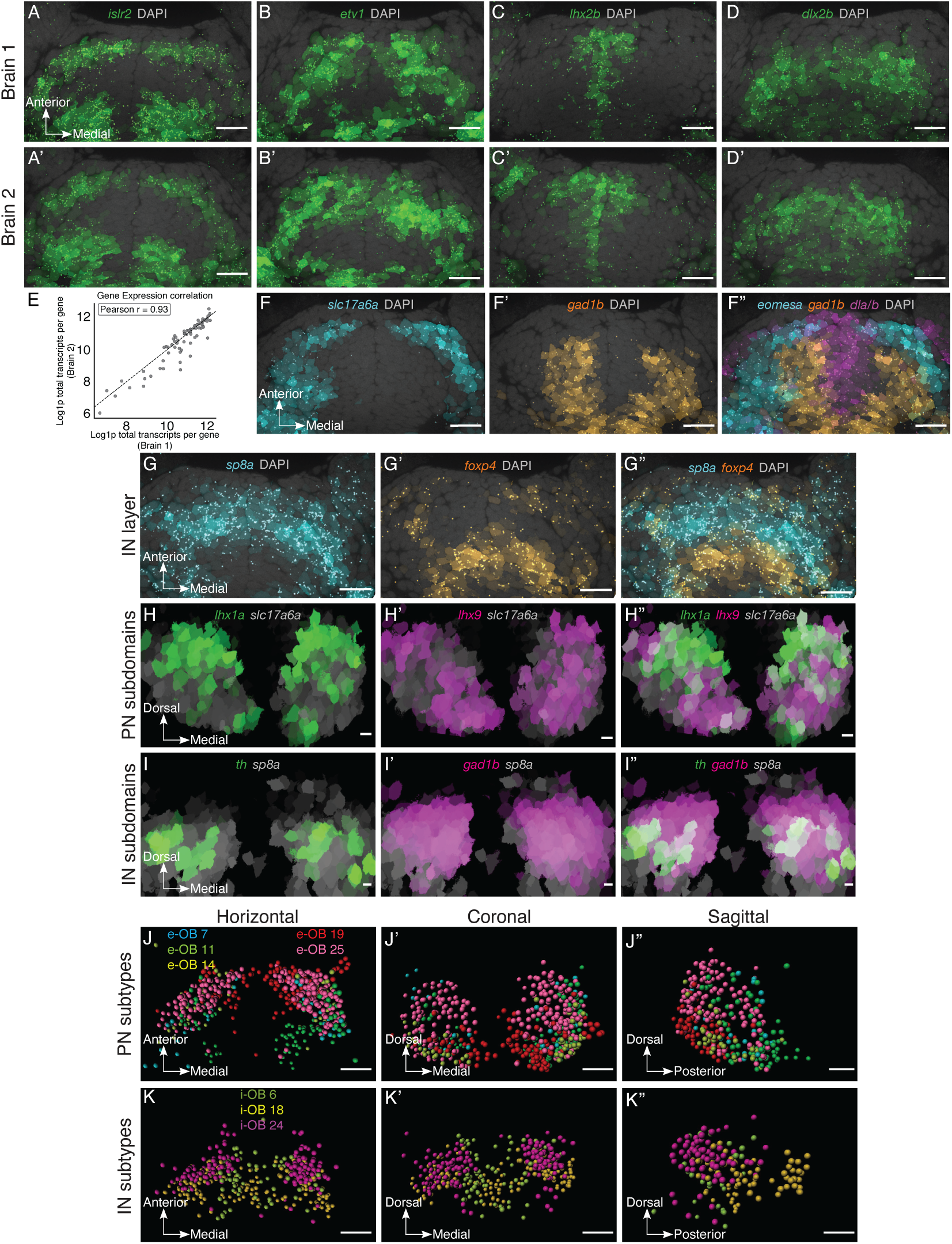
Whole-mount spatial transcriptomics resolves the spatial organization of olfactory bulb neuronal subtypes. **(A–D)** Horizontal maximum-intensity projections of five optical sections from two brains, rows, show expression patterns of selected genes, columns: *islr2* (A,A′), *etv1* (B,B′), *lhx2b* (C,C′), and *dlx2b* (D,D′). DAPI is shown in gray. Colored dots indicate detected transcripts, and cell-mask intensity reflects normalized per-cell gene counts. **(E)** Between-brain reproducibility of spatial transcript detection. Log–log scatter plot shows the total number of detected transcripts per gene across the two brains. **(F–F′′)** Canonical concentric organization of the olfactory bulb. Horizontal maximum-intensity projections of five optical sections show *slc17a6a* marking the projection neuron, PN, layer (cyan; F), *gad1b* marking the interneuron, IN, layer (orange; F′), and *dla/b* marking the mitotic zone (magenta; F′′), with DAPI shown in gray. Display conventions are as in A–D. **(G–G′′)** Anatomical and transcriptional definition of the posterior OB boundary. Horizontal maximum-intensity projections of five optical sections show *sp8a* (cyan; G), *foxp4* (orange; G′), and the merged image with DAPI in gray (G′′). *sp8a* marks an anterior IN territory, whereas *foxp4* delineates the posterior pallial boundary. Display conventions are as in A–D. **(H–H′′)** Spatial organization of PN marker expression. Coronal maximum-intensity projections show *slc17a6a* in gray together with *lhx1a* (green; H) and *lhx9* (magenta; H′,H′′), revealing complementary PN territories. **(I–I′′)** Spatial organization of IN marker expression. Coronal maximum-intensity projections show *sp8a* in gray together with *th* (green; I) and *gad1b* (magenta; I′,I′′), revealing distinct medial and anterior IN territories. **(J–K′′)** Tangram-based imputed subtype distributions within the OB. Imaris-based renderings are shown in horizontal (J,K), coronal (J′,K′), and sagittal (J′′,K′′) orientations. **(J–J′′)** Distribution of PN subtypes: e-OB(7), e-OB(11), e-OB(14), e-OB(19), and e-OB(25). **(K–K′′)** Distribution of IN subtypes: i-OB(6), i-OB(18), and i-OB(24). Colors indicate subtype identity. Scale bars, 10 µm. Axis arrows indicate anatomical orientation.

Consistent with earlier anatomical observations ^23,51–53,56,105,106^, our larval OB atlas revealed a concentric organization, with an outer PN layer (*slc17a6a*), an intermediate IN layer (*gad1b*), and a medial proliferative zone (*dla/b*) (Fig. 4F). Visualizing multiple genes within the same brain resolved finer subdivisions within these layers (Fig. 4G-I). In the *slc17a6a^+^* PN layer, *lhx9* and *lhx1a* partition the OB into complementary domains with *lhx1a* biased medio-dorsally and *lhx9* ventral-laterally. Within the *gad1b^+^*IN layer, tyrosine hydroxylase 1 (*th1*; hereafter referred to as *th*) delineated a distinct medial subdomain. Importantly, *sp8a* defined an anterior strip of the IN layer, in accordance with its broad role as an IN marker in the mammalian OB ^107,108^, whereas *foxp4* provided a sharp posterior landmark at the OB boundary in accordance with its role in mammalian cortical development ^109,110^.

To infer the spatial distribution of transcriptionally defined OB subtypes, we aligned our single-cell RNA-seq dataset to the spatial transcriptomic data using Tangram ^111^, a machine learning–based approach for integrating single-cell and spatial transcriptomics data. Using the 68 measured gene expression patterns as anchors, Tangram projected the single-cell transcriptional atlas into anatomical space (Movie S4). Tangram-based spatial predictions were supported by leave-one-gene-out validation, concordance with independently measured expression patterns, and reproducible subtype composition across brains (r = 0.94; Fig. S8A-E; Movie S4; Movie S5). We then annotated OB-associated subtypes by evaluating the predicted spatial distribution of transcriptional subtypes relative to *foxp4* expression, which provided a reproducible posterior landmark (Fig. 4G–G’’), the symmetric distribution within the OB, and the concentric laminar organization of the bulb to distinguish OB-associated from pallial-associated subtypes.

Using these criteria, five PN subtypes (*slc17a6a*^+^) (e-OB(7), e-OB(11), e-OB(14), e-OB(19), e-OB(25)) and three IN subtypes (*gad1b^+^)* (i-OB(6), i-OB(18), i-OB(24)) localized predominantly to the OB, whereas other subtypes, such as subtype e-Pa(16), mapped to pallial regions (Fig. 4J-K; Movie S4). Notably, i-OB(24), a GABAergic IN subtype, co-expressing *sp8a*, *pbx1a*, *gad1b*, and *slc32a1* (Fig. 3D’), was confined to the OB territory and expressed markers associated with short-axon INs in the mammalian OB ^112–114^, reinforcing the anatomical–transcriptional boundary.

Together, scRNA-seq defined the transcriptional landscape of larval OB neurons, and multiplexed spatial transcriptomics placed these signatures in anatomical context. Spatial mapping confirmed canonical OB lamination and enabled boundary-informed assignment of transcriptionally defined subtypes, resolving five PN and three IN subtypes that localized predominantly to the OB. This spatial subtype map provides a framework for linking transcriptional identity to odor-evoked activity patterns.

### Olfactory bulb neuronal subtypes exhibit cohesive spatial domains

Having identified OB-associated neuronal subtypes, we asked whether they occupy distinct spatial domains or are distributed across the bulb. To address this question, we quantified the spatial cohesion of OB subtypes using a nearest-neighbor–based metric (Fig. S8F–H), defined for each cell as the fraction of nearby cells belonging to the same transcriptional subtype. Higher values indicate stronger local clustering of subtype-matched cells. The relative ranking of subtype cohesion was highly reproducible across datasets (Fig. S8G; Spearman ρ = 0.91). In both brains, individual OB-assigned cells had significantly higher cohesion scores than non-OB cells (Fig. S8H; brain 1: 0.267 vs. 0.200, p = 3.1×10^-18^; brain 2: 0.300 vs. 0.233, p = 3.7×10^-25^), consistent with the OB being organized into spatially segregated transcriptional domains. The same trend held at the subtype level, where OB subtypes had higher median cohesion scores than non-OB subtypes (Fig. S8G-H), supporting a spatial organization into cohesive transcriptional domains.

To further characterize how subtype domains relate to one another in space, we estimated their three-dimensional extents using volumetric kernel density estimation and quantified pairwise territorial overlap (Fig. S8I). Consistent with the laminar organization of the OB, PN and IN domains were largely segregated, sharing only ∼5% of their spatial extent. Within each class, however, territorial overlap differed: IN subtypes showed moderate reciprocal overlap (median Dice = 0.14–0.21), consistent with partially shared or nested spatial niches, whereas PN subtypes were more mutually exclusive (median Dice = 0.04–0.10), consistent with a tiling-like organization.

Together, these analyses show that transcriptionally defined neuronal subtypes are not arranged in a fine-grained mosaic but instead occupy structured, cohesive domains, revealing a domain-like transcriptional organization of the OB.

### Distinct olfactory bulb transcriptional subtypes show preference-biased recruitment

To ask how preference-associated activity patterns relate to transcriptional subtype organization, we compared CaMPARI2-derived activity maps with the spatial transcriptomic atlas of the OB. This comparison suggested coarse spatial correspondence, with activity patterns associated with odors of opposing preference occupying partially overlapping medial regions of the OB while appearing to differ in their finer-scale spatial distributions (Fig. 2C; Fig. S3D-E; Movie S2). Within the IN domains, the attractive-associated pattern was shifted medially, corresponding to regions associated with i-OB(24), whereas the aversive-associated pattern extended more laterally, corresponding to i-OB(18). Within PN domains, the attractive-associated pattern recruited more ventrolateral regions, whereas the aversive-associated pattern engaged more dorsolateral regions consistent with recruitment of distinct PN subtypes.

To assess the relationship between CaMPARI2-activity profiles and transcriptional identity, we used CaMPARI2-seq. CaMPARI2-based photoconversion was coupled to FACS isolation of active neurons and FLASH-seq profiling, a protocol optimized for low cell numbers ^115^ (Fig. 5A; Fig. S9A-B). Larvae were photoconverted under four conditions chosen to partially dissociate chemical class from behavioral preference: the attractive bile acid taurocholic acid (TCA, 300 μM), the attractive nucleoside cytidine (Cyt, 100 μM), the aversive nucleotide adenosine triphosphate (ATP, 500 μM), or an E3 vehicle control. This design enabled a direct comparison of neuronal populations engaged by attractive versus aversive odorants, independent of chemical class.

**Figure 5.**
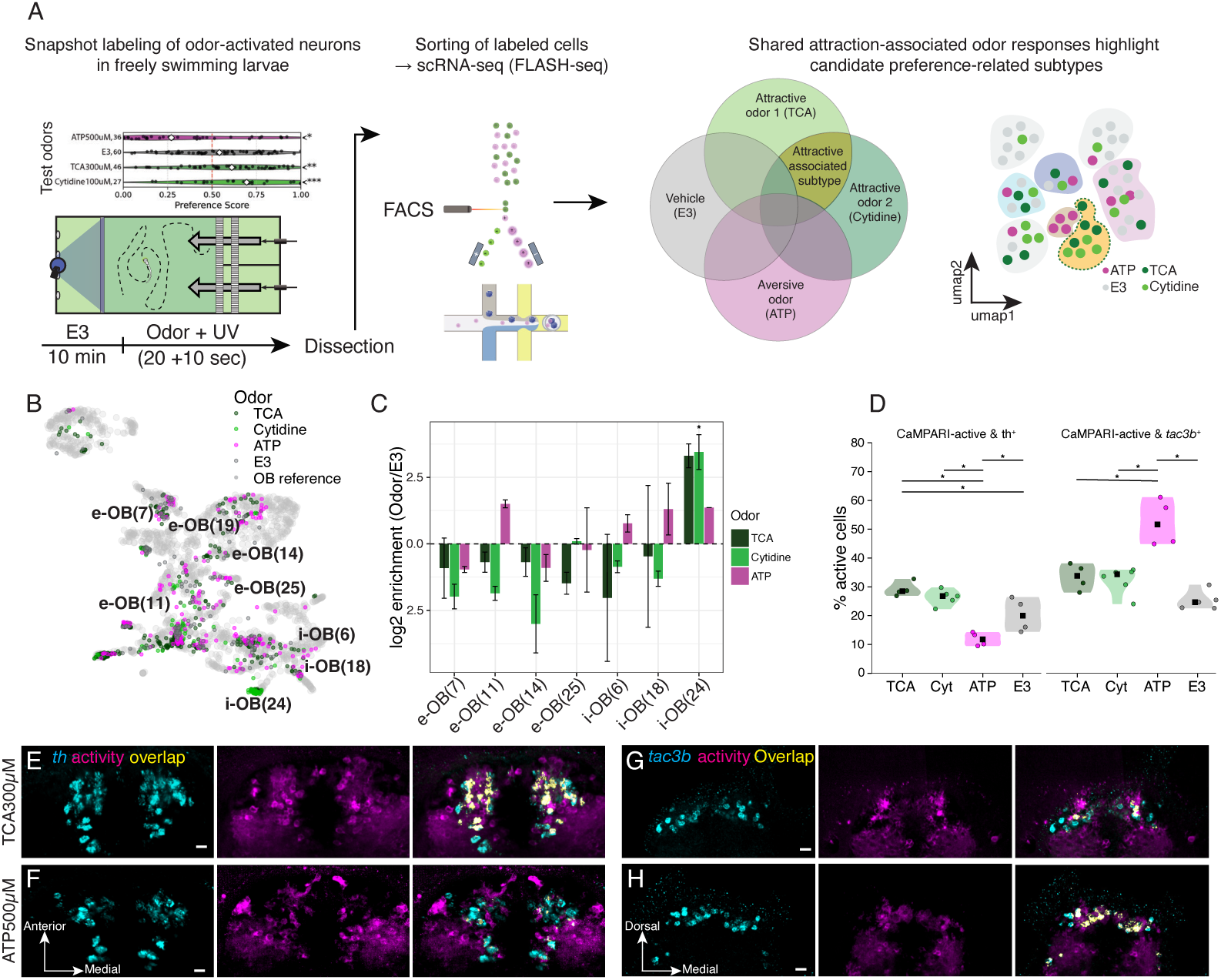
Distinct olfactory bulb transcriptional subtypes are differentially recruited by odors of opposing preference. **(A)** Experimental workflow for CaMPARI2-seq. Freely swimming larvae were exposed to one of four conditions: the attractive bile acid TCA, 300 μM; the attractive nucleoside cytidine, 100 μM; the aversive nucleotide ATP, 500 μM; or E3 vehicle control. These odorants were selected to partially dissociate chemical class from behavioral preference, with preference scores shown at top left. During odorant exposure, larvae were photoconverted to permanently label active neurons. Telencephala were then rapidly dissected, CaMPARI2-photoconverted neurons were isolated by FACS, and single-cell transcriptomes were profiled by FLASH-seq. Right: schematic UMAP illustrating CaMPARI2-seq cells, colored by odorant condition, and their mapping onto the single-cell OB atlas. **(B)** CaMPARI2-seq cells projected onto the OB reference atlas and colored by condition. Labeled clusters indicate inhibitory interneuron, i-OB, and excitatory projection neuron, e-OB, subtypes. Attractive odorants showed the strongest enrichment in i-OB(24), whereas ATP showed enrichment in i-OB(6) and i-OB(18), with additional ATP-associated bias in projection neuron subtypes. **(C)** Transcriptional subtype enrichment following odorant stimulation. Bars show log2 enrichment relative to E3 controls within replicate, calculated from the proportion of mapped CaMPARI2-seq cells assigned to each subtype. Error bars indicate bootstrap 5–95% confidence intervals. Attractive odorants, TCA in dark green and cytidine in light green, showed preferential enrichment in i-OB(24), a subtype not recovered from E3 controls in the sampled dataset. ATP, magenta, showed enrichment in i-OB(6) and i-OB(18), with an additional bias toward e-OB(11). **(D)** Independent validation of subtype recruitment by combined CaMPARI2 labeling and HCR. Violin plots show the fraction of HCR-positive neurons with elevated CaMPARI2 activity, measured as R/G ratio, for *th*^+^ (left) and *tac3b*^+^ (right) populations across conditions. Attractive odorants showed elevated recruitment of *th*^+^ neurons relative to ATP and E3 controls, whereas ATP preferentially recruited *tac3b*^+^ neurons compared with TCA, cytidine, and E3. Points represent individual brains; black squares indicate medians; horizontal bars indicate significant pairwise comparisons. Statistical comparisons were performed using a Kruskal–Wallis test followed by two-sided Mann–Whitney U tests with Benjamini–Hochberg FDR correction. **(E–H)** Representative images showing *th*^+^ (E,F) or *tac3b*^+^ (G,H) HCR signal, cyan, CaMPARI2 activity, magenta, and co-labeled cells, yellow, following exposure to TCA, 300 μM (E,G), or ATP, 500 μM (F,H). Horizontal projections are shown in E–F; coronal projections are shown in G–H. Scale bars, 20 μm.

Mapping CaMPARI2-seq profiles onto the single-cell atlas (Fig. 3D) revealed odor-specific biases in OB subtype recruitment (Fig. 5B–C; Fig. S9C–E). Relative to E3 controls, both attractive odors showed positive enrichment in the short-axon-like IN subtype i-OB(24), whereas the aversive odor ATP was associated with the IN subtypes i-OB(6) and i-OB(18), and the PN subtype e-OB(11). To assess whether the observed preference-associated recruitment pattern exceeded chance, we performed a permutation test in which odor labels were randomly shuffled across cells within each replicate, and computed an empirical null distribution for the attractive-versus-aversive preference contrast per subtype. This confirmed significant aversive-biased recruitment of i-OB(6), i-OB(18), and e-OB(11), and significant attractive-biased recruitment of i-OB(24) (positive-direction empirical q = 0.002), with all three IN subtypes reaching significance compared to only one PN subtype (Fig. S9E). Direct pairwise odor comparisons were consistent with these findings for i-OB(6), i-OB(18), and i-OB(24), including stronger i-OB(24) recruitment by TCA than ATP (mean log2OR = +2.64, q = 0.034), whereas e-OB(11) was significant in the preference-level contrast but not in pairwise comparisons (Fig. S9D-E; empirical q = 0.0085; pairwise q ≥ 0.091). Because some subtypes were represented by few recovered cells, statistical inference for these populations remained limited.

To orthogonally examine OB subtype associations at cellular resolution, we combined CaMPARI2 labeling with HCR fluorescence *in situ* hybridization for markers selectively expressed by candidate subtypes. Overlaying *th* expression, a selective marker for subtype i-OB(24), with CaMPARI2 activity revealed condition-dependent recruitment of *th*^+^ neurons. Activation was higher in TCA- and cytidine-exposed larvae than in ATP-exposed larvae, with intermediate recruitment in E3 controls (TCA, 29.1 ± 1.3%; Cyt, 26.3 ± 1.2%; ATP, 11.8 ± 1.2%; E3, 20.2 ± 2.9%; Fig. 5D–F). For PNs, we focused on a subtype with a readily HCR-compatible selective marker, examining e-OB(25) via *tac3b*, which also showed condition-dependent recruitment, with the highest activation observed in ATP-exposed larvae (ATP, 52.4 ± 4.1%; TCA, 33.6 ± 2.3%; Cyt, 32.5 ± 1.8%; E3, 25.3 ± 1.5%; Fig. 5D,G,H).

Together, these results identify candidate OB subtypes whose recruitment is associated with odor preference. The attractive odorants tested preferentially recruited a dopaminergic short-axon-like interneuron subtype corresponding to i-OB(24), whereas the aversive odorant ATP preferentially recruited i-OB(6) and i-OB(18), with additional biases in PN populations. Thus, odors of opposing preference recruit distinct olfactory bulb transcriptional subtypes.

### Targeted ablation of TH^+^ interneurons selectively impairs attraction while sparing aversion

To test whether dopaminergic short-axon-like INs are required for attractive odor responses, we selectively ablated TH^+^ OB neurons labeled by a GFP reporter driven from the endogenous tyrosine hydroxylase locus (th:QF2>GFP) ^116^ using targeted two-photon irradiation. To control for potential effects of anesthesia, mounting, and laser exposure independent of cell loss, sham-treated larvae underwent identical preparation and targeting procedures but were irradiated at subthreshold laser power insufficient to induce photodamage or GFP loss (Fig. 6A). Post hoc imaging confirmed a ∼50% reduction in GFP^+^ somata in two-photon-ablated larvae relative to sham controls (Fig. 6B), with the glomerular space largely devoid of GFP^+^ innervation, but without gross anatomical disruption or apparent loss of GFP^−^ cells. Baseline locomotion was indistinguishable between two-photon-ablated and sham larvae (Fig. S9F; mean speed, p = 0.199; total distance traveled, p = 0.169), and pilot experiments confirmed that repeated arena exposure did not alter locomotion parameters or odor preference (Fig. S9G-H), confirming that ablation did not impair general locomotor capacity.

**Figure 6.**
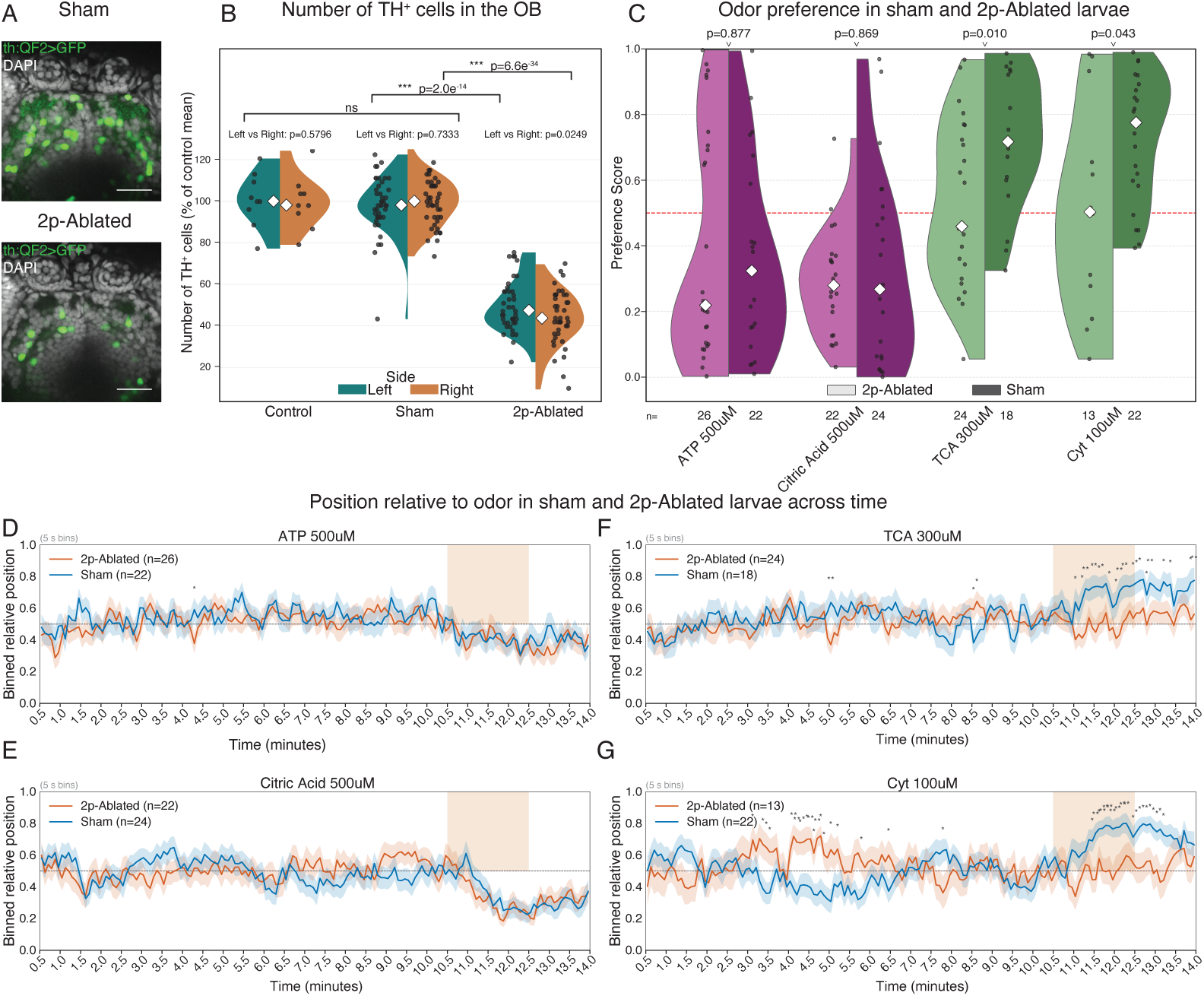
Targeted ablation of TH^+^ olfactory bulb interneurons selectively impairs attraction while sparing aversion. **(A)** Representative confocal single optical sections of the OB in sham-treated and two-photon-ablated larvae at 5 dpf, one day post-irradiation. TH^+^ neurons are visualized by GFP expression from the endogenous *th* locus, green; DAPI, gray, shows nuclei and anatomical context. Sham larvae underwent the same mounting and targeting procedures but were irradiated at subthreshold laser power that did not induce cell loss. **(B)** Quantification of TH^+^ neuron numbers in the OB following sham treatment, two-photon ablation or untreated control larvae are for reference. Violin plots show GFP^+^ cell counts per brain; points represent individual brains and white diamonds indicate medians. Two-photon ablation reduced TH^+^ cell number by ∼50% relative to sham controls. A small left–right difference was detected in the ablated group, but both hemispheres showed robust reduction in glomerular GFP expression. **(C)** Odor preference scores for sham-treated and two-photon-ablated larvae across attractive odorants, TCA, 300 μM, and cytidine, 100 μM, and aversive odorants, ATP, 500 μM, and citric acid, 500 μM. Each point represents one larva; white diamonds indicate medians. Ablation of GFP^+^ TH^+^ neurons significantly reduced attraction to TCA and cytidine, whereas avoidance of aversive odorants was unaffected. The dashed red line indicates chance-level preference, 0.5. Group sizes and *p*-values are indicated below and above each violin, respectively. **(D–G)** Time-resolved mean position relative to the odorant source for sham-treated, gray, and two-photon-ablated, colored, larvae, binned in 5-s intervals and shown as mean ± SEM. Shaded regions indicate the odorant-delivery epoch. Asterisks mark time bins with significant divergence between sham and ablated groups. Group differences were most apparent during exposure to attractive odorants, TCA, 300 μM, and cytidine, 100 μM, whereas traces for aversive odorants, ATP, 500 μM, and citric acid, 500 μM, remained largely overlapping.

Following recovery, larvae were assayed for odor-guided behavior using multiple attractive and aversive odorants per larva. Ablation of GFP^+^ neurons significantly reduced preference for the attractive odors (Fig. 6C), TCA (p = 0.010; n = 24 ablated, 18 sham) and cytidine (p = 0.043; n = 13 ablated, 22 sham). In contrast, avoidance of aversive odors remained intact (ATP, p = 0.877; n = 26 ablated, 22 sham; citric acid, p = 0.869; n = 22 ablated, 24 sham), indicating that aversive behavioral responses, as well as general sensory detection and motor execution, were preserved.

Time-resolved analysis further supported this dissociation. Binned preference traces revealed that divergence between sham and two-photon-ablated larvae emerged selectively during the odor-exposure epoch for attractive odors, while traces for aversive odors remained largely overlapping (Fig. 6D–G).

Together, these results support a selective requirement for TH^+^ interneurons in attraction to chemically diverse odors, while aversive responses and general locomotor capacity remain intact. This selective impairment suggests that attraction and aversion engage at least partially separable circuit elements within the OB, rather than relying on a fully distributed or redundant representation.

## Discussion

Olfactory sensory input is processed in the olfactory bulb (OB), where it is transformed before being relayed to downstream brain regions. How cellular and spatial OB organization relates to behaviorally relevant odor classification has remained incompletely understood. Using a systematic, multimodal approach, we find that the larval zebrafish OB is organized into spatially cohesive, transcriptionally defined neuronal domains that are differentially recruited by chemically diverse odors of opposing preference. Targeted ablation of a dopaminergic short-axon-like interneuron population selectively disrupted attraction while sparing aversion, supporting a subtype-dependent contribution of OB interneurons to preference-biased odor processing.

### The zebrafish olfactory bulb is organized into spatially cohesive transcriptional domains

The combination of scRNA-seq and whole-mount spatial transcriptomics enabled us to define five PN subtypes and three IN subtypes in the OB. Since many individual transcriptional markers are shared across OB and pallial regions, systematic high-confidence subtype assignment required co-detection of at least 32 genes within a single individual sample (Fig. S5F-G) via multiplexed whole-mount spatial transcriptomics (Figs. S6-7, Movie S3). By examining their distribution relative to a reproducible posterior pallial landmark (Fig. 4G), we were able to assign transcriptionally defined subtypes with anatomical precision (Fig. 4J-K). For example, e-Pa(16), which is transcriptionally similar to excitatory projection neurons, mapped to the pallium, while i-OB(24), which resembles subpallial interneuron subtypes, localized specifically to the OB. Thus our boundary-informed approach revealed that several subtypes previously ambiguous in transcriptional space localize either to the OB or to adjacent pallial regions, underscoring the importance of spatial context for cell-type annotation in regions with shared transcriptional programs.

Mapping multiple genes in space and inferring subtype distributions afforded additional insight into the cellular organization of the bulb and how it may relate to odor processing. We found that transcriptionally defined subtypes were not uniformly distributed across the OB, but instead occupied cohesive, largely non-overlapping domains. For example, e-OB(25) consistently occupied a medial-dorsal PN domain (Fig. 4J), whereas i-OB(24) and i-OB(18) occupied largely medial and lateral domains of the interneuron layer, respectively (Fig. 4K).

This organization is notable because olfactory input is itself spatially ordered. Odorant chemistry is represented chemotopically in the OB ^19^, and recent spatial transcriptomic studies further show that receptor expression in the epithelium is aligned with stereotyped glomerular projections to the OB ^65,66^. Yet OB output diverges from sensory input patterns ^27,40^, and downstream regions represent behaviorally relevant dimensions of odor space ^45,46^. These observations raise the question how the cellular organization within the bulb could support the transformation of structured sensory input. One possibility is a uniform “tiled mosaic,” in which transcriptionally distinct neuronal subtypes are broadly distributed across the bulb, providing each glomerular region with a similar complement of cell types. Such an organization could be advantageous to extract a common set of physico-chemical features in each glomerular channel. Alternatively, projection neuron and interneuron subtypes could occupy distinct OB regions, creating spatially restricted cellular domains. Such regional organization could provide a substrate for convergence across glomerular channels, allowing chemically diverse sensory inputs to recruit shared spatial and cellular domains within the bulb. Although the associated glomerular inputs and synaptic connectivity remain to be defined, our data support a domain-like OB architecture in which projection neuron subtypes tile the outer layer, while interneuron subtypes occupy partially structured niches within the intermediate layer. This arrangement may allow interneuron subtypes to organize the integration of information across specific combinations of glomerular channels. Thus, interneuron subtypes may control the flow of information between projection neuron territories and thereby shape activity patterns to represent biologically relevant information such as behavioral preference.

### Odor preference as one axis of olfactory bulb organization

Across species, the OB receives reproducible glomerular activity patterns that reflect aspects of odorant chemistry. Our atlas extends this view by suggesting that OB organization reflects not only chemotopic sensory input, but also odor preference as one behaviorally relevant axis. At the level of population activity, chemically diverse odors that elicited similar preferences converged on shared spatial activity patterns in the OB (Fig. 1B, 2B-E). This population-scale view, provided by CaMPARI2-based activity mapping, complements time-resolved calcium imaging by revealing spatially distributed response structure. Together, these observations suggest that OB population activity captures not only chemical features, but also a behaviorally relevant dimension.

At the cellular level, odors that drive similar behavioral responses converge on shared transcriptionally defined neuronal subtypes (Fig. 5B-H; Fig. S9C-E). Among the subtypes identified, the short-axon-like *th^+^/gad1b^+^*interneuron subtype i-OB(24) showed the most consistent preference-associated recruitment, with preferential activation by the chemically distinct attractive odors TCA and cytidine. Additional subtypes, including i-OB(6), i-OB(18), and the *tac3b^+^* projection neuron subtype e-OB(25), also showed recruitment biases. Nevertheless, broader sampling across attractive and aversive odorants will be required to determine whether these associations reflect general preference-related recruitment or odorant-specific sensitivity. More generally, because many OB subtypes could only be confidently identified through multi-gene combinations, comprehensive mapping of odor responses onto transcriptionally defined cell types remains an important goal for future work.

Notably, preference-biased recruitment was particularly evident among interneuron subtypes. This observation aligns with work linking inhibitory circuit dynamics to behaviorally relevant changes in projection neuron output ^117^ and with evidence that synchronized mitral-cell activity can carry learned odor-value information ^118^. In larval zebrafish, interglomerular connectivity in the OB is mediated largely by interneurons, with reciprocal PN-IN-PN motifs overrepresented in the circuit ^26,106^. Such an arrangement places interneurons in a strong position to integrate chemotopically organized input and shape projection neuron output across glomerular channels ^119–124^. In our system, targeted ablation of the TH^+^ interneuron population selectively impaired attraction to chemically diverse odors while sparing aversive responses and baseline locomotion (Fig. 6C-G, Fig. S9F-H), suggesting that this population contributes to attractive odor-guided behavior. Whether TH^+^ interneurons exert this effect through direct modulation of projection neuron output, through interactions with other interneuron subtypes, or through a combination of both remains to be determined. Together, these results suggest that early olfactory circuits are organized not only to represent odor chemistry, but also in ways that may support preference-biased transformations of sensory input.

Several models have been proposed to explain innate odor valence, including models in which specific receptors or glomeruli contribute to attraction or aversion ^34,125^, and models in which valence-related representations emerge primarily in downstream brain regions ^44–46,48,126–130^. The view emerging from our study is consistent with work suggesting that odor pleasantness or valence is already represented at early stages of olfactory processing, and that OB spatial organization extends beyond a simple chemotopic map of physicochemical features ^85,131^. More broadly, behaviorally relevant representations may emerge progressively along sensory pathways, including through discrete interneuron bottlenecks early in sensory processing, as recently shown in Drosophila gustatory circuits ^132^. Together, our findings suggest that the OB represents an early node in the progressive emergence of behaviorally relevant representations along the olfactory pathway.

## Author contribution

O.M., R.W.F., and A.F.S. conceptualized the study. O.M. performed data curation, formal analysis, methodology development, validation, and visualization. J.N.A., Y.W., V.H., and S.P. contributed to methodology. R.W.F. and A.F.S. provided supervision and mentorship. O.M., R.W.F. and A.F.S., acquired funding. O.M. and A.F.S. wrote the original draft. All authors contributed to writing, reviewing and editing.

## Acknowledgements

We thank Adam Mazur and the Research IT Core Facility at the Biozentrum for expert advice and computational support in data analysis; all members of the Imaging Core Facility (IMCF) at the Biozentrum, particularly Oliver Biehlmaier, Alexia Loynton-Ferrand, Kai Schleicher, and Sebastien Herbert—for guidance and technical assistance with image analysis; Agata Nowacka and Peter Scheiffele for access to the two-photon microscopy setup and for their valuable guidance; Stella Stefanova and the Biozentrum FACS Core Facility for assistance with cell sorting; the Genomics Facility Basel, especially Christian Beisel and Mirjam Judith Feldkamp, for support with library preparation and sequencing; the Mechanical and Electrical Workshops and the Research Instrumentation Facility at the Biozentrum—particularly Patrick Schlenker, Simon Saner, and Ludovit Pavel Zweifel—for their expertise in designing and implementing the behavioral rig. We are grateful to Thomas Frank and Bethan Jenkins for insightful discussions, sharing of preliminary data, critical reading of the manuscript and thoughtful collaboration that strengthened this work. We thank Johannes Kappel, Lukas Anneser, and Ruth Montano for insightful discussions and critical reading of the manuscript, Ruth Montano for generously sharing preliminary information, and members of the Schier and Friedrich laboratories for insightful discussions, collegiality, and continued support throughout the project.

O.M. was supported by an EMBO Postdoctoral Fellowship (ALTF 572-2020).

This work was supported by SNSF grant 310030_212236 awarded to R.W.F.

## Declaration of generative AI and AI-assisted technologies in the writing process

During the preparation of this work, the authors used ChatGPT (OpenAI), Gemini (Google), and Claude (Anthropic) to assist with language editing and code generation. After using these tools, the authors carefully reviewed and edited all outputs and take full responsibility for the content of the published article. All scientific concepts, analyses, interpretations, and conclusions were conceived, implemented, and verified by the authors.

## Methods

### Zebrafish Husbandry and Transgenic Lines

All experiments were performed on Danio rerio larvae raised at 28.5°C on a 14:10 h light/dark cycle in E3 medium. Larvae were used at 5 days post-fertilization (dpf), prior to feeding and sexual differentiation. To enrich for olfactory bulb (OB) and telencephalic neurons, we used the Tg*(-8kb:cldnb:lynEGFP)* line ^91^. For activity-based labeling, we generated a stable Tg(HuC(elavl3):CaMPARI2) line by Tol2-mediated transgenesis. Briefly, 25pg of pTol2-elavl3:CaMPARI2 plasmid (Addgene #137185; a gift from Eric Schreiter) was co-injected with 25pg Tol2 transposase mRNA into one-cell stage TLAB embryos. Injected embryos were raised to adulthood and screened for germline transmission based on pan-neuronal CaMPARI2 expression in F1 progeny. Stable carriers were subsequently maintained by outcrossing.

To improve optical clarity for imaging, Tg(elavl3:CaMPARI2) and other reporter lines were crossed into the nacre^-/-^ background ^133^. Wild-type larvae for behavior were obtained from TLAB incrosses (generated by TL × AB outcrosses). For spatial transcriptomics, TLAB fish were outcrossed to nacre^-/-^ mutants and subsequently incrossed to generate homozygous nacre^-/-^ mutants. Dopaminergic neurons were visualized using the th^m1512Tg^(2A-QF2); Tg(QUASr:GFP)^c403, 116^, hereafter referred to as th:QF2>GFP, outcrossed to nacre^-/-^ mutants. All procedures complied with institutional and national animal care guidelines.

### Single-cell RNA Sequencing

#### Tissue Dissection and Dissociation

To enrich for OB neurons, we dissected ∼30 telencephala from 5 dpf Tg(*-8kb:cldnb:lynEGFP*) zebrafish larvae under a stereomicroscope in ice-cold 1× Ringer’s solution. Dissected brains were immediately transferred into 2mL Lo-Bind tubes (Eppendorf #0030108078) containing ∼50μL of Ringer’s and kept on ice. Tissue dissociation was performed in 1mL of enzyme solution containing 10mg/mL BI protease (Sigma #P5380), 0.5mM EDTA, 25μL of 20mg/mL DNase I (Sigma #10104159001), and 1× B-27 supplement (Thermo Fisher 17504044) in DPBS. Samples were incubated on ice for 30–45minutes, with gentle trituration using a P1000 pipette every 3 minutes to promote dissociation. Digestion was quenched by adding 400μL of STOP solution (30% FBS- Thermo Fisher 10082147, 0.8mM CaCl₂, 1× DPBS), and suspensions were filtered sequentially through 70μm (pluriStrainer®-43-10070-70) and 40μm (Sigma #BAH136800040) strainers. Cells were pelleted by centrifugation at 500g for 6 minutes at 4°C, washed twice in 1× DPBS with 5% FBS, 0.8mM CaCl₂, and 1× B-27, and resuspended in 80μL 1× DPBS with 5% FBS, 0.8mM CaCl₂, and 1× B-27. Cell viability and concentration were assessed by DAPI exclusion and manual hemocytometer counting.

#### Fluorescence-Activated Cell Sorting (FACS) and Library Preparation

GFP^+^ cells were isolated via fluorescence-activated cell sorting (FACS) and loaded onto the Chromium Single Cell 3′ v3.1 platform (10x Chromium Controller, 10x Genomics). Single-cell cDNA and sequencing libraries were generated following the manufacturer’s protocol. Libraries (Chromium Single Cell 3’ v3.1) were sequenced on an Illumina NovaSeq 6000 S4 flowcell using the recommended read lengths.

#### Preprocessing and Quality Control

Raw sequencing reads were processed using CellRanger ^134^ with alignment to the zebrafish reference transcriptome (GRCz11), and processed in RStudio mainly using the Seurat package v4.3.0 ^135^. Cells with fewer than 200 or more than 3,000 detected genes or >10% mitochondrial content were excluded (Fig. S5A). Data were log-normalized (scale factor=10,000), and the top 2,000 highly variable genes were identified using Seurat and the *vst* method. Principal component analysis (PCA) was performed on these genes, and graph-based clustering was executed using the Louvain algorithm.

#### Selection of OB-Enriched Postmitotic Neurons

To isolate OB-enriched neuronal populations, we applied a two-stage filtration strategy. In the first stage, dividing and non-neuronal cells were removed by clustering at low resolution (resolution = 0.2, 30 PCs). This yielded 14 clusters: 2 corresponding to replicating cells, 5 to non-neuronal cell types, 6 to neurons, and 1 to glia. Neuronal identity was determined by the expression of broad neuronal markers (*elavl3, snap25a, slc17a6a/b, slc32a1, gad1b/gad2*) and the absence of proliferative or progenitor markers (*pcna, her4.2, dla, tal1, uncx*) ^56,98,101,136–143^ (Fig. 3B-B’).

To remove immature and enrich for OB neurons, the neuronal subset was re-clustered after normalization using PCA and UMAP (resolution = 0.1, 30 PCs), producing 9 clusters classified as inhibitory/excitatory neurons (*snap25a⁺, slc17a6a/b⁺, slc32a1⁺, gad1b/2⁺, tal1⁻, uncx⁻, dla⁻, her4.2⁻*), immature inhibitory/excitatory neurons (same neuronal markers with *tal1⁺, uncx⁺, dla⁺, her4.2⁺*), glia (*gfap⁺*; ^144^), or neuroendocrine (*pomca⁺, cga⁺*; ^145–147^ ; Fig. 3C-C’). The final filtered dataset comprised 2,071 postmitotic neurons enriched for OB identity (Fig. 3D-D’).

#### Clustering and Validation

To optimize clustering resolution, we adopted a similar approach as previously described ^93^. We evaluated multiple parameter sets and selected the solution with the highest silhouette score (Fig. S5B), yielding 26 clusters. To assess whether transcriptional clusters could be recovered from transcription factor expression alone, we trained a random forest classifier using only transcription factors (KEGG dre03000) or ribosomes (KEGG dre03010). Cells were randomly split into training and test sets (70/30), and a classifier was trained to predict cluster identity based on normalized expression.

#### Minimal Gene Set for OB Subtype Recovery

To estimate the minimal number of genes required to recover OB transcriptional subtypes in the context of the full single-cell dataset, we trained random forest classifiers using cluster labels transferred from the OB-enriched subset. Classifiers were trained on normalized expression values using highly variable genes (HVGs) or transcription factors as input features and evaluated using a 70/30 train–test split.

Genes were ranked by importance (MeanDecreaseAccuracy) from a random forest trained on the top 2000 HVG set or full transcription factor repertoire, and classifiers were re-trained using incrementally increasing subsets of the top-ranked genes. Prediction performance was quantified as OB subtype classification accuracy, computed by restricting evaluation to OB-labeled cells in the held-out test set. Classification accuracy increased with gene number and approached a plateau (∼87% accuracy at ∼32 genes; Fig. S5F). We defined the minimal gene set as the smallest subset achieving ≥ 95% of maximal accuracy. This analysis indicates that reliable recovery of OB transcriptional identities in the unfiltered transcriptomic space requires the coordinated expression of multiple genes rather than any single marker.

#### Inter-individual Variability in Spatial Gene Expression Patterns

To assess variability of gene expression patterns across samples, we analyzed the mapzebrain dataset, in which each gene was independently measured in at least three animals ^89,148^. Pairwise Pearson correlations were computed across biological replicates for each gene, with and without Gaussian blurring (σ = 0–10μm) and correlations were averaged across pairs of comparisons. Average correlation plateaued near r=0.8 at σ = 10μm (Fig. S5G), consistent with reported spatial registration precision (∼0.22–14μm; ^89^). These results suggest that independent measurements of marker genes across samples are insufficient to reconstitute single-cell clusters.

### Whole Mount Spatial Transcriptomics

#### Target Gene Selection

To enable robust, cell-level identification of Seurat-defined neuronal subtypes and permit multiplexed co-localization, we curated a compact, information-rich gene panel. Candidate markers were drawn from three complementary sources: (1) the top three transcription-factor markers per subtype from our scRNA-seq clustering, (2) the most informative genes identified by a random-forest classifier (Fig. S5C’’), and (3) broad neuronal markers. From an initial pool of ∼200 candidates we ranked genes by random-forest importance and selected the top 68 for visualization and spatial profiling (Supplementary Table 3; Figs. S6-7, Movie S3).

#### Primary and Secondary Probe Design

Primary probes were designed following a previously described pipeline ^100^. In brief, each primary probe set comprised 18–60 distinct 40-mer target sequences complementary to the mRNA of a given gene. We first built a 17-nt hash table enumerating the occurrence of every 17-mer in the unspliced zebrafish transcriptome (GRCz11, NCBI RefSeq). We then scanned each candidate 40-mer sequence for a given gene transcript and estimated its off-target potential based on the corresponding 17-mer counts. Candidate 40-mers were filtered using the same selection criteria described previously ^100^. For each 40-mer probe, we appended three unique gene-specific 20-nt readout sequences and flanked the construct with two 20-nt PCR primer sites. Candidate readout sequences and PCR primers were selected as previously described ^149^ and screened against the zebrafish transcriptome to minimize cross-hybridization.

Primary probe pools were synthesized as oligonucleotide pools (Twist Biosciences) and prepared as previously described ^150^. Briefly, pools were amplified by limited-cycle PCR using the forward and reverse primers described above and purified using the Zymo Research Oligo Clean & Concentrator kit (D4003). Purified DNA was used as a template for in vitro transcription with T7 polymerase (NEB, E2040S), and the resulting RNA products were reverse-transcribed into single-stranded DNA using Maxima Reverse Transcriptase (Thermo Scientific, EP0751). RNA was removed by alkaline hydrolysis, and DNA oligos were purified using the Zymo Research Oligo Clean & Concentrator kit (D4006). To enable covalent incorporation of primary probes into acrylamide gels, we used 5′-acrydite modifications on both the forward PCR primer and the primer used during reverse transcription, yielding acrydite-tagged primary probes.

Linker oligonucleotides were gene-specific and consisted of a 20-nt sequence reverse-complementary to the gene’s readout sequence plus two tandem repeats of the readout probe binding site (20-nt each) ordered from IDT.

Readout probes were 20-nt oligonucleotides labeled at both the 5′ and 3′ ends with fluorescent dyes: Cy3 for genes imaged with a 561-nm laser and Cy5 for genes imaged with a 638-nm laser. Readout oligos were HPLC-purified and ordered from Eurofins Scientific. Sequences for all linker probes, readout probes, and primers are provided in Supplemental Table 3, primary probe sequences are deposited at Zenodo.org (10.5281/zenodo.19607911).

#### Sample Preparation and Probe Hybridization

Spatial transcriptomic profiling was performed using sequential fluorescence in situ hybridization (seqFISH), based on a previously described weMERFISH protocol ^100^ adapted here for whole-mount larval zebrafish brain tissue. Five dpf larvae were fixed overnight at 4°C in 4% PFA prepared in PBS containing 0.1% Triton X-100 (PBS-T), then washed three times for 5 min each in PBS-T at room temperature. Samples were dehydrated through a methanol series in PBS-T (25%, 50%, 75%, and 100% MeOH) and stored overnight at −20°C. Larvae were rehydrated for 10 min in 50% MeOH/50% 2× SSC containing 0.1% Triton X-100 (2× SSC-T), followed by three 10 min washes in 2× SSC-T at room temperature. Brains were subsequently dissected under a stereomicroscope.

Pre-hybridization was carried out for 30min at room temperature in 40% formamide in 2× SSC-T supplemented with RNase inhibitor (New England Biolabs #M0314S at 1:100 dilution). Samples were then incubated in the primary probe library eluted in hybridization buffer consisting of 10% dextran sulphate and 50% formamide in 2× SSC-T (with RNase inhibitor 1:100) for 48–60h at 47°C with gentle agitation. After hybridization, brains were washed in 40% formamide in 2× SSC-T for 1h at room temperature, followed by five 5 minute washes in 2× SSC-T, and were kept in 2× SSC-T until hydrogel embedding.

#### Gel Embedding and Tissue Clearing

For immobilization and optical stabilization, brains were embedded in a 4% acrylamide/bis-acrylamide (A/BA) hydrogel. Coverslips were first functionalized by applying a bind-silane solution (1% bind-silane in ethanol, pH adjusted to ∼3.5 with glacial acetic acid Sigma #A6283) to one side, heat-curing at 90 °C for 30 min, and cooling to room temperature. Immediately before sample mounting, the silanized surface was coated with 2 µL poly-L-lysine (Sigma, #P8920) and allowed to air dry.

For hydrogel embedding, brains were incubated overnight at 4 °C in freshly prepared and degassed hydrogel solution (1 mL 40% A/BA stock; Bio-Rad, #1610144), 0.6 mL Tris-HCl pH 8.0, 0.6 mL 5 M NaCl, and Milli-Q H₂O to 10 mL total volume, supplemented with RNase inhibitor (1:5000). The following day, samples were washed once with fresh hydrogel solution and oriented on the poly-L-lysine–coated coverslips.

Immediately before casting, 10% ammonium persulfate (APS; Sigma, #A3678) and TEMED (Thermo Fisher, #17919) were added to the hydrogel (25 µL APS and 2.5 µL TEMED per 5 mL solution). Approximately 150 µL of this initiator-supplemented gel was dispensed onto a GelSlick-coated glass slide (RUWAG Handels AG, #LZ-50640). A 750 µm gasket (Bioptechs, #1907-1422-750) was used to form a sandwich with the coverslip. Polymerization was performed for at least 3 h at room temperature in a nitrogen-filled chamber, after which the coverslip was gently separated from the casting slide.

Hydrogel-embedded specimens were washed three times in 2× SSC-T, then cleared overnight at 47 °C in 2.5% SDS (Invitrogen, #AM9820) with Proteinase K (0.4 mg/mL; Thermo Fisher, #AM2548) in 2× SSC. Samples were washed three times in 2× SSC-T to remove residual SDS and stored in 2× SSC-T until readout hybridization and imaging.

#### Sequential Hybridization and Imaging

Sequential *in situ* hybridization and imaging were carried out following the previously defined weMERFISH workflow ^100^ with adaptations for whole-mount larval zebrafish brains. Readout hybridizations were performed using a custom fluidics setup adapted from previously described MERFISH-style systems ^150,151^ and integrated with a Nikon Ti2 inverted microscope under custom macro control. For each hybridization round, two genes were detected by flowing ∼1.5mL of gene-specific linker oligonucleotides (100nM per probe) diluted in 35% formamide in 2× SSC-T and incubating for 15–20min at room temperature. Excess linker was removed by three washes in ∼2mL of 30% formamide in 2× SSC-T (3min incubation per wash), after which ∼1.5mL of Cy3/Cy5 fluorescent readout probes (200nM each) supplemented with DAPI was introduced and incubated for 15min. A second set of three 3-min washes in 30% formamide in 2× SSC-T was then performed, followed by 3mL of 2× SSC-T to remove residual formamide, and finally ∼2mL of imaging buffer (2× SSC, 50mM Tris-HCl pH 8, 10% w/v glucose, 2mM Trolox, 0.5mg/mL glucose oxidase, 40µg/mL catalase, 4µg/mL DAPI) was flown in. After buffer exchange, flow was halted for 1 min to allow for sample stabilization and imaging commenced.

Images were acquired on a Nikon Ti2 equipped with a CSU-W1 spinning disk confocal scanner (50µm pinhole) and a Plan Apo VC 60× water-immersion objective (NA 1.2). Exposure times were 500ms (Cy3), 1s (Cy5), and 50ms (DAPI) per plane. Volumes were collected at 0.11µm × 0.3µm (xy × z) sampling, spanning ∼750 z-planes to capture the full tissue thickness preserved during embedding. Focus stability during multipass imaging was maintained using Nikon’s Perfect Focus System.

Between imaging rounds, readout probes were stripped by three 5-min incubations in 70% formamide in 2× SSC-T at room temperature, after which samples were washed back into 2× SSC-T before the next hybridization cycle. This hybridize–image–strip sequence was repeated until all readouts in the 68-gene panel had been imaged.

#### Image Processing, Spot Detection and Assignment

Raw Cy3 and Cy5 gene-channel images were first processed by flat-field correction and then deconvolved using the point spread function (PSF) as previously defined ^100^. Images were normalized, and Gaussian peaks were detected using σXY = 1.85 pixels, σZ = 2.5 pixels, and a minimum intensity threshold of 80. For each detected spot, the pipeline reported intensity values and Gaussian-correlation scores in both raw and deconvolved image space, which were used for manual quality control and downstream filtering. All hybridization rounds were rigidly registered to a DAPI-based reference round using the previously described alignment pipeline^100^.

Cell segmentation was performed on the reference DAPI image using a custom-trained Cellpose model ^152^ trained on sequential slices from the reference z stack and run on the 3D image. Detected transcript coordinates were assigned to segmented cells based on 3D spatial overlap. A final cell-by-gene count matrix was constructed for each brain, with transcript counts normalized to the median cell volume to correct for segmentation variability. Low-quality cells with low total transcript counts or aberrant morphologies were excluded from downstream analysis.

#### Tangram-Based Gene Imputation and Spatial Mapping of Neuronal Subtypes

To infer the fine-grained spatial distribution of transcriptional subtypes, we aligned our single-cell RNA-seq reference dataset to the spatial gene expression map using Tangram ^111^. The full set of 68 measured spatial genes was used as ground truth for model guidance. Prior to integration, spatially resolved cells were filtered to include only neurons based on the expression of canonical glutamatergic and GABAergic markers (*slc17a6a/b, slc32a1, gad1b, elavl3*), ensuring that downstream alignment operated on the same neuronal population represented in the scRNA-seq reference.

Tangram was first applied in gene-imputation mode, enabling direct comparison between predicted and experimentally measured spatial expression patterns. Imputation accuracy was evaluated using a leave-one-gene-out (LOGO) cross-validation scheme, in which each of the 68 genes was iteratively withheld from training and subsequently predicted from the remaining genes (Fig. S8A,C). Predicted and measured spatial patterns were compared using Pearson correlation to estimate per-gene reconstruction fidelity. As described in the Results, most genes showed measurable predictive agreement, and qualitative inspection confirmed that Tangram recovered appropriate spatial domains even for genes with modest correlation values. Predictions for genes outside the measured panel were also compared with independent spatial reference atlases, including mapzebrain,^89,148^, providing external validation of inference consistency (Movie S5).

After gene-level validation, Tangram was applied in ‘clusters’ mode to project Seurat-defined neuronal subtypes into spatial coordinates. Subtype identities were assigned to spatially resolved cells, and reproducibility across animals was assessed by comparing subtype composition profiles between brains (Movie S4).

#### Quantification of Subtype Spatial Cohesion and Territorial Overlap

To quantify the spatial organization of transcriptional subtypes we employed a 3D nearest-neighbor cohesion score. For each cell, the fraction of *k* nearest neighbors sharing the same Tangram-imputed subtype identity was calculated. We tested a range of *k* values (*k* = 2–500) and found that subtype-level median cohesion rankings stabilized for *k* ≥ 30 (Spearman ρ > 0.99 across *k* values; Fig. S8F). Based on this stability, *k* = 30 was chosen for subsequent analyses as a measure of local compactness.

To independently assess spatial intermingling between subtypes, we modeled three-dimensional spatial territories using Gaussian kernel density estimation with a fixed bandwidth (bw) of 15µm. For each brain, cell coordinates were mapped onto a 10µm voxel grid. For each subtype, we extracted the highest-density region enclosing 90% of the total probability mass to generate a binarized volumetric territory. To ensure spatial cohesion, isolated fragments smaller than 25 voxels were removed. Pairwise territorial relationships were quantified using the Jaccard index, Dice coefficient, and overlap coefficient to distinguish between discrete spatial tiling and modular integration.

### Odor Preference Assay

#### Behavioral Setup, Odor delivery, and Odorants

To quantify naïve odor preferences in larval zebrafish, we developed a custom-built, flow-based two-choice behavioral assay with automated laminar flow control and odor delivery (diagram and blueprints in Supplementary Information). The assay chamber was housed within an outer enclosure adapted from previous designs, with a top-mounted camera and infrared and white LED illumination from below ^153^ and consisted of two adjacent flow lanes: one carrying E3 embryo medium as vehicle control and the other delivering an odor solution. Flow was established using dual-head peristaltic pumps (Masterflex® L/S® Digital Drive MFLX07522-30 with pump heads #7535-08 and #7519-06) to maintain constant unidirectional flow (Fig. 1A, Fig. S1A). Inflow lines were connected to narrow tubing ports affixed to opposite sides of the chamber, whereas outflow lines were fitted with wider tubing to prevent overflow and routed flow to the drain. To characterize laminar flow speed and profile, IR dye (Sigma #543349) was perfused through one side of the arena, and dye dynamics were quantified within fixed ROIs. Flow speed was estimated from the delay in dye saturation between upstream and downstream ROIs (∼3 mm/s). Dye perfusion confirmed stable, largely non-mixing flow streams across the arena. Flow separation was validated visually after each odor trial by flowing IR dye through one side of the arena.

Odor selection and delivery were coordinated through two solenoid valve manifolds (Masterflex #MFLX01356-20), each capable of routing one of seven odorants or E3 to the corresponding chamber inlet. Valve switching and flow timing were controlled through a custom electronics interface integrating Arduino microcontrollers and a LabVIEW GUI (see Supplementary Data). Each experimental trial consisted of a 10 min baseline habituation period, a 2 min odor delivery phase, and an 8 min washout period. At the end of each experimental day, tubing used for each odorant was flushed with 0.3% H2O2 to minimize odor carryover between experiments.

A panel of 32 odorants, including amino acids, bile acids, nucleotides, and complex odorants, were tested. Several of these were assayed at multiple concentrations, yielding 46 defined odorant conditions, in addition to a spontaneous activity/control condition (Supplementary Table 1). All odorants were prepared fresh on the day of testing by dissolving them in buffered E3, and neutral pH (∼7) was verified using pH strips (VWR #1.09531.0001). As controls, we used either a continuous stream of E3 from the main reservoir, referred to as the spontaneous condition, or E3 routed through an alternative solenoid valve, to assess the effect of solenoid valve switching.

E3 was prepared by diluting a premade 50x stock solution to 1x, yielding final concentrations of 5.0 mM NaCl, 0.169 mM KCl, 0.33 mM CaCl₂·2H₂O, and 0.33 mM MgSO₄·7H₂O, and buffered with 2.4 mM sodium bicarbonate.

Food odor was generated by incubating 250 mg of Zebrafeed <100 (Sparos) in 50 ml of E3 for 1-2 hours, filtering the solution first through filter paper (VWR #516-0289) and then through a 0.45 µm syringe filter (Sarstedt #83.1826), and diluting it 1:100 in E3. Adult and cichlid odors were prepared by placing four adult zebrafish or cichlids (2 males and 2 females) in fresh E3, followed by filtering as described for the food odor, and diluting the filtrate 1:100 in E3. AAmix consisted of alanine (Ala), arginine (Arg), lysine (Lys), and tryptophan (Trp) at 100 μM each, while BAmix contained taurocholic acid (TCA), taurodeoxycholic acid (TDCA), and glycocholic acid (GCA) at 100 μM each.

#### Behavioral Recording and Preprocessing

Larvae (5dpf) were tested individually, and each trial was recorded at 10Hz using a high-speed monochrome camera (Grasshopper3, Flir #GS3-U3-23S6M-C) fitted with an IR filter. Videos were processed using ZebraZoom ^154^, an automated tracking software configured to extract head and tail positions at frame-level resolution (configuration file available in Supplementary Materials). Trials were discarded if they exhibited: (1) prolonged immobility (>150 seconds, n = 126), (2) excessive missing data (>50% of dropped frames during odor exposure, n = 22), or (3) significant pre-odor side bias (defined as >70% occupancy on one side during the pre-odor stage, n = 181). After exclusion, a total of 1,692 larvae remained for analysis.

#### Behavioral Quantification

Movement bouts were identified using peak detection with a speed prominence threshold of 0.2cm/s. Heading angles were calculated as the angle between the head position and the most distal visible tail point. For each trajectory, we computed a set of behavioral features: swim speed, interbout interval, distance between bouts, and heading change. Outlier values for all parameters were filtered using interquartile range thresholds (e.g., 1.5x IQR above Q3 or below Q1). Thigmotaxis (wall-following behavior) was quantified by measuring the time spent near the walls of the arena binned over time. We verified that larvae’s behavior remained naturalistic in this environment: they exhibited typical thigmotaxis (preferring arena edges) as well as rheotaxis (tending to orient with flow), yet did not remain fixed at the flow source (Fig. S1B). A 10-minute acclimatization period was selected based on spontaneous swim trials, where thigmotaxis stabilized after ∼560 seconds (Fig. S1B). Median metrics were computed separately for the odor and control sides (defined by the larva’s normalized Y-position relative to the midline).

#### Preference Calculation and Statistical Analysis

Odor preference was quantified as the proportion of time each larva spent on the odor side of the arena (normalized y > 0.5) during the 2 min odor exposure window. For Figure 1B, statistical significance was assessed by comparing each odor condition to the E3 control group. Depending on distribution normality, assessed using the Shapiro-Wilk test, comparisons were performed using either independent t-tests or Mann-Whitney U tests. P-values were adjusted for multiple comparisons using Bonferroni correction. Asterisks in Figure 1B denote significance relative to the E3 control group (*p < 0.05, **p < 0.01, ***p < 0.001). To assess whether locomotor features scaled with odor preference, we performed linear regression analyses between per-odorant median preference scores and extracted swim parameters. On the odorant side, preference scores were significantly correlated with heading change (R^2^ = 0.35, p = 2.7×10^-5^), inter-bout interval (R^2^ = 0.09, p = 0.05), inter-bout distance (R^2^ = 0.09, p = 0.05), and swim speed (R^2^ = 0.2, p = 0.002). In contrast, no significant relationships were observed on the control side, indicating that these behavioral modulations are specific to odor-guided sampling (Fig. 1C).

To test whether odor chemical class explains behavioral preference, we performed a one-way ANOVA on preference scores grouped by odor class, excluding complex odors. An Ordinary Least Squares (OLS) model was used, and effect size was quantified by R^2^. Odorants were grouped according to chemical class. To assess whether the observed effect exceeded chance expectations, we performed a permutation test in which odor-class labels were randomly shuffled across observations (10,000 iterations), and the ANOVA was recomputed for each permutation to generate a null distribution of F-statistics and R^2^ values. Empirical p-values were calculated as the proportion of permutations yielding values greater than or equal to the observed statistic.

#### Clustering of Post-Odor Behavioral Preferences During Odorant Washout

To identify shared patterns of post-odor behavioral dynamics, we clustered odors based on larval position during the washout phase. For each trial, normalized y-positions were binned into 10 second intervals, and the mean per-bin value was computed. Aggregated traces were summarized by odor (median across larvae). Odors were hierarchically clustered using Ward’s linkage on Euclidean distances, revealing structure in recovery trajectories not captured by immediate preference.

### CaMPARI2-Based Activity Mapping in Freely Swimming Larvae

#### Rationale and Transgene Expression

To map odor-evoked neuronal activity at cellular resolution, we used CaMPARI2, a photoconvertible calcium integrator that irreversibly switches from green to red fluorescence in active neurons exposed to 405nm light. Although odor representations in the zebrafish OB involve dynamic temporal patterns ^155^, multiple studies report that these responses stabilize within seconds and persist during sustained odor exposure ^22,73,156,157^, making CaMPARI2 well-suited for capturing spatial activity maps.

While HuC-driven transgene expression was previously suggested to preferentially label projection neurons ^22^, we observed broad expression across the OB, including deeper GABAergic layers (Fig. S3A). Reduced signal in medial regions likely reflects lower expression in progenitor populations, consistent with *pcna* localization in the mapzebrain and Wullimann atlases ^56,148^. To account for baseline expression differences and interindividual variability, activity was quantified using red-to-green (R/G) CaMPARI2 fluorescence ratios on a per-pixel basis.

#### Photoconversion During Odor Stimulation in Freely Swimming Larvae

We designed a custom adapter for UV light delivery to enable CaMPARI2 photoconversion during behavioral odor stimulation in freely swimming larvae (Fig. 2A; supplemental materials). Larvae were exposed to odorants for 20 seconds, allowing the arena to saturate, followed by a 10-second pulse of 405 nm UV light at ∼60 mW/mm² at the level of the larvae, distributed evenly across the behavioral arena. This same protocol was used for both CaMPARI2 imaging and CaMPARI2-seq experiments (Fig. 2A, Fig. 5A).

#### Sample Preparation, Live Imaging, and Image Registration

After exposure, larvae were anesthetized in Tricaine (MS-222 Sigma #E10521) and embedded in 2% low-melt agarose (Sigma #A9414). Whole-OB volumes were imaged using a Yokogawa spinSR CSU-W1 spinning disk confocal microscope equipped with a 30×/1.05 NA silicon immersion objective. Z-stacks were acquired at 1μm steps with excitation at 488nm (200ms dwell) and 561nm (500ms dwell). Images were captured at 0.43μm XY resolution.

To enable cross-animal comparisons, volumes were registered to a common local reference using CMTK with the following parameters: -T 32 -awr 010203 -X 52 -G 80 -R 3 -A ’--accuracy 0.8’ -W ’--accuracy 1.6’ -s.

The local reference was then registered to the mapzebrain atlas (standard_brain_fixed_lynTagRFP) using ANTs. The ANTs registration command used was:

antsRegistration -d 3 --float 1 -o [${outputPrefix},1] --interpolation WelchWindowedSinc --use-histogram-matching 1 \

-r [${template},${input1},1] \

-t rigid[0.1] -m MI[${template},${input1},1,32,Regular,0.25] -c [200x200x200x0,1e-8,10] \

-x [${mask_fixed},NULL] --shrink-factors 12x8x4x2 --smoothing-sigmas 4x3x2x1vox \

-t Affine[0.1] -m MI[${template},${input1},1,32,Regular,0.25] -c [200x200x200x0,1e-8,10] \

--shrink-factors 12x8x4x2 --smoothing-sigmas 4x3x2x1 \

-t SyN[0.1,6,0.4] -m CC[${template},${input1},1,2] -c [200x200x200x200x10,1e-7,10] \

--shrink-factors 12x8x4x2x1 --smoothing-sigmas 4x3x2x1x0 \

--winsorize-image-intensities [0.005,0.995]

Transformed coordinate files were generated using antsApplyTransformsToPoints. CaMPARI2 spot positions were scaled to physical units, Z-flipped to match atlas orientation, and batch-transformed into atlas space, enabling voxel-wise analyses across datasets.

#### Cell Segmentation and Activity Quantification

Individual neurons were segmented from registered CaMPARI2 imaging volumes using a custom-trained Cellpose model. The model was trained on manually annotated reference slices and applied to the full 3D image stack.

To enrich for OB associated neurons, we leveraged the invariant glomerular anatomy ^55^ as a spatial filter. Glomeruli were manually segmented in a reference brain by labeling nuclei with the nuclear dye Draq5 (1 μM in E3 overnight; LubioScience #17558) in live HuC:CaMPARI2 larvae. These larvae were imaged using the same acquisition parameters as for CaMPARI2 experiments, registered to the reference volume, and nuclei-negative regions corresponding to glomerular structures were delineated. Guided by electron microscopy reconstructions showing that OB neuron somata are typically positioned close to the glomeruli they innervate ^26,106^, we used the resulting glomerular masks as anatomical landmarks to filter the neuronal dataset. Cells were retained if they were located within 10 μm of a glomerulus or fell within the expected somatic volume range (100–1,000 μm³), thereby enriching for anatomically plausible OB neurons while excluding clear segmentation artifacts.

Activity values were computed as the red-to-green (R/G) fluorescence ratio at the pixel level and averaged per segmented cell (Fig. S4). To generate odor-evoked activity maps, we selected the top 10% of cells ranked by activity within each dataset. This within-brain percentile threshold was used to identify the most strongly photoconverted cells while reducing sensitivity to inter-animal differences in baseline expression, photoconversion efficiency, and imaging conditions. Each odor activity map was derived from a single brain exposed to a single odor during CaMPARI2 photoconversion, and odor presentations were repeated across multiple animals to assess reproducibility.

#### Activity Pattern Correlations and Clustering

To compare odor-evoked population activity maps, we computed 3D kernel density estimates (KDEs) for each odor using a Gaussian kernel (bandwidth = 5μm, voxel size = 2.5μm). KDE volumes were masked to retain the top 90% of cumulative density and compared pairwise by voxelwise Pearson correlation, yielding an odor–odor similarity matrix. Similarity matrices were hierarchically clustered (average linkage). Clusters containing fewer than two trials were excluded from downstream analyses. Cluster separation was quantified using silhouette analysis computed from correlation-derived distances (1 − Pearson r). Mean silhouette scores were calculated across trials to assess geometric separation between clusters. Full list of odors, cluster assignment, silhouette value and cluster valence label available in Supplemental Table 2.

#### Cluster Stability and Odor Consistency Evaluation

Cluster robustness was evaluated using nonparametric bootstrap resampling of trials (1,000 replicates). For each replicate, trials were resampled with replacement, the similarity matrix was reclustered, and co-cluster probabilities were computed for all trial pairs. Cluster stability was summarized as the mean within-cluster co-cluster probability. Cluster stabilities were: Cluster 1 = 0.52, Cluster 2 = 0.79, Cluster 3 = 0.72, Cluster 4 = 0.99, Cluster 5 = 0.73.

Reproducibility across individuals was quantified for each odorant–concentration pair as the majority-cluster ratio, defined as the fraction of trials assigned to the modal cluster for that odor. Across odors, the median ratio was 0.88 (IQR 0.67–1.00; range 0.45–1.00).

To test whether activity maps were more similar within odors than between odors, voxelwise correlations were compared for within-odor versus between-odor trial pairs. Statistical significance was assessed using permutation testing (10,000 label shuffles). Within-odor correlations were significantly higher than between-odor correlations (mean r = 0.171 vs −0.021; p = 1.0 × 10^-4^).

#### Permutation Test Linking Activity Patterns to Behavioral Preference

To test whether clusters corresponded to behavioral valence, we calculated the mean behavioral preference score for all trials within each cluster. Significance was assessed using a permutation test in which odor–preference score pairings were randomly shuffled across trials (10,000 permutations) to generate a null distribution of cluster means. Clusters were classified as attractive, aversive, or neutral based on the direction and significance of deviation from the null distribution. P-values were corrected across clusters using the Benjamini–Hochberg false discovery rate procedure.

#### Odorant Chemical-Class Enrichment

To examine whether activity patterns reflected odorant chemistry, odorants were assigned to clusters based on their majority cluster across trials. A contingency table (cluster × chemical class) was constructed and tested using Pearson’s χ^2^ test. Significance was further evaluated using a permutation test in which chemical class labels were randomly reassigned across odorants (10,000 iterations). Contributions of individual cluster–class combinations were assessed using standardized residuals from the χ^2^ test and one-sided Fisher’s exact tests (alternative = “greater”; Haldane–Anscombe 0.5 correction where required). P-values were corrected across comparisons using the Benjamini–Hochberg procedure. Global association between cluster identity and chemical class was significant (χ^2^(14) = 33.79, p = 0.00221; permutation p = 0.0016; Cramér’s V = 0.943).

#### Odor Activity Pattern visualization

To generate representative activity-pattern maps while preventing odorants with more larvae from dominating the visualization, larvae assigned to each pattern were first averaged within odorant condition. This generated one normalized odorant-level KDE for each odorant represented in the pattern. The odorant-level KDEs were then averaged across odorants to generate the final pattern KDE. Pattern KDEs were max-normalized for visualization, imported into Imaris v10.1, and rendered as horizontal and coronal maximum-intensity projections for Fig. S3D-E and Movie S2.

For three-dimensional scatter renderings, we generated a shared reference cloud by sampling neurons from all odorant conditions and larvae after olfactory bulb proximity and cell-volume filtering. The same reference cloud was reused for every pattern, and each reference neuron was colored according to the value of the corresponding pattern KDE at its spatial coordinate. Dot opacity was scaled with normalized KDE intensity, allowing pattern-enriched regions to be emphasized while preserving identical anatomical sampling across patterns.

#### Cross-Modal Comparison of CaMPARI2 Odor Maps and Two-Photon Brain-Wide Odor Responses

To compare CaMPARI2-derived odor responses with an independent two-photon brain-wide imaging dataset ^88^, we quantified voxel-wise spatial similarity after registration to the mapzebrain reference space. For each odor, CaMPARI2-positive points were filtered by intensity and volume, restricted to the olfactory bulb using a binary OB mask, and converted to 3D coordinates. These coordinates were used to generate kernel density estimate (KDE) volumes (Gaussian kernel, bandwidth 5 µm) evaluated on a shared grid spanning all odors and Jenkins points. The Jenkins dataset provides voxel-wise regression coefficients for positive-and negative-tuned components, which were processed identically to form matching KDE volumes.

Spatial similarity between CaMPARI2 and two-photon KDE maps was quantified using masked voxel-wise Pearson or Spearman correlations, computed only within high-density regions defined by a threshold corresponding to the top 90% of KDE mass across either map.

Statistical confidence was assessed using permutation tests (1,000 shuffles of two-photon voxels) and bootstrap resampling (1,000 iterations) to obtain empirical p-values and 95% confidence intervals.

For each odor, a valence-bias score (ΔR) was computed as the difference between its positive-and negative-tuned masked correlations. Behavioral preference scores were aggregated per odor from two-choice flow assays (median across larvae, with IQR/2 as a variability estimate). ΔR values were then compared to behavioral preference using Spearman correlations to assess monotonic alignment between spatial valence patterns and odor-evoked behavior.

### Sequencing of Odor-Activated Neurons by CaMPARI2-seq

#### Odor Stimulation, Photoconversion, and Cell isolation

To link odor-evoked neuronal activity to transcriptional identity, we used CaMPARI2-seq. Larvae were exposed to one of four conditions, TCA 300 μM, cytidine 100 μM, ATP 500 μM, or E3 vehicle, and photoconverted under UV illumination while odorant was delivered from all sides to saturate the arena. Immediately after photoconversion, larvae were rapidly anesthetized, and telencephala were dissected in ice-cold 1× Ringer’s solution. Tissue was dissociated using the same enzymatic protocol used for the atlas scRNA-seq dataset (see Single-cell RNA Sequencing). CaMPARI2-photoconverted red neurons were isolated by FACS and processed immediately using a modified FLASH-seq protocol (Fig. 5A; Fig. S9A; ^115^).

#### Library Preparation and Sequencing

Single-cell RNA-seq libraries were prepared using the FLASH-seq protocol as previously described ^115^ following the associated protocols.io (https://www.protocols.io/view/flash-seq-protocol-kxygxzkrwv8j/v4). The I.DOT (Dispendix) was used for all dispensing steps, the Agilent Bravo system with 96ST head (Agilent Technologies) and Alpaqua 384 Post Magnet (NimaGen) for magnetic bead clean-up (SPRIselect reagent kit, Beckman Coulter) and the Mosquito HV system (SPT Labtech) for cDNA normalisation and setting up adapter ligation reactions. The RT-PCR reaction was performed with 21 cycles. QC of a subset of the cDNAs and the final library pools was done with the Fragment Analyzer HS NGS 1-6000 bp Kit (Agilent Technologies). Concentrations were measured with Quant-iT PicoGreen (Thermo Fisher Scientific) using the Infinite M1000 Pro (Tecan). The tagmentation reaction was set up with “in-house” Tn5 transposase ^158^ (produced by the Protein Production and Structure Core Facility, EPFL). Indexing primers were ordered from IDT; sequences of primers are available on protocols.io. Library pools were sequenced SR76 with NextSeq 500/550 High Output Kit v2.5 (75 Cycles) (Illumina).

#### Read Processing and Annotation

Reads were aligned to the zebrafish genome (GRCz11; Ensembl v103) using STAR (v2.7.1a), and gene-level counts were generated with featureCounts. Gene annotations were obtained via biomaRt (Ensembl v103), and duplicated gene symbols were resolved by retaining the highest mean–expressed mapping and uniquely renaming lower-expressed entries. Count matrices were imported into Seurat (v4.3.0). Cells with 200–12,000 detected genes and <10% mitochondrial RNA were retained (Fig. S9B), yielding 2,272 high-quality cells (median ∼4,000 genes per cell) from TCA (663), cytidine (498), ATP (580), and E3 controls (531).

#### CaMPARI2-seq Integration with scRNA-seq Atlas and Subtype Enrichment Analysis

To assign transcriptional identities to odor-activated CaMPARI2-seq cells, each CaMPARI2-seq dataset (TCA, cytidine, ATP, and E3 control) was mapped onto the OB single-cell RNA-seq reference atlas generated from 10x Genomics data using Seurat label transfer. Transfer anchors were computed using FindTransferAnchors with the reference PCA reduction (dimensions 1–30) and a feature set comprising zebrafish transcription factor genes together with top marker genes identified in the OB atlas. Reference cluster identities were predicted for each query cell using TransferData, and query cells were projected into the reference UMAP embedding using MapQuery with the reference UMAP model.

Mapping quality was assessed using the Seurat prediction confidence score (prediction.score.max). The distribution of prediction scores was summarized across predicted clusters and experimental conditions to verify consistent mapping performance.

To quantify subtype recruitment, we calculated the number of CaMPARI2-seq cells assigned to each reference cluster within each condition and biological replicate (n=2). For each cluster *k*, enrichment relative to the E3 control was defined as log2(p_odor,k / p_E3,k), where p_odor,k and p_E3,k represent the proportions of cells assigned to cluster *k* in the odor and E3 conditions, respectively, within the same biological replicate.

To stabilize effect-size estimates for clusters with low cell counts, a small pseudocount (0.5) was added to cluster counts prior to calculating enrichment ratios. Pseudocounts were used only for calculating enrichment effect sizes and were not used in statistical testing.

Statistical significance of over-representation was assessed independently for each replicate and cluster using one-sided Fisher’s exact tests (alternative = “greater”), comparing counts assigned to cluster *k* versus all other clusters between odor and E3 conditions. Resulting p-values were corrected for multiple testing using the Benjamini–Hochberg false discovery rate procedure within each replicate.

To examine odor-specific recruitment independent of E3 baseline, pairwise comparisons between odor conditions were performed using Fisher’s exact test within each biological replicate. For each cluster, a 2×2 contingency table was constructed from the cell counts of two odor conditions, and enrichment was quantified as the log_2_ odds ratio with a Haldane-Anscombe correction (pseudocount = 0.5) for numerical stability. P-values were adjusted for multiple comparisons across clusters within each replicate using the Benjamini-Hochberg procedure, and log_2_ odds ratios were averaged across replicates to obtain summary statistics. To test whether valence-associated differences in subtype recruitment could arise by chance, we performed a permutation analysis in which odor identities (TCA, cytidine, ATP) were randomly reassigned across cells within each biological replicate, while preserving cluster identity, replicate structure, total cell counts per odor condition, and the E3 control population unchanged. For each of 5,000 permutations, enrichment relative to E3 was recomputed and a valence contrast statistic was defined per cluster as the difference between mean log₂ enrichment across attractive odors (TCA, cytidine) and the aversive odor (ATP). Directional and two-sided empirical p-values were calculated using the continuity-corrected formula (number of permutations meeting or exceeding the observed statistic + 1) / (total permutations + 1), and corrected across clusters using the Benjamini-Hochberg procedure.

#### Hybridization Chain Reaction (HCR) Labeling of Odor-Responsive Neurons

To link odor-evoked neuronal activity to gene expression, we combined CaMPARI2-based functional labeling with multiplexed HCR fluorescence in situ hybridization. Transgenic Tg(*HuC:CaMPARI2*) larvae were exposed to odor stimuli and photoconverted following the same protocol as for CaMPARI2 imaging and CaMPARI2-seq experiments (see above). After photoconversion and imaging, individual larvae were transferred to separate wells of a 48-well plate to preserve sample identity throughout fixation, staining, and imaging, following previously described workflow ^148^.

Larvae were fixed overnight at 4 °C in 4% paraformaldehyde (PFA) prepared in PBS-T (1× PBS, 0.1% Triton-X-100) with gentle agitation. The following day, samples were washed 3 × 5 min in PBS-T to terminate fixation and permeabilized in cold methanol (50% MeOH in PBS-T for 5 min, then 100% MeOH for ≥ 2 h at − 20 °C). Rehydration was carried out stepwise by first incubating in 50% MeOH for 5 min followed by 3 × 5 min washes in PBS-T.

Pre-hybridization was performed by incubating each larva in 250 µL of pre-warmed HCR3.0 Hybridization Buffer (Molecular Instruments) for 30 min at 37 °C. Samples were then hybridized overnight at 37 °C in 250 µL of the same buffer containing 2 pmol of primary probe per target gene. The next day, probes were removed by three 20-min washes at 37 °C in pre-warmed HCR Wash Buffer (Molecular Instruments), followed by three 5-min washes in 5× SSCT (5× SSC + 0.1% Tween-20) at room temperature.

For signal amplification, larvae were first incubated for 30 min in 250 µL of HCR Amplification Buffer (Molecular Instruments). Separately, hairpins h1 and h2 (30 pmol each) were snap-cooled by heating 3 µM stock solutions at 95 °C for 90 s and cooling to room temperature in the dark for 30 min. The hairpins were combined in 100 µL of amplification buffer with a final dilution of 1:50 supplemented with 1:1000 1mg/ml DAPI stock (Sigma #D9542) and added to each well for overnight incubation at room temperature in the dark. Excess hairpins were removed the following day by three 20-min washes in 5× SSCT containing 1:1000 DAPI (1 mg/mL). Samples were then imaged directly using a Yokogawa spinSR CSU-W1 spinning disk confocal microscope equipped with a 30×/1.05 NA silicon immersion objective. Z-stacks were acquired at 1μm steps with excitation at 488nm (200ms dwell) and 561nm (500ms dwell). Images were captured at 0.43μm XY resolution.

Live CaMPARI2 images were registered to the corresponding fixed HCR images using the green CaMPARI2 signal as a reference channel. Registration was performed with ANTs using the following command:

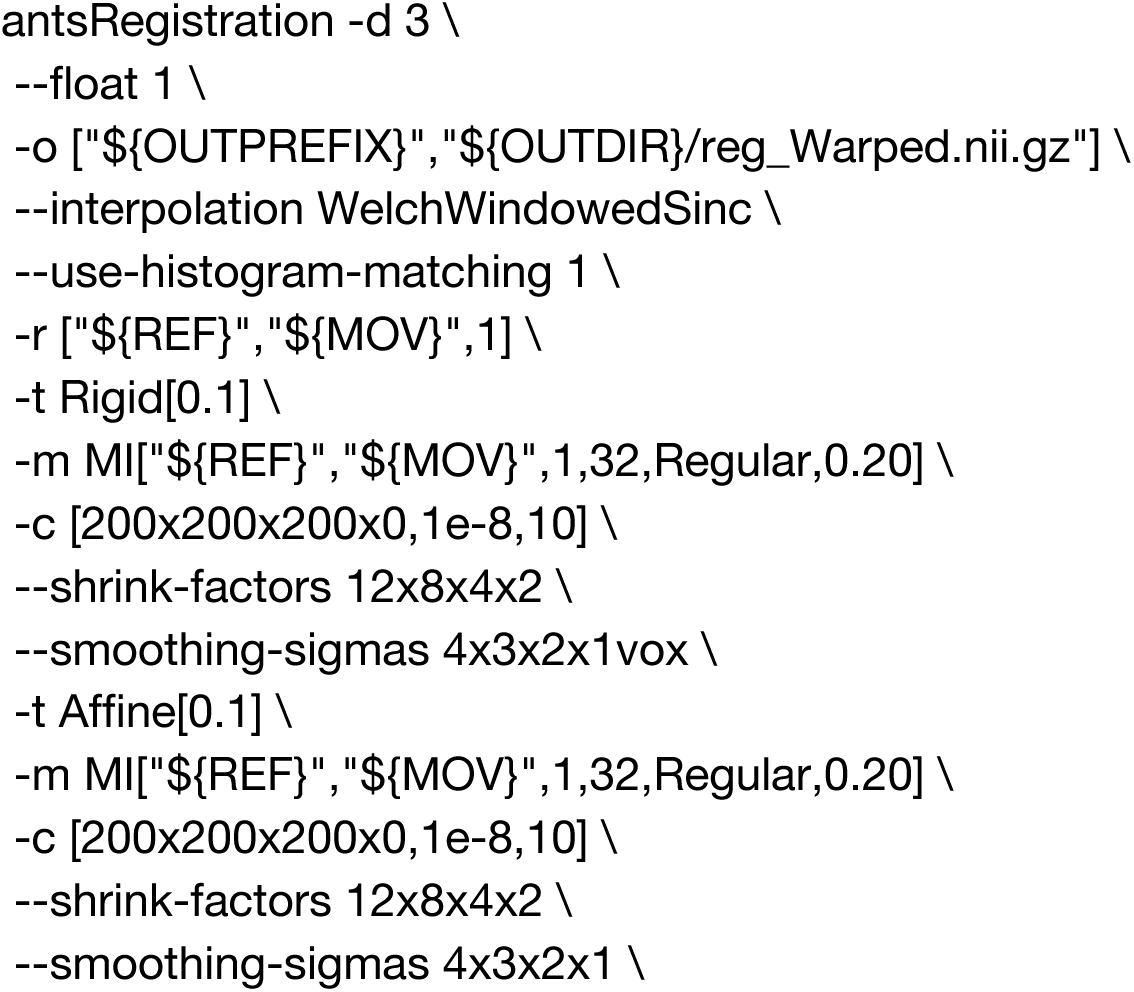
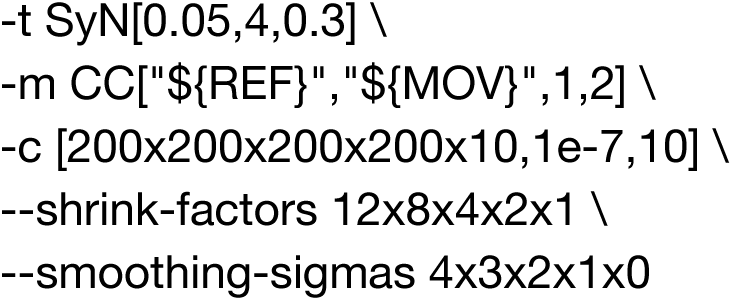

After registration, HCR-positive OB neurons were identified in 3D image stacks, and for each brain we quantified the fraction of HCR⁺ cells exhibiting elevated CaMPARI2 red-to-green (R/G) ratios, normalized per brain. This fraction served as a measure of odor-evoked recruitment for each gene and odor condition. Statistical comparisons across odors were performed using nonparametric tests, with an omnibus Kruskal–Wallis test followed by two-sided Mann–Whitney U tests for pairwise comparisons, applying Benjamini–Hochberg correction within each panel to control the false discovery rate.

#### Two-Photon Laser Ablation of TH⁺ Olfactory Bulb Interneurons

Targeted ablation of dopaminergic interneurons in the OB was performed using two-photon laser irradiation. We used heterozygous transgenic larvae generated by outcrossing th:QF2>GFP fish to a nacre^-/-^ line, resulting in GFP labeling of tyrosine hydroxylase–expressing (TH^+^) OB interneurons.

Larvae were screened for GFP expression at 24 hours post-fertilization (hpf). At 4 dpf, selected larvae were anesthetized in E3 medium containing tricaine methanesulfonate (MS-222) and mounted dorsal-side up in 2% low-melting point agarose. To minimize scattering, the agarose covering the dorsal region of the forebrain was carefully removed. Larvae were imaged using a Scientifica system equipped with a Nikon 16×/0.8 NA water-dipping objective, controlled by ScanImage v2023.1.1 ^159^ with the Photostimulation module.

GFP-expressing neurons were visualized using a fixed-wavelength 920 nm two-photon excitation laser (FemtoFiber Ultra 920 with AOM-Toptica). Ablation targets were restricted to the OB and selected manually based on GFP expression and soma morphology. Targeted cells were illuminated using a point-stimulation architecture at maximal laser power (290 mW at 800nm) from a Mai Tai multiphoton laser (Spectra-Physics). Each target received six sequential repetitions of 1 ms pulses. Sham controls were processed in parallel using identical mounting and targeting procedures but were irradiated at a subthreshold laser power (60 mW at 800nm), insufficient to induce detectable photodamage or GFP bleaching.

Following ablation or sham irradiation, larvae were gently freed from the agarose and transferred to fresh E3 medium for overnight recovery at 28°C. Survival was 100%, and no overt morphological damage was observed in recovered animals.

#### Behavioral Assays Following Ablation

At 5 dpf, recovered larvae were tested individually in the two-choice olfactory preference arena described above. To assess the integrity of general olfactory function post-ablation, each larva underwent multiple independent trials using distinct odorants. Each trial consisted of a 10-minute acclimatization period with E3 flowing from both inlets, followed by a 2-minute odor exposure epoch (side randomized across trials), and a washout period. Between trials, larvae were transferred to individual wells containing fresh E3 and allowed to rest for approximately 1 hour. We further required active arena sampling during odor presentation: trials were included only if larvae visited both sides of the arena during the odor epoch, and trials exhibiting clear tracking or recording artifacts were excluded from downstream preference analyses.

To control for habituation or trial-order effects, pilot experiments confirmed that locomotion parameters (speed, total distance) and odor preference remained stable across repeated introductions to the arena (*P* = 0.19, 0.12 and *P* = 0.622, respectively; paired tests, *n* = 12).

#### Post-hoc Imaging and Quantification

To verify the extent of photoablation, larvae were anesthetized following behavioral testing and fixed overnight at 4°C in 4% PFA in PBS-T (1× PBS, 0.1% Triton-X-100) with gentle agitation. Samples were washed (3 × 5 min) in PBS-T and stained overnight with DAPI (1 mg/mL, 1:1000; Sigma #D9542). Whole-OB volumes were imaged using a Yokogawa spinSR CSU-W1 spinning disk confocal microscope equipped with a 30×/1.05 NA silicon immersion objective. Z-stacks were acquired at 1 μm intervals with excitation at 488 nm (200 ms dwell) and 405 nm (50 ms dwell) at a 0.43 μm XY resolution. Photoablation efficacy was quantified by GFP⁺ cell counts in the olfactory bulb, revealing a ∼50% reduction in total GFP^+^ somata in 2p-ablated larvae relative to sham and control animals (median totals: 50 vs. 105 cells), with a small left–right difference detected in the ablated group.

## Resource Availability

### Lead Contact

Requests for further information and resources should be directed to and will be fulfilled by the corresponding authors.

### Materials Availability

Lines and reagents generated in this study are available from the corresponding authors upon request and with a completed materials transfer agreement.

### Data and Code Availability

All analysis scripts are available through the github repository: https://github.com/Mayseless/zebrafish-olfactory-bulb-odor-preference. Single cell and CaMPARI2-seq data, as well as behavior data and CaMPARI2 odor maps will be available upon publication (10.5281/zenodo.19607911).

