## Supplementary material for "Odor preference maps to cohesive transcriptional domains in the olfactory bulb": 265 Adapter Power Board.pdf

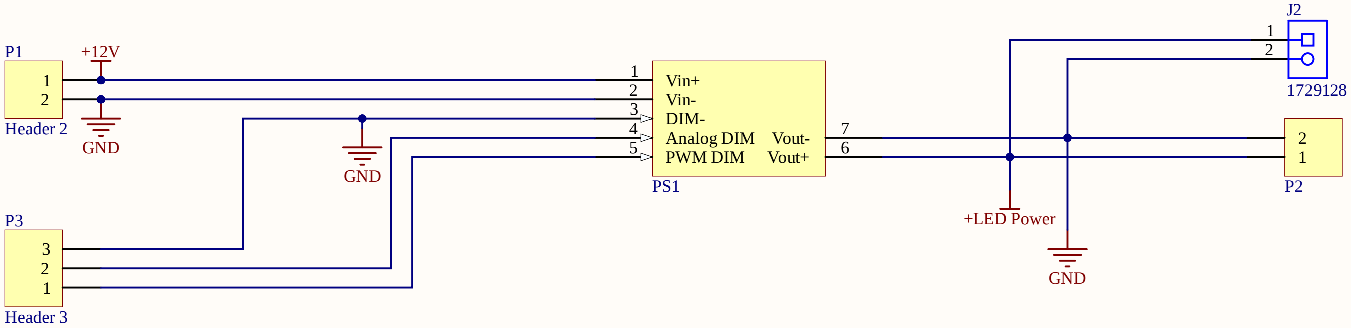


Title *265 Adapter Power Board*

Size: A4 Number: 1 Revision: 1.0

Date: 26.01.2024 Time: 14:28:00 Sheet1 of 1

File: U:\SG Electronics Workshop\Group\Projekte\ELWE PROJECTS\265 Valve Led Driver Oded Group Schier\Altium\265 Adapter Power Board\265 Adapter Power Board\265 Adapter

*Biozentrum Uni Basel*
*Zentrale Elektronikwekstatt*
*Spitalstrasse 41 CH-4056 Basel*

BIOZENTRUM
Universität Basel
The Center for
Molecular Life Sciences

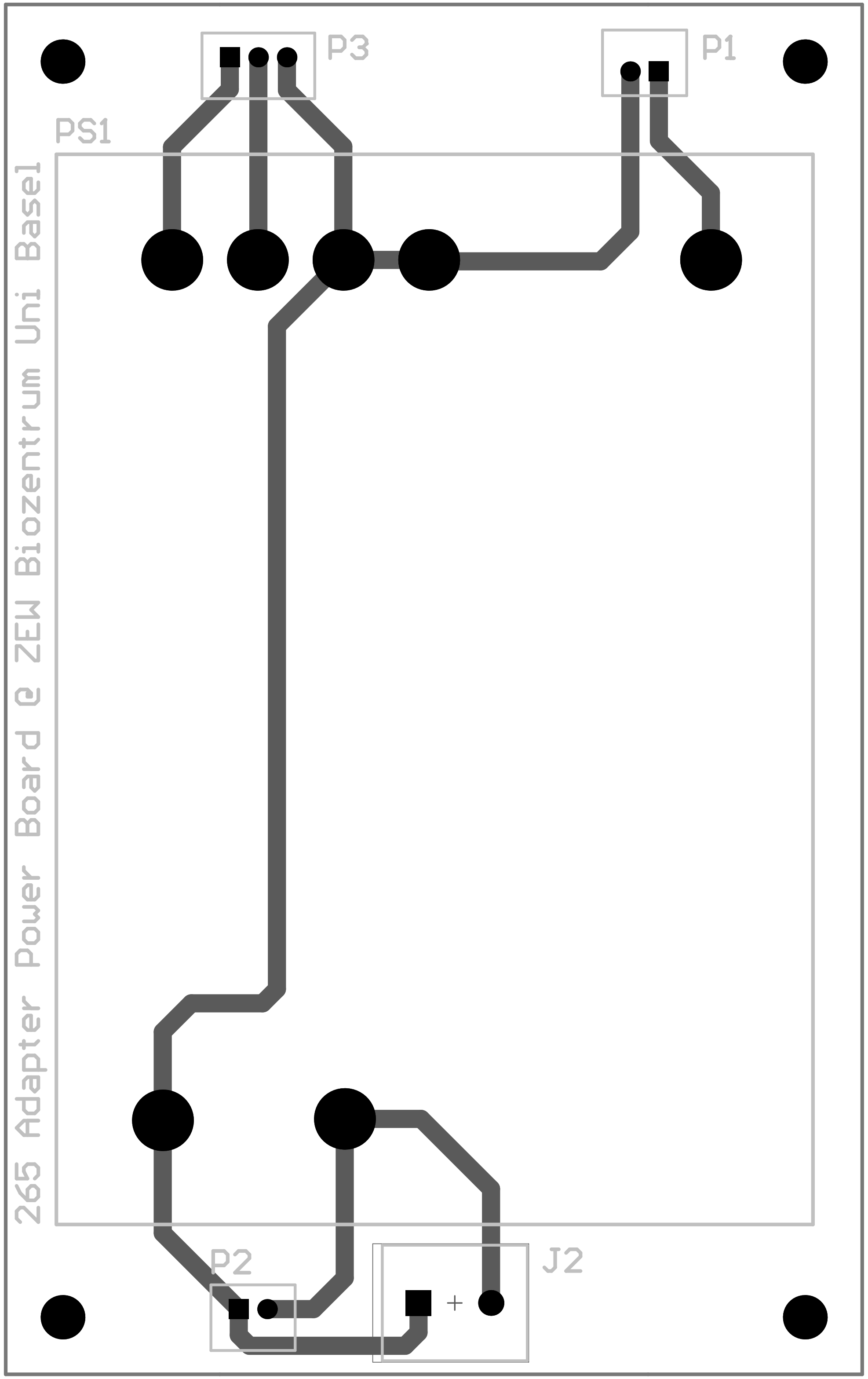
P3
P1
PS1
265 Adapter Power Board @ ZEW Biozentrum Uni Basel
P2
J2
+
