## Supplementary material for "Odor preference maps to cohesive transcriptional domains in the olfactory bulb": 265 LED Board Multi Channel Oded.pdf

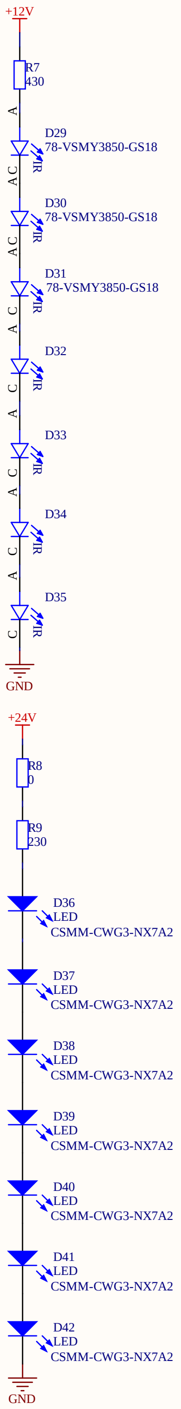

| | | |
|---|---|---|
| Title **_LED Board Multi Channel Oded_** | | |
| Size: A3 | Number: 1 | Revision: 1.0 |
| Date: 26.01.2024 Time: 14:22:35 | | Sheet1 of 16 |
| File: U:\SG Electronics Workshop\Group\Projekte\ELWE PROJECTS\265 Valve Led Driver Oded Group Schier\Altium\265 LED Board\265 LED Board Multi CHannel Oded\LED Strip_2.Sch | | |

*Biozentrum Universität Basel*
*Zentrale Elektronikwekstatt*
*Spitalstrasse 41 CH-4056 Basel*

BIOZENTRUM
Universität Basel
The Center for
Molecular Life Sciences

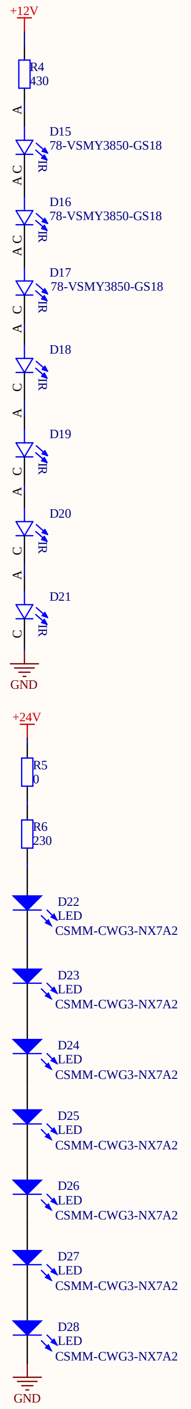

| Title | | | |
|---|---|---|---|
| Size: A3 | Number: 1 | Revision: 1.0 | |
| Date: 26.01.2024 | Time: 14:22:36 | Sheet2 | of 16 |
| File: U:\SG Electronics Workshop\Group\Projekte\ELWE PROJECTS\265 Valve Led Driver Oded Group Schier\Altium\265 LED Board\265 LED Board Multi CHannel Oded\LED Strip_3.Sch | | | |

*Biozentrum Universität Basel*
*Zentrale Elektronikwekstatt*
*Klingelbergstrasse 70 CH-4056 Basel*
BIOZENTRUM
Universität Basel
The Center for
Molecular Life Sciences

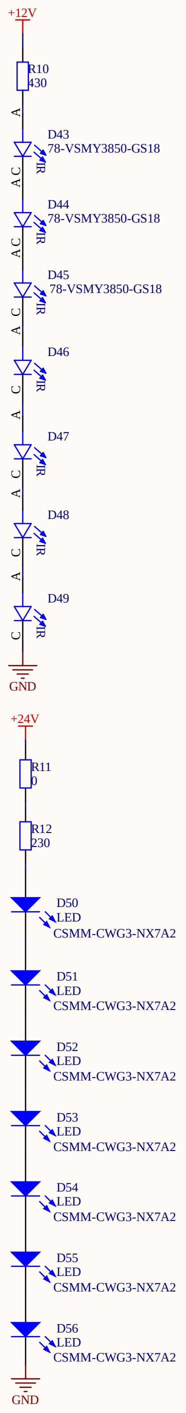

| | | | |
|---|---|---|---|
| Title | | | *Biozentrum Universität Basel*<br>*Zentrale Elektronikwekstatt*<br>*Klingelbergstrasse 70 CH-4056 Basel*<br>*Simon Saner Tel:+41 61 267 04 75*<br>*Serafin Wick Tel: +41 61 267 20 87* |
| Size: A3 | Number: 1 | Revision: 1.0 | |
| Date: 26.01.2024 Time: 14:22:36 | | Sheet3 of 16 | |

BIOZENTRUM
Universität Basel
The Center for
Molecular Life Sciences

File: U:\SG Electronics Workshop\Group\Projekte\ELWE PROJECTS\265 Valve Led Driver Oded Group Schier\Altium\265 LED Board\265 LED Board Multi CHannel Oded\LED Strip_4.Sc

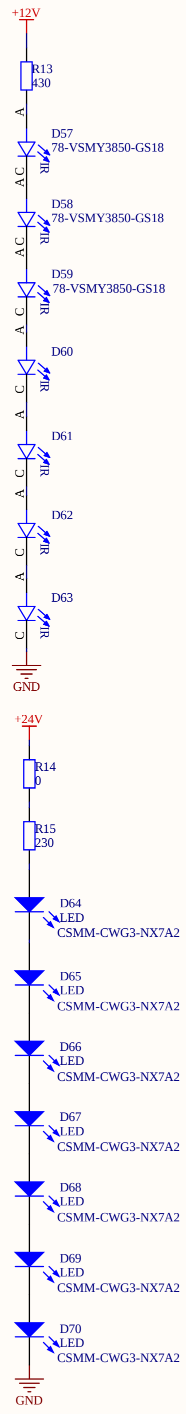

| Title | | |
|---|---|---|
| Size: A3 | Number: 1 | Revision: 1.0 |
| Date: 26.01.2024 Time: 14:22:36 | | Sheet4 of 16 |
| File: U:\SG Electronics Workshop\Group\Projekte\ELWE PROJECTS\265 Valve Led Driver Oded Group Schier\Altium\265 LED Board\265 LED Board Multi CHannel Oded\LED Strip_5.Sch | | |

*Biozentrum Universität Basel*
*Zentrale Elektronikwekstatt*
*Klingelbergstrasse 70 CH-4056 Basel*

BIOZENTRUM
Universität Basel
The Center for
Molecular Life Sciences

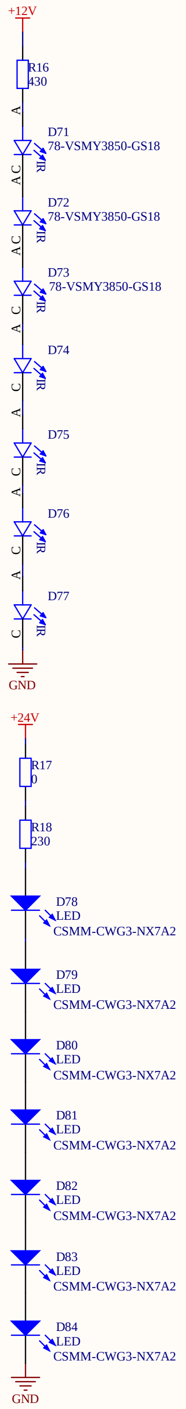

| | | |
|---|---|---|
| Title | | |
| Size: A3 | Number: 1 | Revision: 1.0 |
| Date: 26.01.2024 Time: 14:22:36 | | Sheet5 of 16 |

File: U:\SG Electronics Workshop\Group\Projekte\ELWE PROJECTS\265 Valve Led Driver Oded Group Schier\Altium\265 LED Board\265 LED Board Multi CHannel Oded\LED Strip_6.Sch

*Biozentrum Universität Basel*
*Zentrale Elektronikwekstatt*
*Klingelbergstrasse 70 CH-4056 Basel*
BIOZENTRUM
Universität Basel
The Center for
Molecular Life Sciences

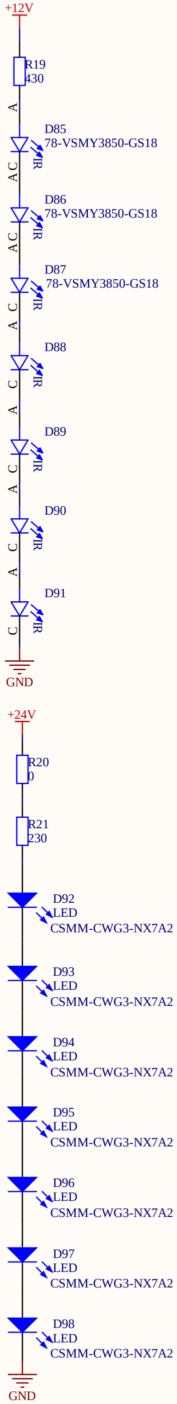

| Title | | | |
|---|---|---|---|
| Size: A3 | Number: 1 | Revision: 1.0 | |
| Date: 26.01.2024 | Time: 14:22:37 | Sheet6 | of 16 |
| File: U:\SG Electronics Workshop\Group\Projekte\ELWE PROJECTS\265 Valve Led Driver Oded Group Schier\Altium\265 LED Board\265 LED Board Multi CHannel Oded\LED Strip_7.Sch | | | |

*Biozentrum Universität Basel*
*Zentrale Elektronikwekstatt*
*Klingelbergstrasse 70 CH-4056 Basel*

BIOZENTRUM
Universität Basel
The Center for
Molecular Life Sciences

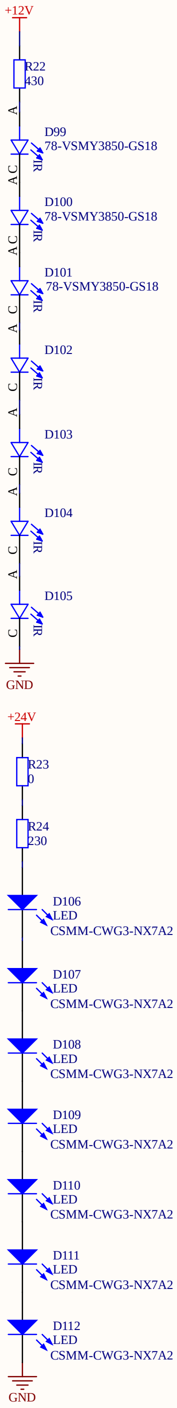

| | |
|---|---|
| Title | |
| Size: A3 | Number: 1 | Revision: 1.0 |
| Date: 26.01.2024 Time: 14:22:37 | Sheet7 of 16 |
| File: U:\SG Electronics Workshop\Group\Projekte\ELWE PROJECTS\265 Valve Led Driver Oded Group Schier\Altium\265 LED Board\265 LED Board Multi CHannel Oded\LED Strip_8.Sc[illegible] | |

*Biozentrum Universität Basel*
*Zentrale Elektronikwekstatt*
*Klingelbergstrasse 70 CH-4056 Basel*

BIOZENTRUM
Universität Basel
The Center for
Molecular Life Sciences

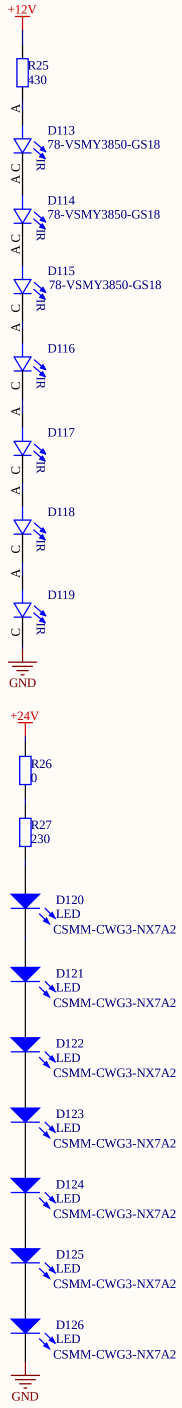

| | | | |
|---|---|---|---|
| Title | | | *Biozentrum Universität Basel*<br>*Zentrale Elektronikwekstatt*<br>*Klingelbergstrasse 70 CH-4056 Basel*<br>*Simon Saner Tel:+41 61 267 04 75*<br>*Serafin Wick Tel: +41 61 267 20 87* |
| Size: A3 | Number: 1 | Revision: 1.0 | |
| Date: 26.01.2024 Time: 14:22:37 | | Sheet8 of 16 | |
| File: U:\SG Electronics Workshop\Group\Projekte\ELWE PROJECTS\265 Valve Led Driver Oded Group Schier\Altium\265 LED Board\265 LED Board Multi CHannel Oded\LED Strip_9.Sc[illegible] | | | |

BIOZENTRUM
Universität Basel
The Center for
Molecular Life Sciences

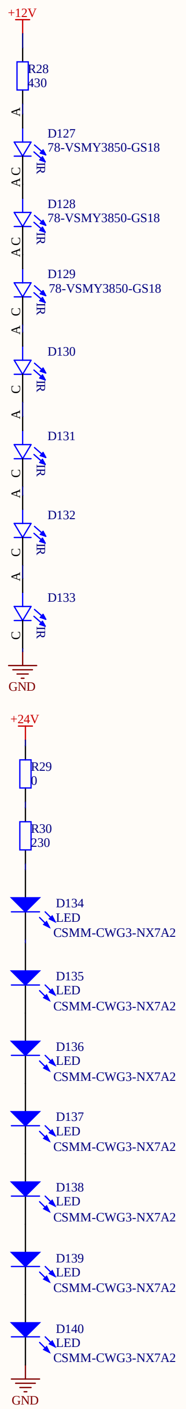

| Title | | |
|---|---|---|
| Size: A3 | Number: 1 | Revision: 1.0 |
| Date: 26.01.2024 Time: 14:22:37 | | Sheet9 of 16 |
| File: U:\SG Electronics Workshop\Group\Projekte\ELWE PROJECTS\265 Valve Led Driver Oded Group Schier\Altium\265 LED Board\265 LED Board Multi CHannel Oded\LED Strip_10.Sc | | |

*Biozentrum Universität Basel*
*Zentrale Elektronikwekstatt*
*Klingelbergstrasse 70 CH-4056 Basel*

BIOZENTRUM
Universität Basel
The Center for
Molecular Life Sciences

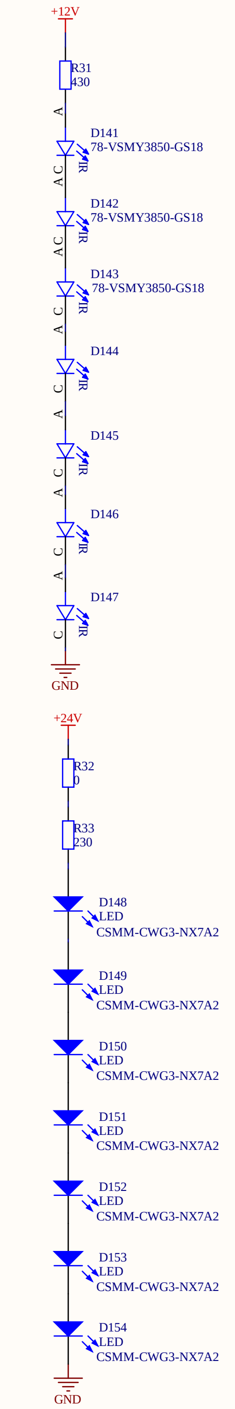

| Title | | |
|---|---|---|
| Size: A3 | Number: 1 | Revision: 1.0 |
| Date: 26.01.2024 | Time: 14:22:37 | Sheet 10 of 16 |

File: U:\SG Electronics Workshop\Group\Projekte\ELWE PROJECTS\265 Valve Led Driver Oded Group Schier\Altium\265 LED Board\265 LED Board Multi CHannel Oded\LED Strip_11.Sc

*Biozentrum Universität Basel*
*Zentrale Elektronikwekstatt*
*Klingelbergstrasse 70 CH-4056 Basel*

BIOZENTRUM
Universität Basel
The Center for
Molecular Life Sciences

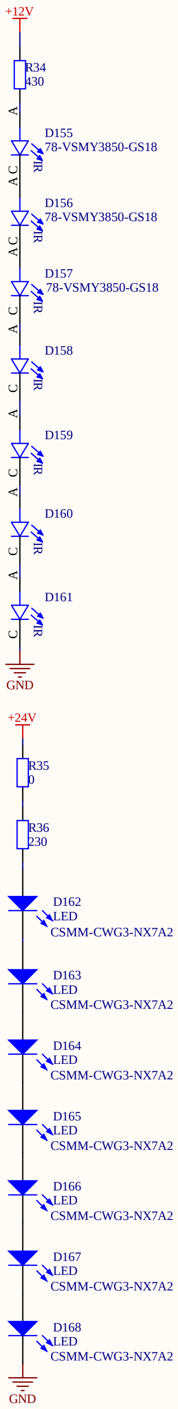

| | | |
|---|---|---|
| Title | | |
| Size: A3 | Number: 1 | Revision: 1.0 |
| Date: 26.01.2024 Time: 14:22:38 | | Sheet 11 of 16 |

File: U:\SG Electronics Workshop\Group\Projekte\ELWE PROJECTS\265 Valve Led Driver Oded Group Schier\Altium\265 LED Board\265 LED Board Multi CHannel Oded\LED Strip_12.Sc

*Biozentrum Universität Basel*
*Zentrale Elektronikwekstatt*
*Klingelbergstrasse 70 CH-4056 Basel*

BIOZENTRUM
Universität Basel
The Center for
Molecular Life Sciences

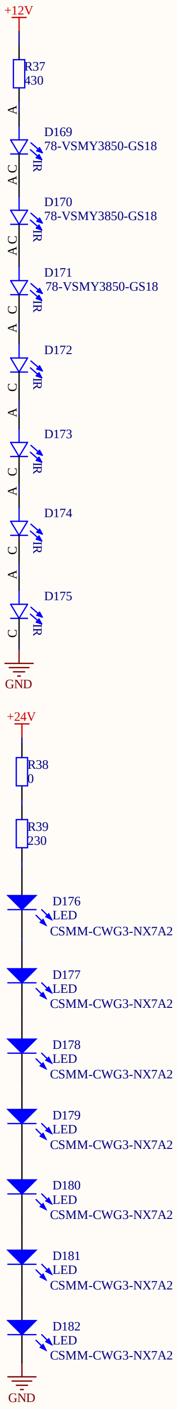

| | | | |
|---|---|---|---|
| Title | | | *Biozentrum Universität Basel* <br> *Zentrale Elektronikwekstatt* <br> *Klingelbergstrasse 70 CH-4056 Basel* <br> *Simon Saner Tel:+41 61 267 04 75* <br> *Serafin Wick Tel: +41 61 267 20 87* |
| Size: A3 | Number: 1 | Revision:1.0 | |
| Date: 26.01.2024 Time: 14:22:38 | | Sheet12 of 16 | |
| File: U:\SG Electronics Workshop\Group\Projekte\ELWE PROJECTS\265 Valve Led Driver Oded Group Schier\Altium\265 LED Board\265 LED Board Multi CHannel Oded\LED Strip_13.S[illegible] | | | |

BIOZENTRUM
Universität Basel
The Center for
Molecular Life Sciences

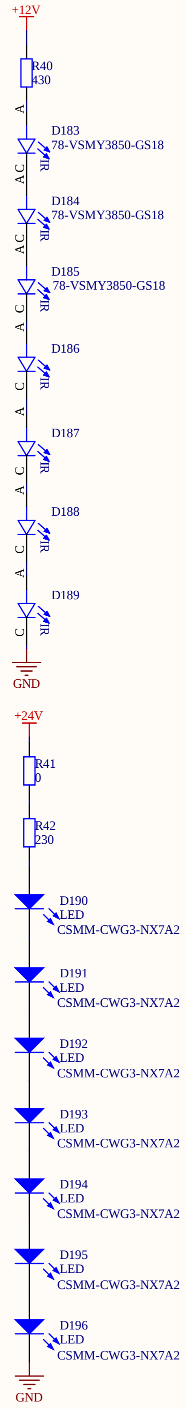

| Title | | |
|---|---|---|
| Size: A3 | Number: 1 | Revision: 1.0 |
| Date: 26.01.2024 | Time: 14:22:38 | Sheet 13 of 16 |
| File: U:\SG Electronics Workshop\Group\Projekte\ELWE PROJECTS\265 Valve Led Driver Oded Group Schier\Altium\265 LED Board\265 LED Board Multi CHannel Oded\LED Strip_14.Sc | | |

*Biozentrum Universität Basel*
*Zentrale Elektronikwekstatt*
*Klingelbergstrasse 70 CH-4056 Basel*

BIOZENTRUM
Universität Basel
The Center for
Molecular Life Sciences

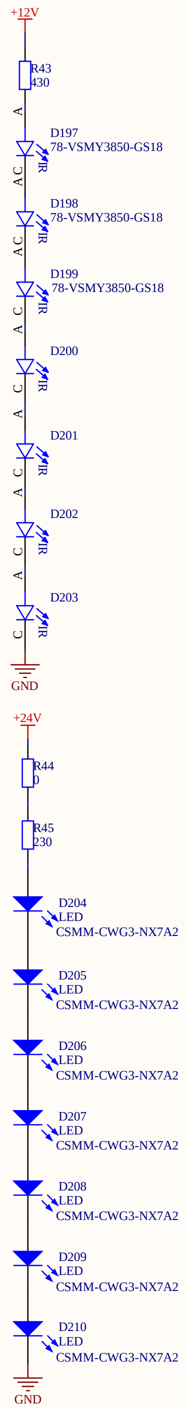

| Title | | |
|---|---|---|
| Size: A3 | Number: 1 | Revision: 1.0 |
| Date: 26.01.2024 | Time: 14:22:38 | Sheet 14 of 16 |
| File: U:\SG Electronics Workshop\Group\Projekte\ELWE PROJECTS\265 Valve Led Driver Oded Group Schier\Altium\265 LED Board\265 LED Board Multi CHannel Oded\LED Strip_15.Sc | | |

*Biozentrum Universität Basel*
*Zentrale Elektronikwekstatt*
*Klingelbergstrasse 70 CH-4056 Basel*

BIOZENTRUM
Universität Basel
The Center for
Molecular Life Sciences

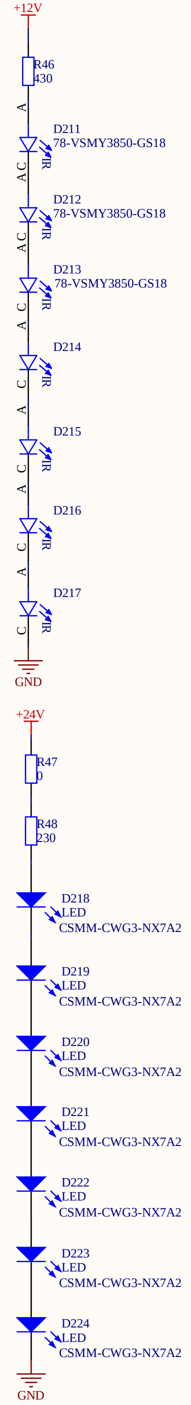

| Title | | | |
|---|---|---|---|
| Size: A3 | Number: 1 | Revision: 1.0 | |
| Date: 26.01.2024 | Time: 14:22:38 | Sheet 15 | of 16 |
| File: U:\SG Electronics Workshop\Group\Projekte\ELWE PROJECTS\265 Valve Led Driver Oded Group Schier\Altium\265 LED Board\265 LED Board Multi CHannel Oded\LED Strip_16.Sc | | | |

*Biozentrum Universität Basel*
*Zentrale Elektronikwekstatt*
*Klingelbergstrasse 70 CH-4056 Basel*

BIOZENTRUM
Universität Basel
The Center for
Molecular Life Sciences

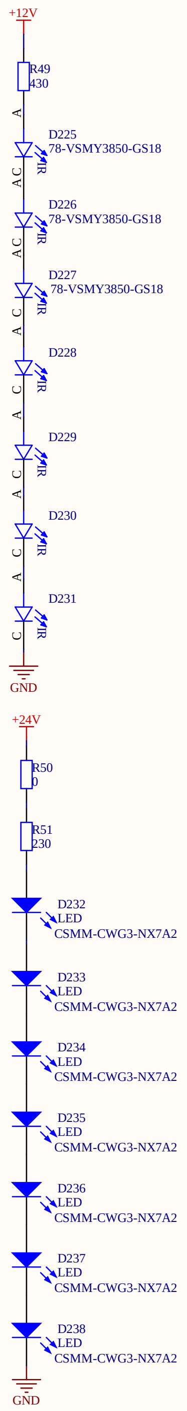

| | | |
|---|---|---|
| Title | | |
| Size: A3 | Number: 1 | Revision: 1.0 |
| Date: 26.01.2024 Time: 14:22:38 | | Sheet 16 of 16 |
| File: U:\SG Electronics Workshop\Group\Projekte\ELWE PROJECTS\265 Valve Led Driver Oded Group Schier\Altium\265 LED Board\265 LED Board Multi CHannel Oded\LED Strip_17.Sc | | |

*Biozentrum Universität Basel*
*Zentrale Elektronikwekstatt*
*Klingelbergstrasse 70 CH-4056 Basel*

BIOZENTRUM
Universität Basel
The Center for
Molecular Life Sciences

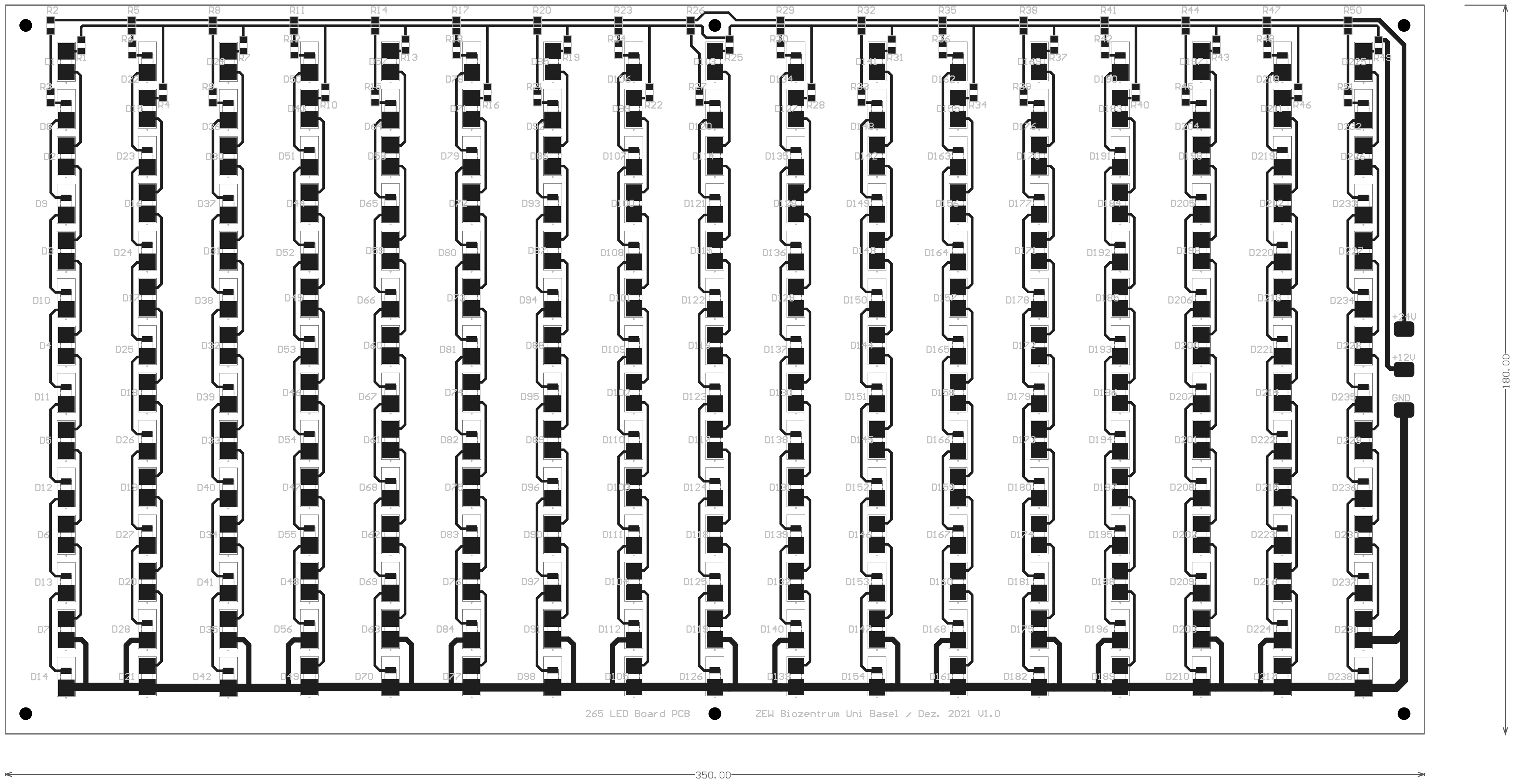
265 LED Board PCB
ZEW Biozentrum Uni Basel / Dez. 2021 V1.0
+24V
+12V
GND
350.00
180.00
