## Supplementary material for "Odor preference maps to cohesive transcriptional domains in the olfactory bulb": UV_405nm.pdf

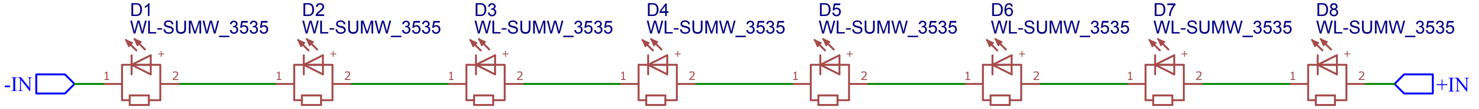


8x 15335340AA350
https://www.mouser.ch/ProductDetail/Wurth-Elektronik/15335340AA350?qs=sGAEpiMZZMusoohG2hS%252B107%252BEN%252B%252BjBF0nIqSWF4w5JNymEhdRVtDug%3D%3D

2 PCBs für ein komplettes Setup

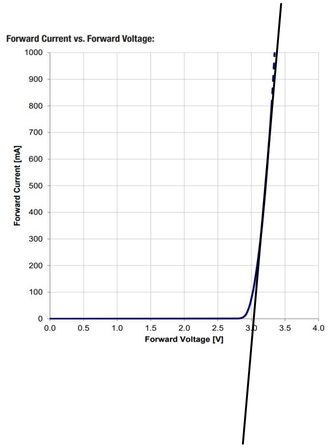


I_f aus experiment: 700mA passt.
U_f aus Datenblatt, ca.3.5V *8 LEDs= 28V
P=19.6W

Für die Verwendung von zwei LED-Boards (in Serie) eignet sich Meanwell LDH-65-1050W

P_total = 40 W
I_total = 700mA
U_total = 56V

LED Regler kann Spannung erhöhen, aus Versuch:
24 V Eingang mit 1.73A (=41.5 W)
Ausgang: 56V, 700mA (Stromgeregelt) mit U_DIM=5.5V

R0=dU/dI=0.25V/A = 0.25Ohm

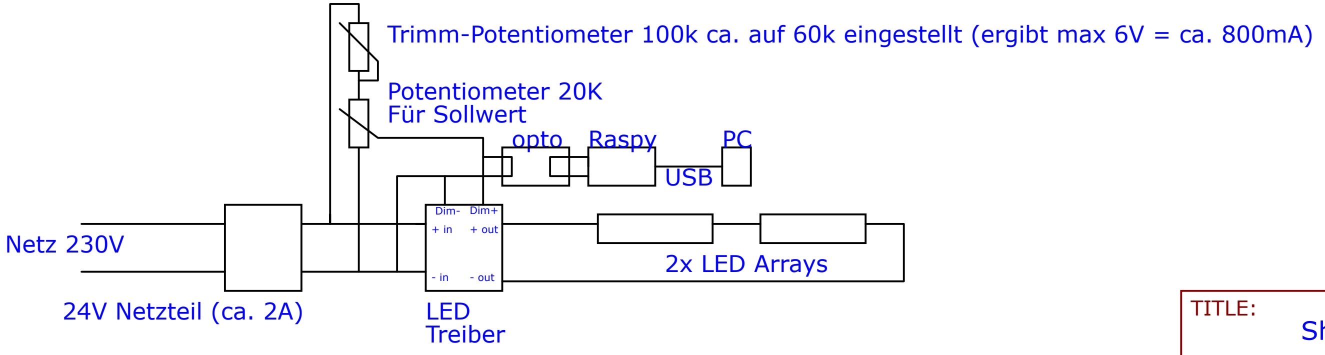


| | | |
|---|---|---|
| TITLE: Sheet_1 | | REV: 1.0 |
| EasyEDA | Company: Biozentrum Uni Basel ZEW | Sheet: 1/1 |
| | Date: 2023-10-12 Drawn By: christianoxe | |
