## Supplementary material for "Odor preference maps to cohesive transcriptional domains in the olfactory bulb": valve_driver_V1.ino

```
/*
 * 13.07.2022 Biozentrum der Universität Basel, Christian Oxé
 * 
 * Simple command.handler for a LabVIEW Interface
 * Controlls 16 Valves and a PWM LED according to a String from the host PC
 * Format for Serial communication:
 * sxxxxxxxxxxxxxxxxyye
 * s and e are fix flags to show start and end of message.
 * xx...   are the single bits for each valve, starting with 1
 * yy...   is the %-value for the LED (visible)
 * x can be 0 or 1 (0=valve closed, 1=valve open) any other value is seen as 0
 * yy must be a 2-digit String, (numbers belwo 10 must start with 0) 0...99
 */


String data;                                  // Stores the Data from Serial as String

void setup() {
  Serial.begin(9600);                         // start serial port at 9600 boud

// Block 1

pinMode(5,OUTPUT);
pinMode(4,OUTPUT);
pinMode(3,OUTPUT);
pinMode(2,OUTPUT);
pinMode(24,OUTPUT);
pinMode(10,OUTPUT);
pinMode(9,OUTPUT);
pinMode(6,OUTPUT);

//Block 2

pinMode(28,OUTPUT);
pinMode(27,OUTPUT);
pinMode(26,OUTPUT);
pinMode(25,OUTPUT);
pinMode(32,OUTPUT);
pinMode(31,OUTPUT);
pinMode(30,OUTPUT);
pinMode(29,OUTPUT);
  
}

void loop() {

delay(10);                                   // Slow down the programm

  if (Serial.available()){
    data="";                                  //if new data available, first delete the existing data
  }
  
  while (Serial.available()){
    data+=(char(Serial.read()));              //write all new data in the String "data"
  }

   Serial.println(data);                      // for debugging: returns the new data

  if(data.length()>15 && data.charAt(0) == 's' && data.charAt(19) == 'e'){     // Check if Data follws the rule
    Serial.println("valid data");               // for debugging

    // Block 1

    if(data.charAt(1) == '1'){
      digitalWrite(5,HIGH);
    }else{
      digitalWrite(5,LOW);
    }
    
    if(data.charAt(2) == '1'){
      digitalWrite(4,HIGH);
    }else{
      digitalWrite(4,LOW);
    }
    if(data.charAt(3) == '1'){
      digitalWrite(3,HIGH);
    }else{
      digitalWrite(3,LOW);
    }
    
    if(data.charAt(4) == '1'){
      digitalWrite(2,HIGH);
    }else{
      digitalWrite(2,LOW);
    }
    
    if(data.charAt(5) == '1'){
      digitalWrite(24,HIGH);
    }else{
      digitalWrite(24,LOW);
    }
    
    if(data.charAt(6) == '1'){
      digitalWrite(10,HIGH);
    }else{
      digitalWrite(10,LOW);
    }
    if(data.charAt(7) == '1'){
      digitalWrite(9,HIGH);
    }else{
      digitalWrite(9,LOW);
    }
    
    if(data.charAt(8) == '1'){
      digitalWrite(6,HIGH);
    }else{
      digitalWrite(6,LOW);
    }
    
    //Block 2

    if(data.charAt(9) == '1'){
      digitalWrite(28,HIGH);
    }else{
      digitalWrite(28,LOW);
    }
    
    if(data.charAt(10) == '1'){
      digitalWrite(27,HIGH);
    }else{
      digitalWrite(27,LOW);
    }
    if(data.charAt(11) == '1'){
      digitalWrite(26,HIGH);
    }else{
      digitalWrite(26,LOW);
    }
    
    if(data.charAt(12) == '1'){
      digitalWrite(25,HIGH);
    }else{
      digitalWrite(25,LOW);
    }
    if(data.charAt(13) == '1'){
      digitalWrite(32,HIGH);
    }else{
      digitalWrite(32,LOW);
    }
    
    if(data.charAt(14) == '1'){
      digitalWrite(31,HIGH);
    }else{
      digitalWrite(31,LOW);
    }
    if(data.charAt(15) == '1'){
      digitalWrite(30,HIGH);
    }else{
      digitalWrite(30,LOW);
    }
    
    if(data.charAt(16) == '1'){
      digitalWrite(29,HIGH);
    }else{
      digitalWrite(29,LOW);
    }

    String LED_value_str=data.substring(17,19); // get LED value from Input String
    int LED_value_int =LED_value_str.toInt();   // convert LED from String to integer
    //Serial.println(LED_value_str);          // for debugging
    //Serial.println(LED_value_int);          // for debuging
    analogWrite(35,LED_value_int*2.6);         // write Analog output (0...255)
    
  }else{
    Serial.println("wrong format");             // for debugging
  }
  


}
```
