## Supplementary material for "Odor preference maps to cohesive transcriptional domains in the olfactory bulb": 1003040.pdf

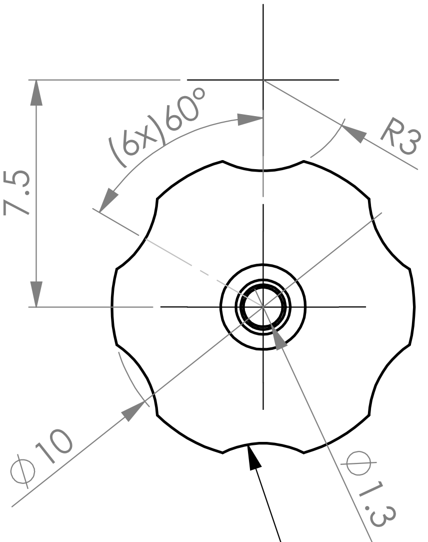


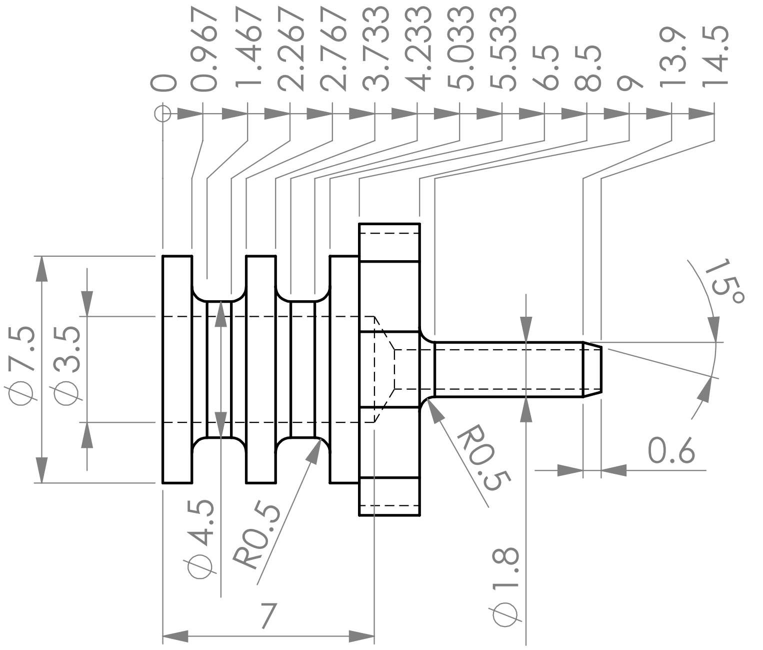


Alternativ auch gestossene Randrierung möglich!

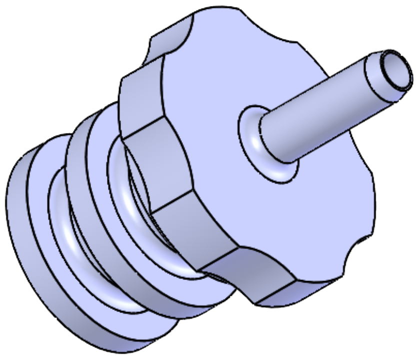

Alle Kanten gebrochen

| Stückzahl | Pos. | Gegenstand | Werkstoff | Bemerkung |
|---|---|---|---|---|
| 1 | 1 | | 1.4301 (X5CrNi18-10) | |

| | | |
|---|---|---|
| UNI BASEL | Massstab 4:1 | Gezeichnet: P. Schlenker |
| | | Datum: 11.01.2021 |
| | | Auftraggeber: |
| | | Blatt Anzahl: 1 von 1 |
| Biozentrum | Zentral Werkstatt | No. 1003040 |
