## Supplementary material for "Odor preference maps to cohesive transcriptional domains in the olfactory bulb": 1003051.pdf

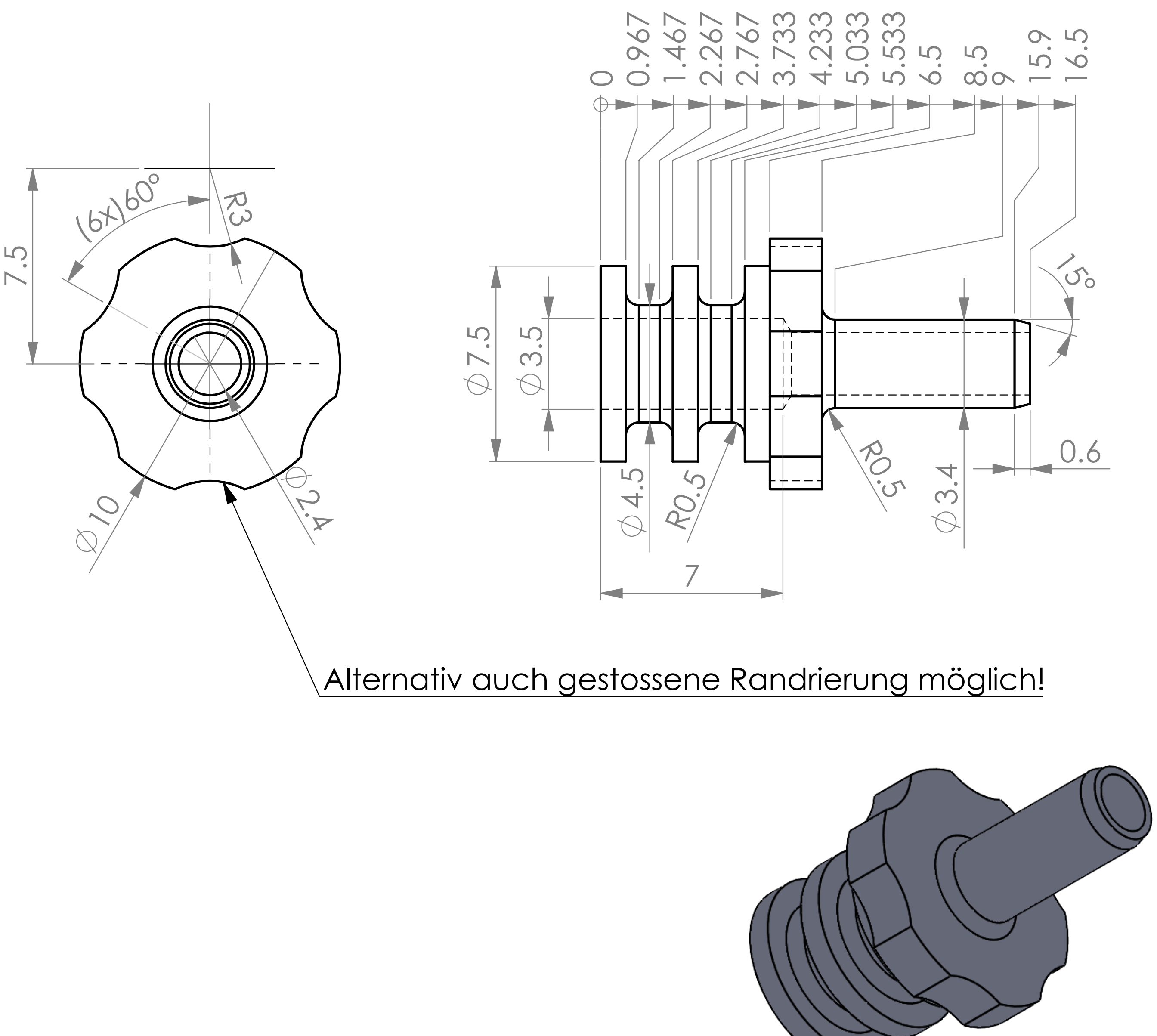


Alle Kanten gebrochen

| | | | | | |
|---|---|---|---|---|---|
| 1 | 1 | | 1.4301 (X5CrNi18-10) | | |
| Stückzahl | Pos. | Gegenstand | Werkstoff | Bemerkung | |
| UNI BASEL | | | Massstab 4:1 | Gezeichnet | P. Schlenker |
| | | | | Datum | 11.01.2021 |
| | | | | Auftraggeber | |
| | | | | Blatt Anzahl | 1 von 1 |
| | | Biozentrum | Zentral Werkstatt | No. | 1003051 |
