## Supplementary material for "Odor preference maps to cohesive transcriptional domains in the olfactory bulb": 1003259.pdf

Alle Aussenkanten entgratet
Alle Masse +/-0.1mm

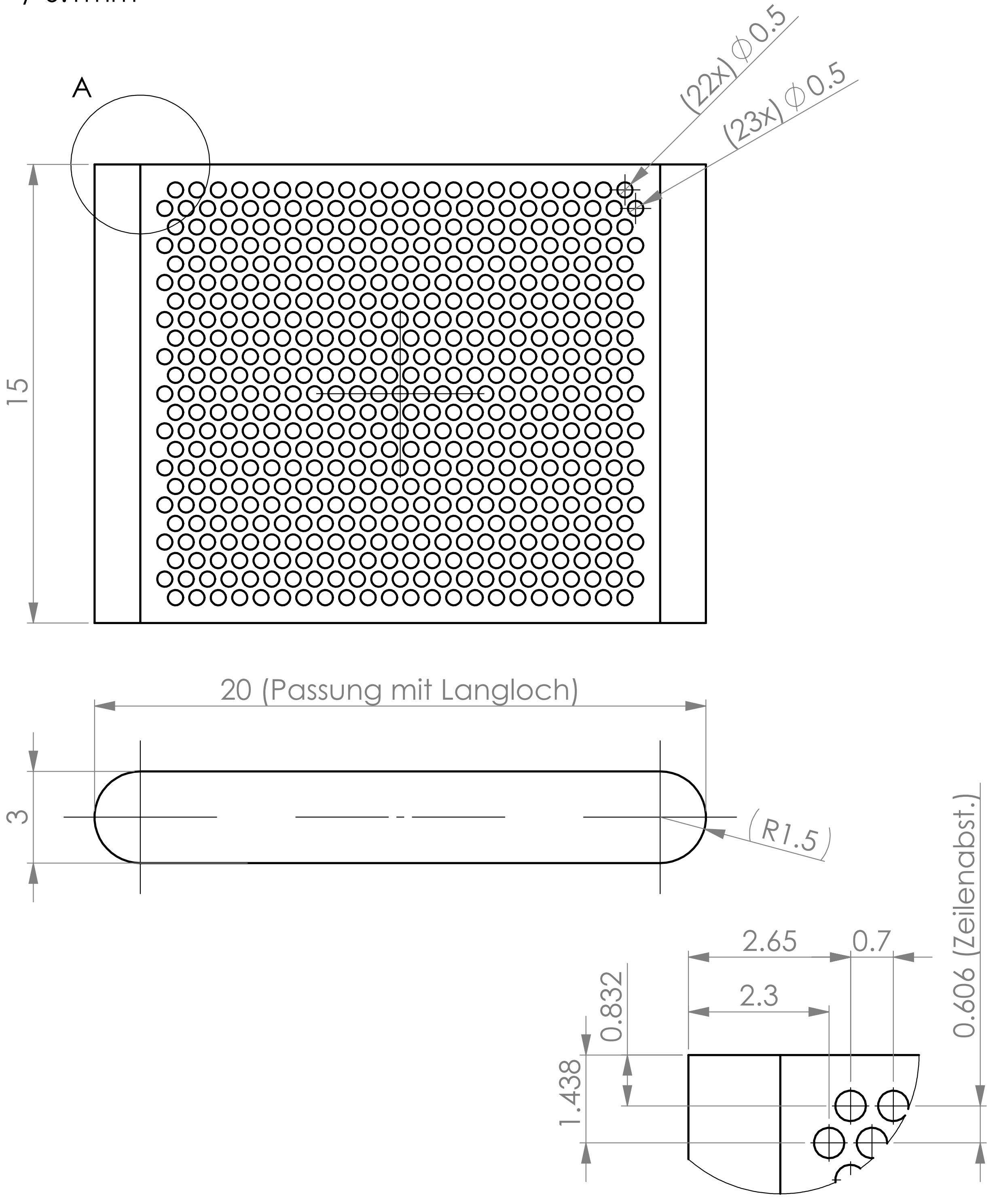


A (10 : 1)

| | | | | |
|---|---|---|---|---|
| 1 | 1 | | POM weiss | |
| Stückzahl | Pos. | Gegenstand | Werkstoff | Bemerkung |

| | | | | |
|---|---|---|---|---|
| UNI BASEL | Schieber für Fish Rig V4 | Massstab 5:1 | Gezeichnet | P. Schlenker |
| | | | Datum | 28.10.2021 |
| | | | Auftraggeber | |
| | | | Blatt Anzahl | 1 von 1 |
| | Biozentrum Zentral Werkstatt | No. | 1003259 | |
