## Supplementary material for "Odor preference maps to cohesive transcriptional domains in the olfactory bulb": 1003266.pdf

Alle Kanten leicht gebrochen

Alle Masse +/-0.1mm

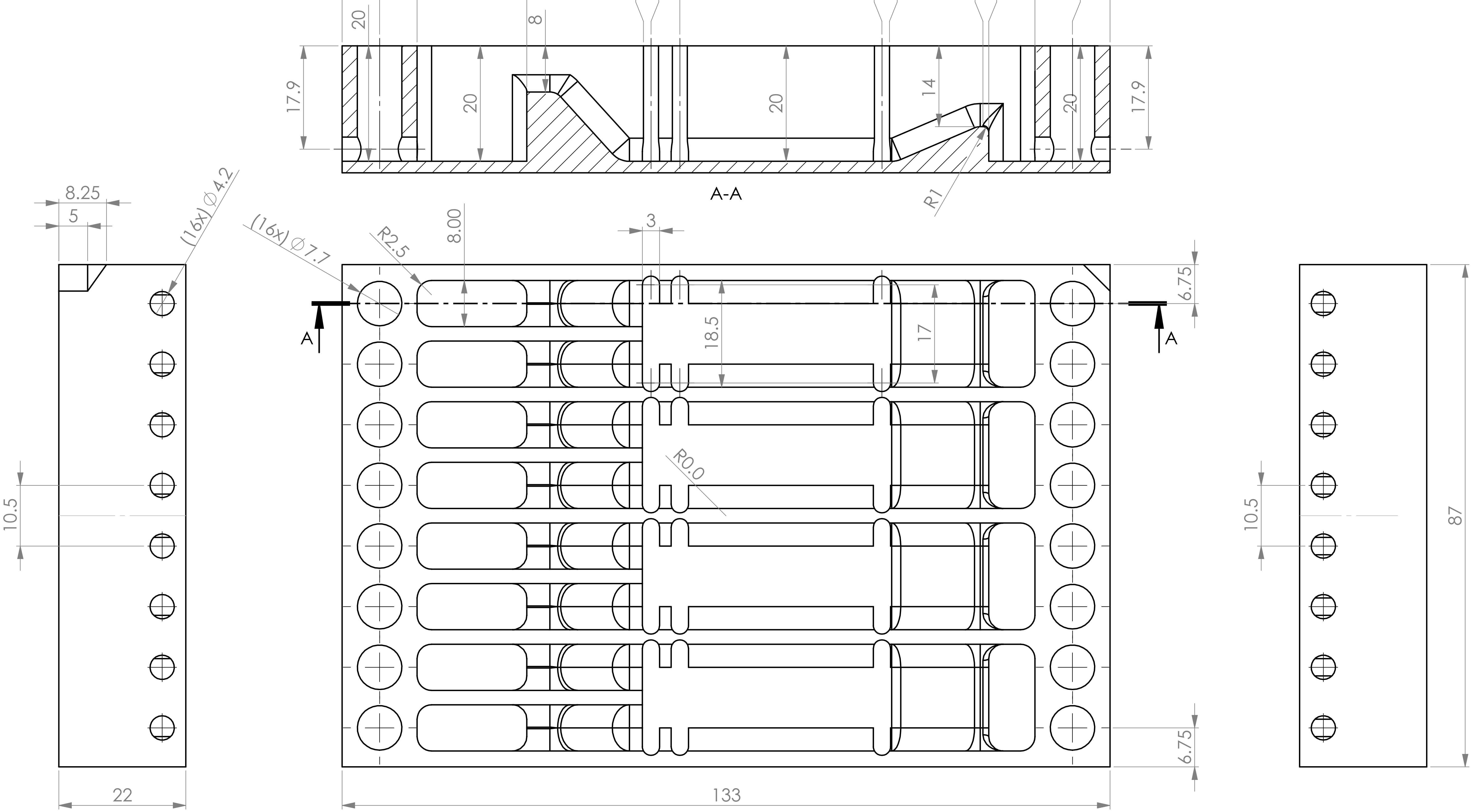


| Stückzahl | Pos. | Gegenstand | Werkstoff | Bemerkung |
|---|---|---|---|---|
| 1 | 1 | | POM Weiss | |

| | | | |
|---|---|---|---|
| UNI BASEL | Fish Rig V4.1 | Massstab 1.5:1 | Gezeichnet: P. Schlenker |
| | | | Datum: 09.11.2021 |
| | | | Auftraggeber: |
| | | | Blatt Anzahl: 1 von 2 |
| | Biozentrum | Zentral Werkstatt | No. 1003266 |

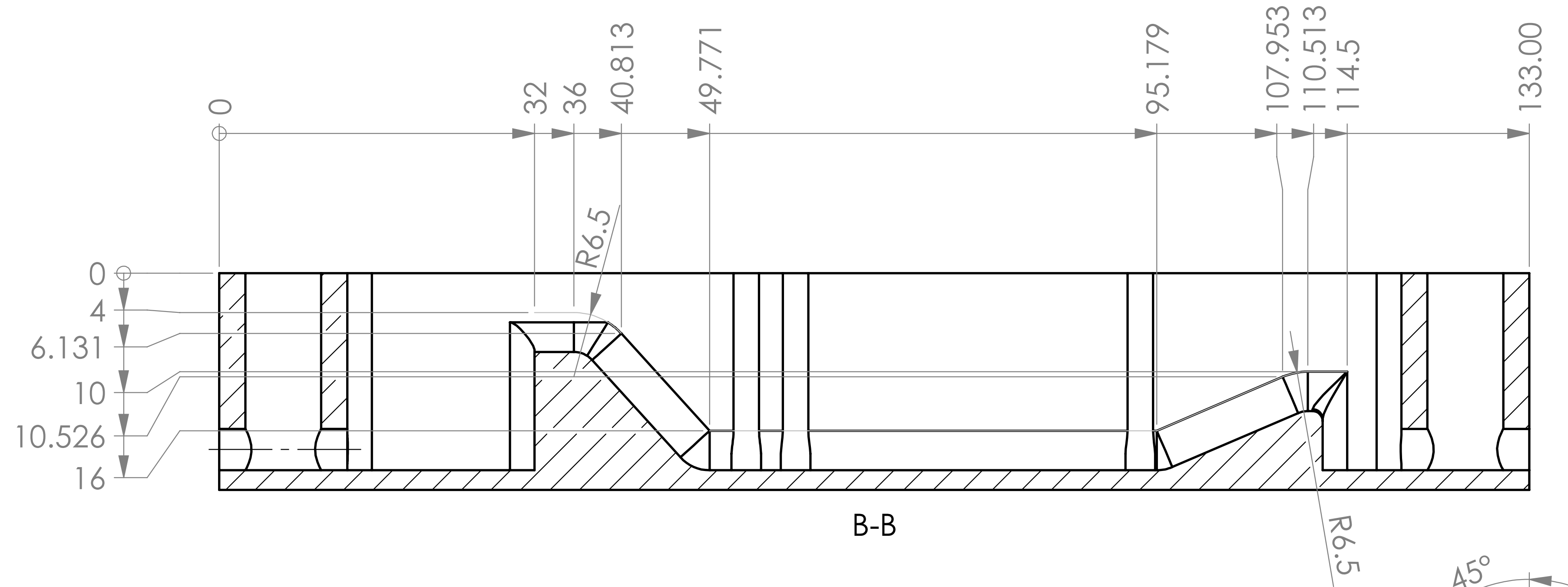


B-B

**ACHTUNG:**
**Vermasste Bahn für Mittelpunkt Kugelkopf KF D8mm**

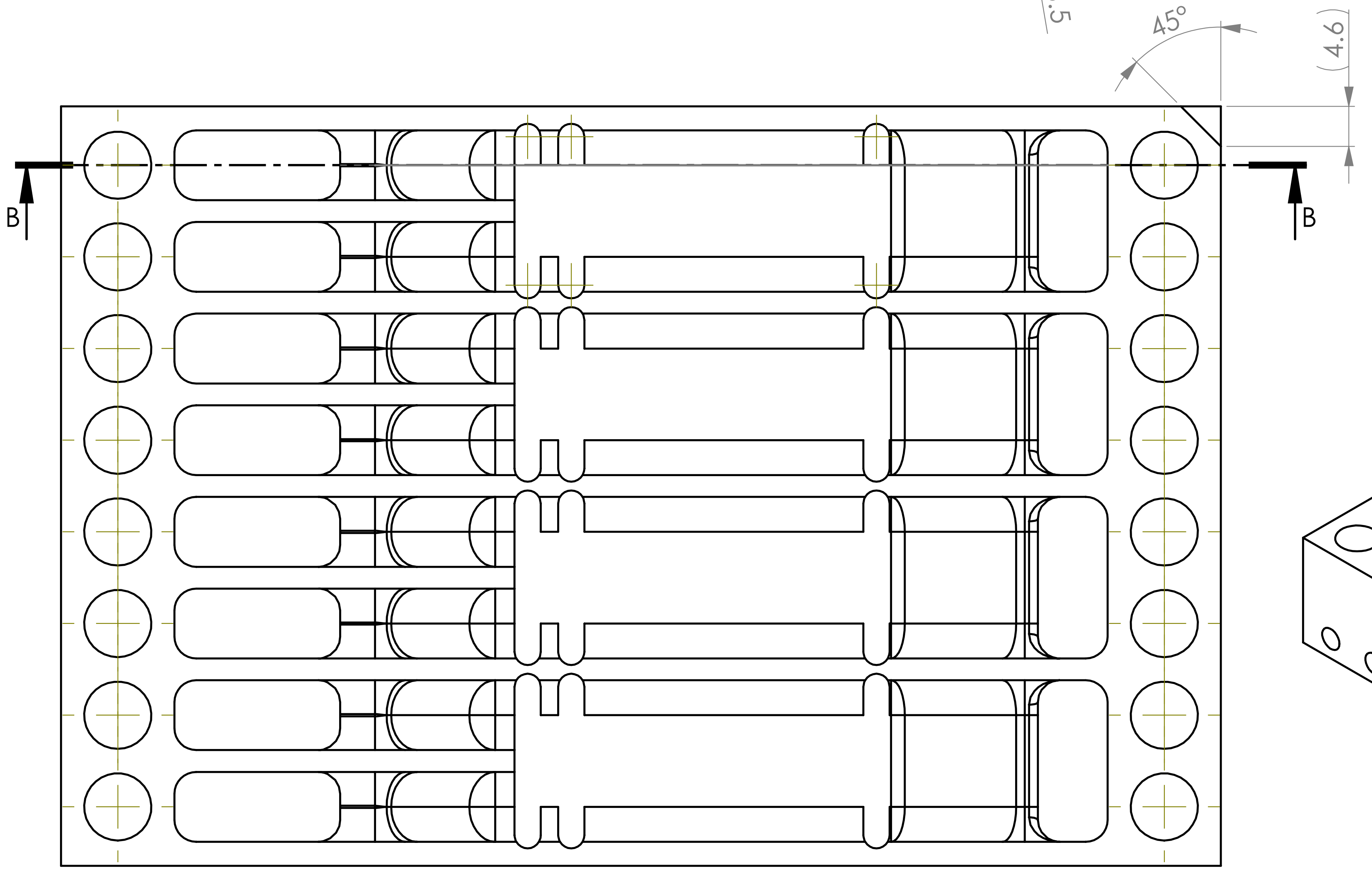


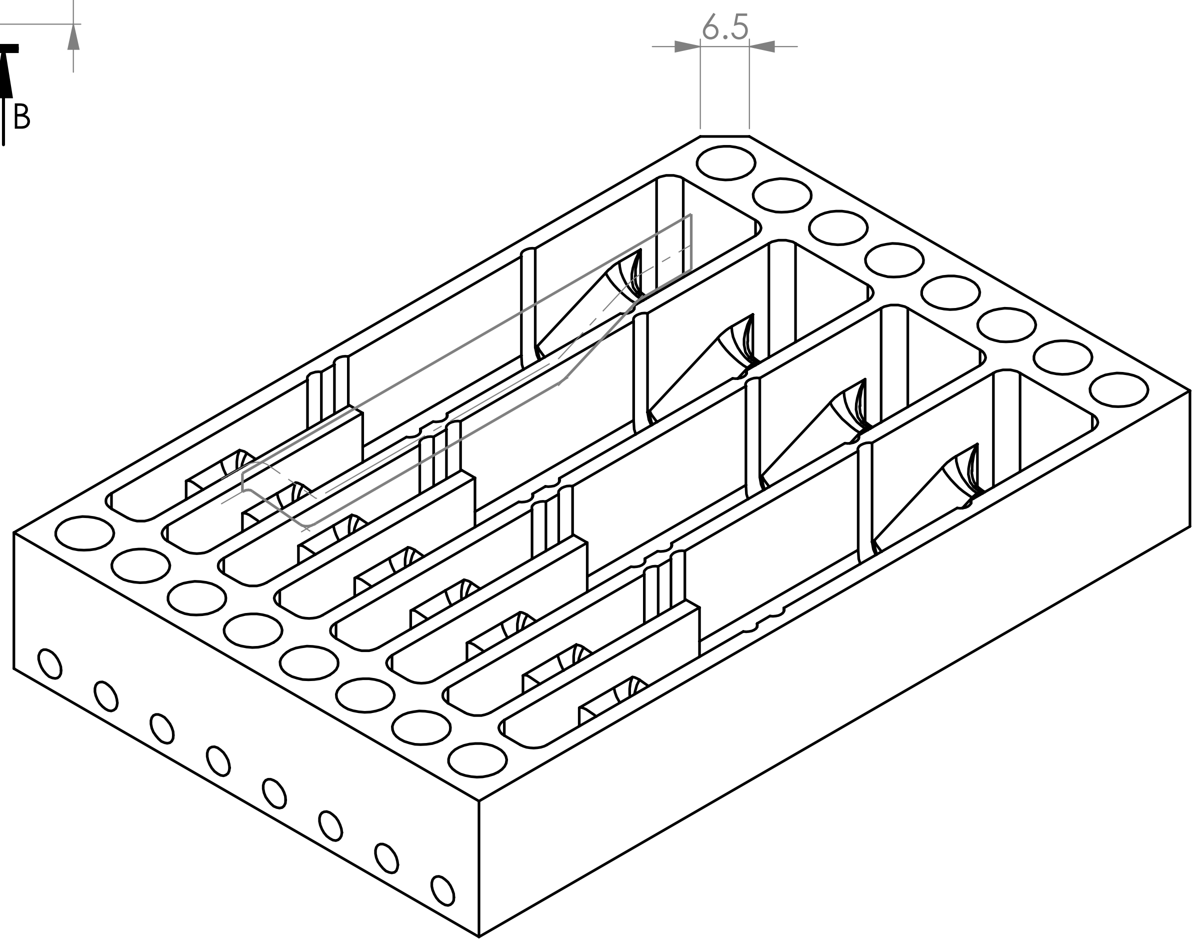


| | | | | |
|---|---|---|---|---|
| 1 | 1 | | POM Weiss | |
| Stückzahl | Pos. | Gegenstand | Werkstoff | Bemerkung |
| UNI BASEL | Fish Rig V4.1 | Massstab 1.5:1 | Gezeichnet | P. Schlenker |
| | | | Datum | 09.11.2021 |
| | | | Auftraggeber | |
| | | | Blatt Anzahl | 2 von 2 |
| | Biozentrum | Zentral Werkstatt | No. | 1003257 |
