## Supplementary material for "Odor preference maps to cohesive transcriptional domains in the olfactory bulb": 1003281.pdf

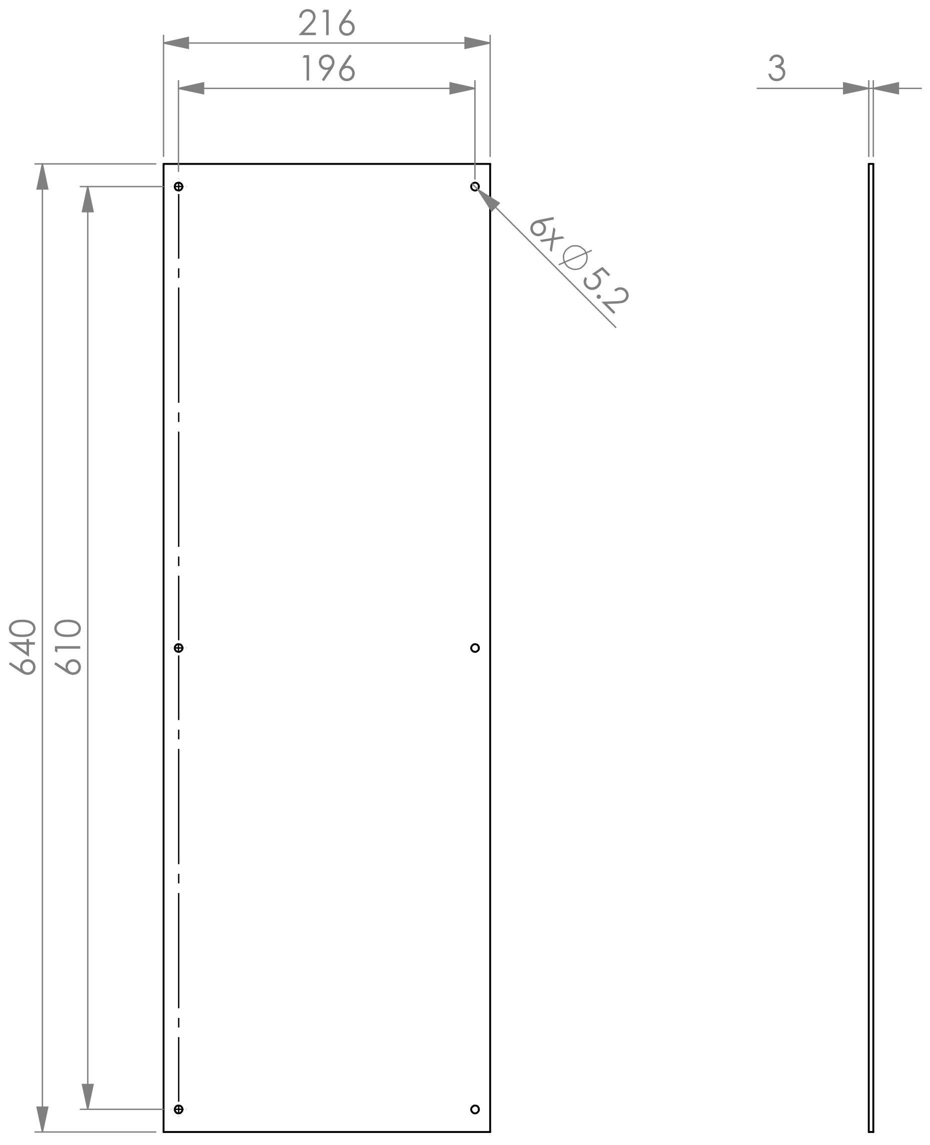


| | |
|---|---|
| WENN NICHT ANDERS DEFINIERT:<br>BEMASSUNGEN SIND IN MILLIMETER<br>OBERFLÄCHENBESCHAFFENHEIT:<br>TOLERANZEN:<br>LINEAR:<br>WINKEL: | OBERFLÄCHENGÜTE: |

ENTGRATEN UND SCHARFE KANTEN BRECHEN

ZEICHNUNG NICHT SKALIEREN

ÄNDERUNG

| | NAME | SIGNATUR | DATUM |
|---|---|---|---|
| GEZEICHNET | | | |
| GEPRÜFT | | | |
| GENEHMIGT | | | |
| PRODUKTION | | | |
| QUALITÄT | | | |

BENENNUNG:

WERKSTOFF: PVC hart

GEWICHT:

ZEICHNUNGSNR. 1003281

A4

MASSSTAB:1:5

BLATT 1 VON 1
