## Supplementary material for "Odor preference maps to cohesive transcriptional domains in the olfactory bulb": 1003282.pdf

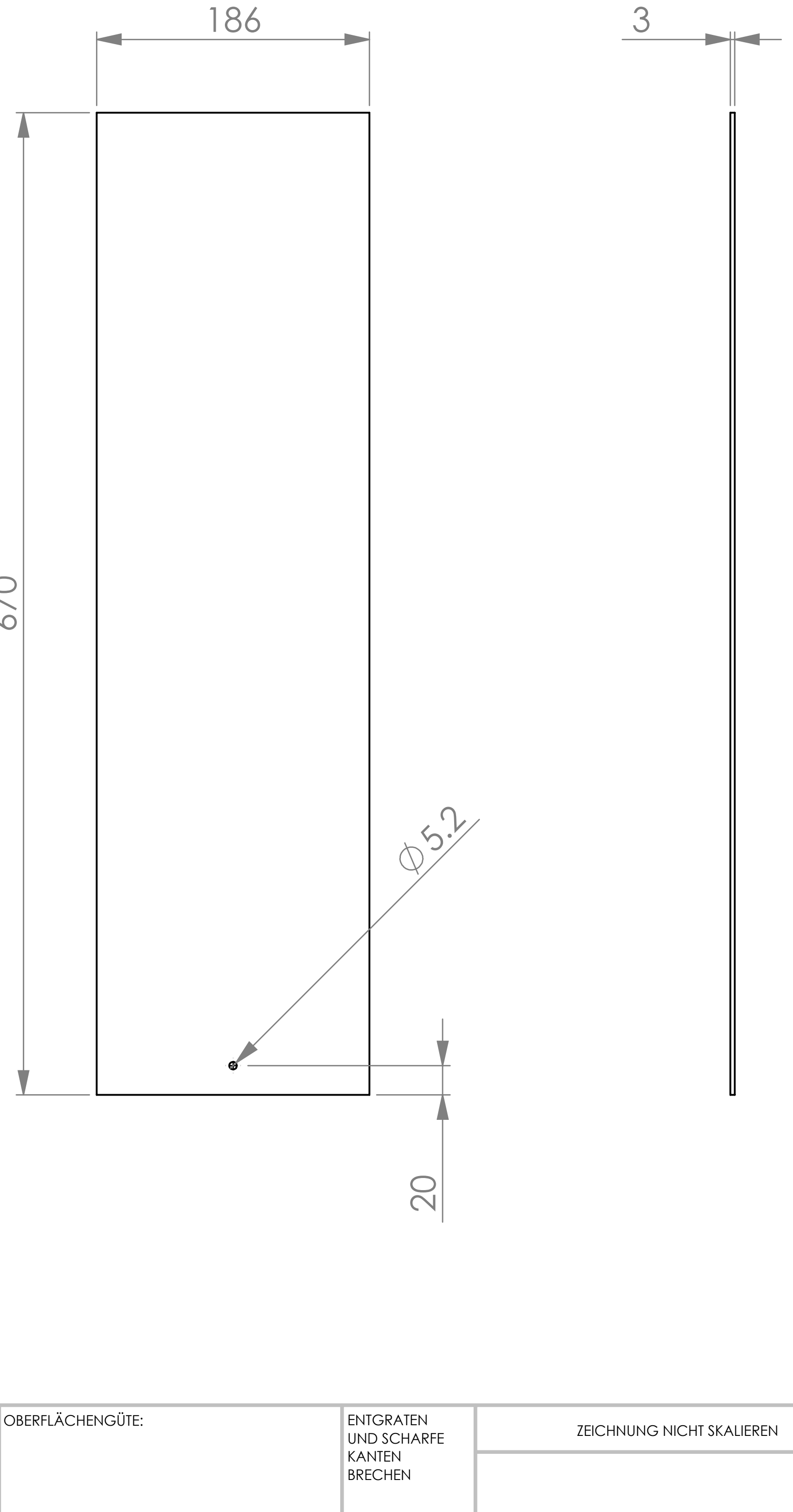


| | | | | | | |
|---|---|---|---|---|---|---|
| WENN NICHT ANDERS DEFINIERT:<br>BEMASSUNGEN SIND IN MILLIMETER<br>OBERFLÄCHENBESCHAFFENHEIT:<br>TOLERANZEN:<br>LINEAR:<br>WINKEL: | OBERFLÄCHENGÜTE: | | | ENTGRATEN UND SCHARFE KANTEN BRECHEN | ZEICHNUNG NICHT SKALIEREN | ÄNDERUNG |
| | NAME | SIGNATUR | DATUM | | BENENNUNG: | |
| GEZEICHNET | | | | | | |
| GEPRÜFT | | | | | | |
| GENEHMIGT | | | | | | |
| PRODUKTION | | | | | | |
| QUALITÄT | | | | WERKSTOFF: PVC hart | ZEICHNUNGSNR. 1003282 | A4 |
| | | | | GEWICHT: | MASSSTAB:1:5 | BLATT 1 VON 1 |
