## Supplementary material for "Odor preference maps to cohesive transcriptional domains in the olfactory bulb": 1003287.pdf

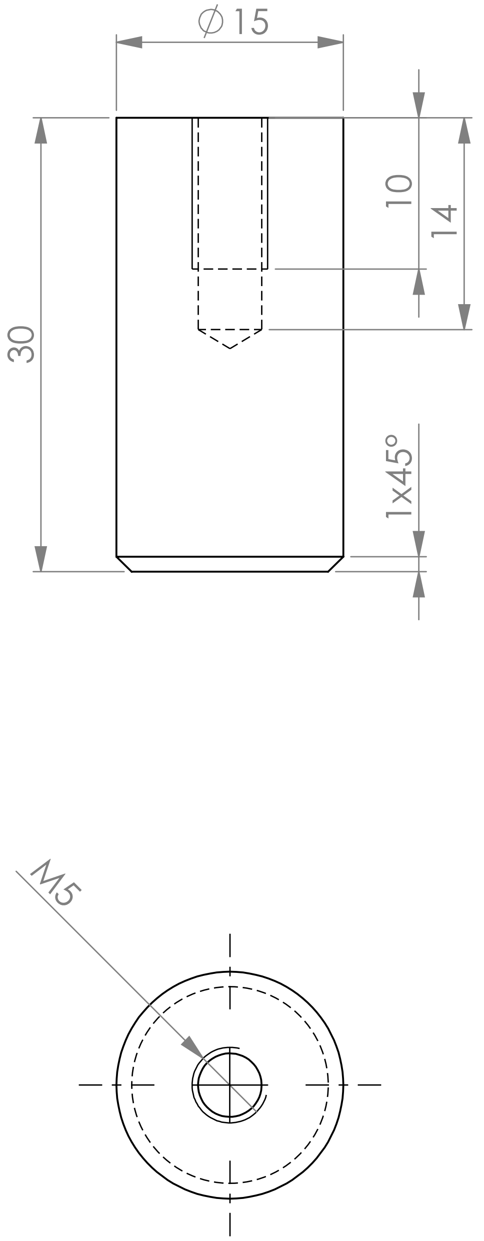

| WENN NICHT ANDERS DEFINIERT:<br>BEMASSUNGEN SIND IN MILLIMETER<br>OBERFLÄCHENBESCHAFFENHEIT:<br>TOLERANZEN:<br>LINEAR:<br>WINKEL: | OBERFLÄCHENGÜTE: | ENTGRATEN UND SCHARFE KANTEN BRECHEN | ZEICHNUNG NICHT SKALIEREN | ÄNDERUNG |
|---|---|---|---|---|

| | NAME | SIGNATUR | DATUM |
|---|---|---|---|
| GEZEICHNET | | | |
| GEPRÜFT | | | |
| GENEHMIGT | | | |
| PRODUKTION | | | |
| QUALITÄT | | | |

BENENNUNG:

WERKSTOFF: 1060 Legierung

ZEICHNUNGSNR. 1003287

A4

GEWICHT:

MASSSTAB:2:1

BLATT 1 VON 1
