## Supplementary material for "Odor preference maps to cohesive transcriptional domains in the olfactory bulb": 1003288.pdf

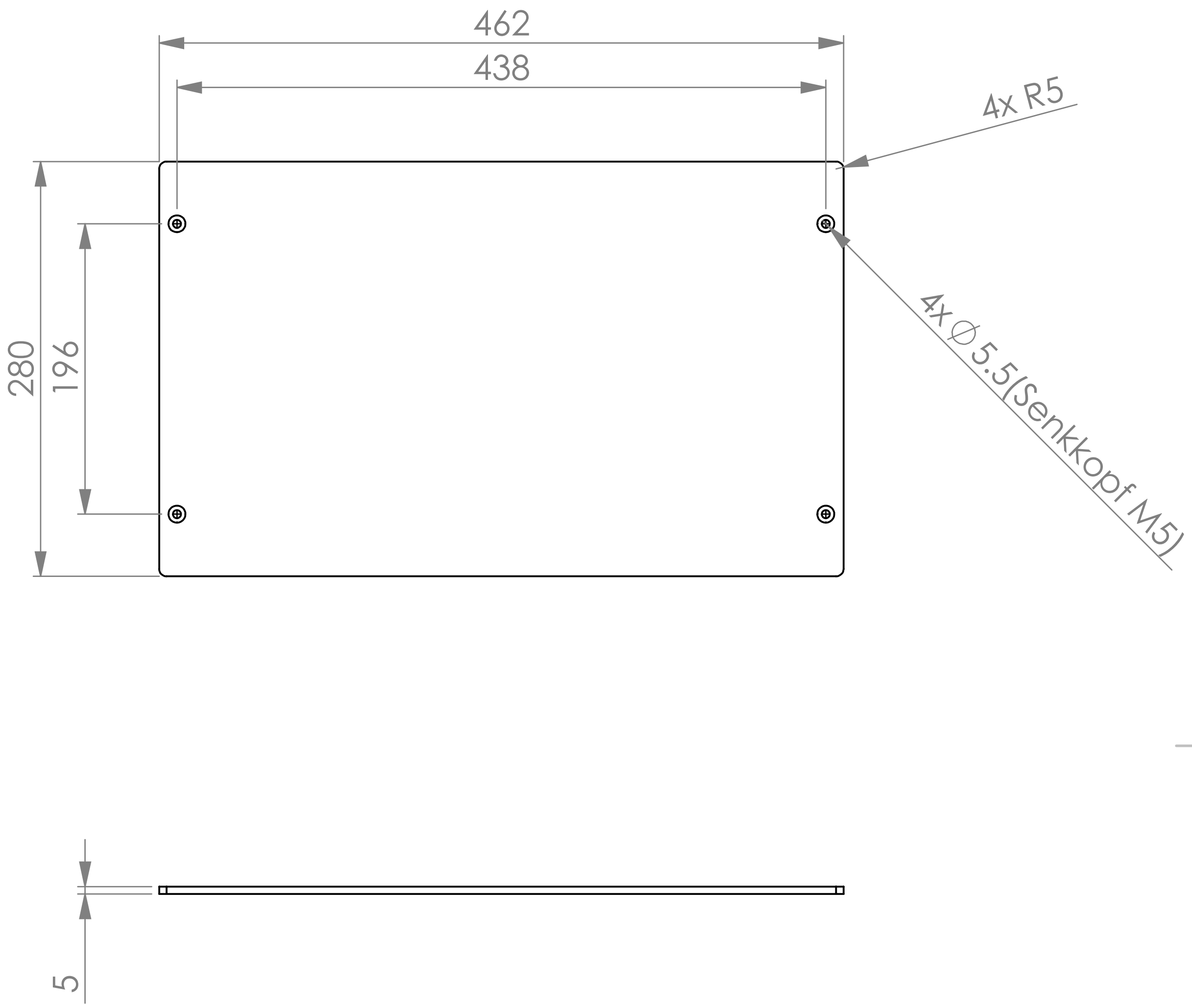

| | | | | | |
|---|---|---|---|---|---|
| WENN NICHT ANDERS DEFINIERT: BEMASSUNGEN SIND IN MILLIMETER OBERFLÄCHENBESCHAFFENHEIT: TOLERANZEN: LINEAR: WINKEL: | OBERFLÄCHENGÜTE: | | ENTGRATEN UND SCHARFE KANTEN BRECHEN | ZEICHNUNG NICHT SKALIEREN | ÄNDERUNG |
| | NAME | SIGNATUR | DATUM | BENENNUNG: | |
| GEZEICHNET | | | | | |
| GEPRÜFT | | | | | |
| GENEHMIGT | | | | | |
| PRODUKTION | | | | | |
| QUALITÄT | | | WERKSTOFF: 1060 Legierung | ZEICHNUNGSNR. 1003288 | A4 |
| | | | GEWICHT: | MASSSTAB:1:10 | BLATT 1 VON 1 |
