## Supplementary material for "Odor preference maps to cohesive transcriptional domains in the olfactory bulb": 1003289.pdf

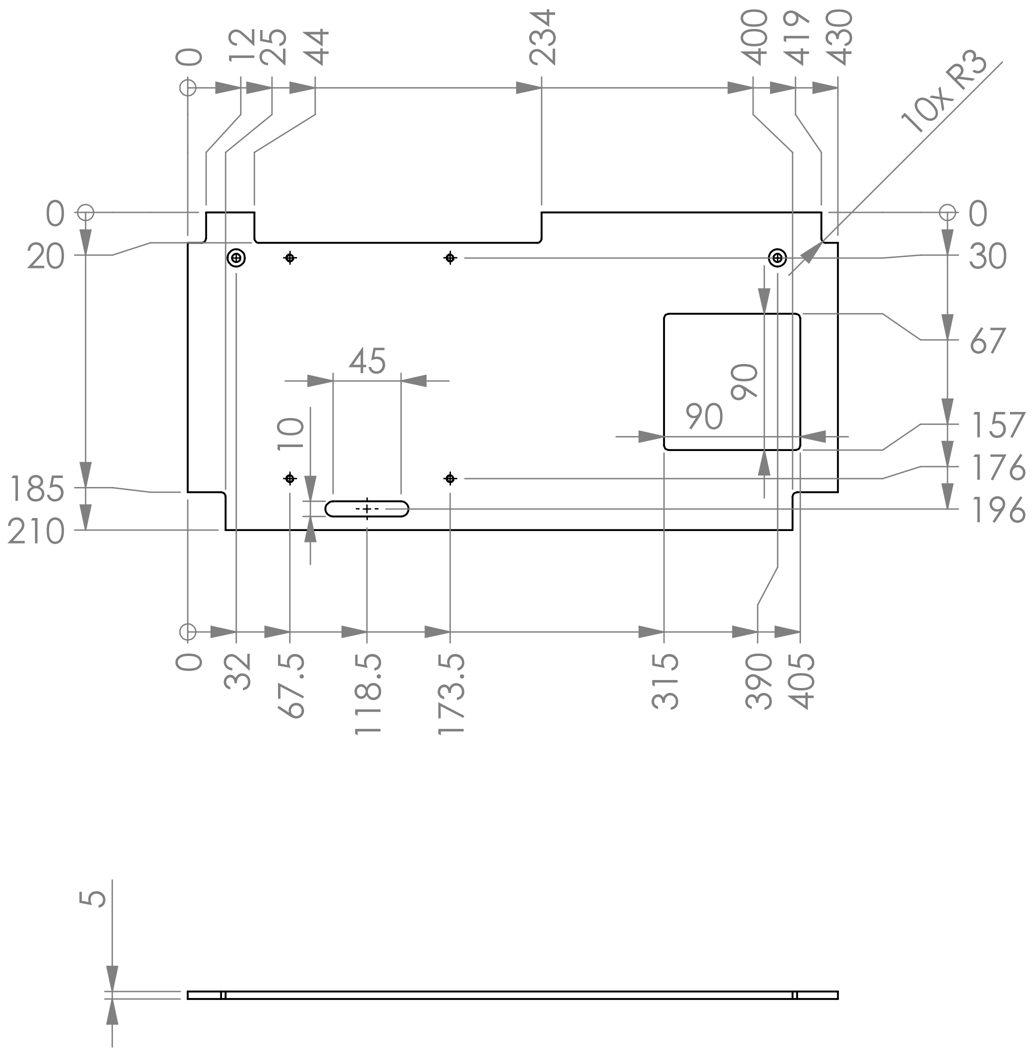


| | | | |
|---|---|---|---|
| WENN NICHT ANDERS DEFINIERT:<br>BEMASSUNGEN SIND IN MILLIMETER<br>OBERFLÄCHENBESCHAFFENHEIT:<br>TOLERANZEN:<br>LINEAR:<br>WINKEL: | OBERFLÄCHENGÜTE: | ENTGRATEN<br>UND SCHARFE<br>KANTEN<br>BRECHEN | ZEICHNUNG NICHT SKALIEREN / ÄNDERUNG |

| | NAME | SIGNATUR | DATUM |
|---|---|---|---|
| GEZEICHNET | | | |
| GEPRÜFT | | | |
| GENEHMIGT | | | |
| PRODUKTION | | | |
| QUALITÄT | | | |

BENENNUNG:

WERKSTOFF:

ZEICHNUNGSNR.

1003289

A4

GEWICHT:

MASSSTAB:1:5

BLATT 1 VON 1
