## Supplementary material for "Odor preference maps to cohesive transcriptional domains in the olfactory bulb": 1003313.pdf

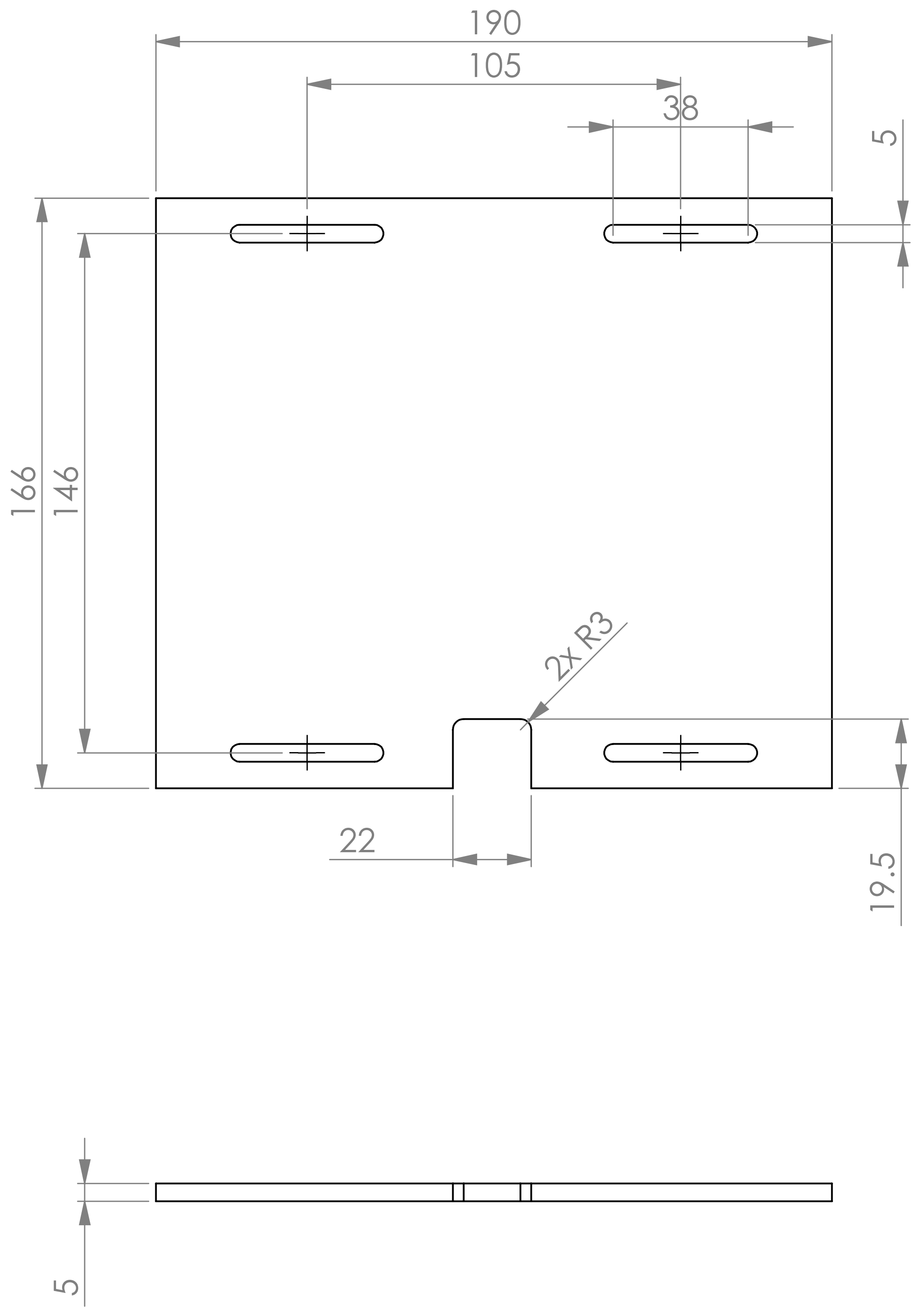


WENN NICHT ANDERS DEFINIERT:
BEMASSUNGEN SIND IN MILLIMETER
OBERFLÄCHENBESCHAFFENHEIT:
TOLERANZEN:
LINEAR:
WINKEL:

OBERFLÄCHENGÜTE:

ENTGRATEN UND SCHARFE KANTEN BRECHEN

ZEICHNUNG NICHT SKALIEREN

ÄNDERUNG

| | NAME | SIGNATUR | DATUM |
|---|---|---|---|
| GEZEICHNET | | | |
| GEPRÜFT | | | |
| GENEHMIGT | | | |
| PRODUKTION | | | |
| QUALITÄT | | | |

BENENNUNG:

WERKSTOFF:

ZEICHNUNGSNR.

1003313

A4

GEWICHT:

MASSSTAB:1:5

BLATT 1 VON 1
