## Supplementary figures and images for "Odor preference maps to cohesive transcriptional domains in the olfactory bulb"

### 265 Valve Led Driver_PCB.pdf

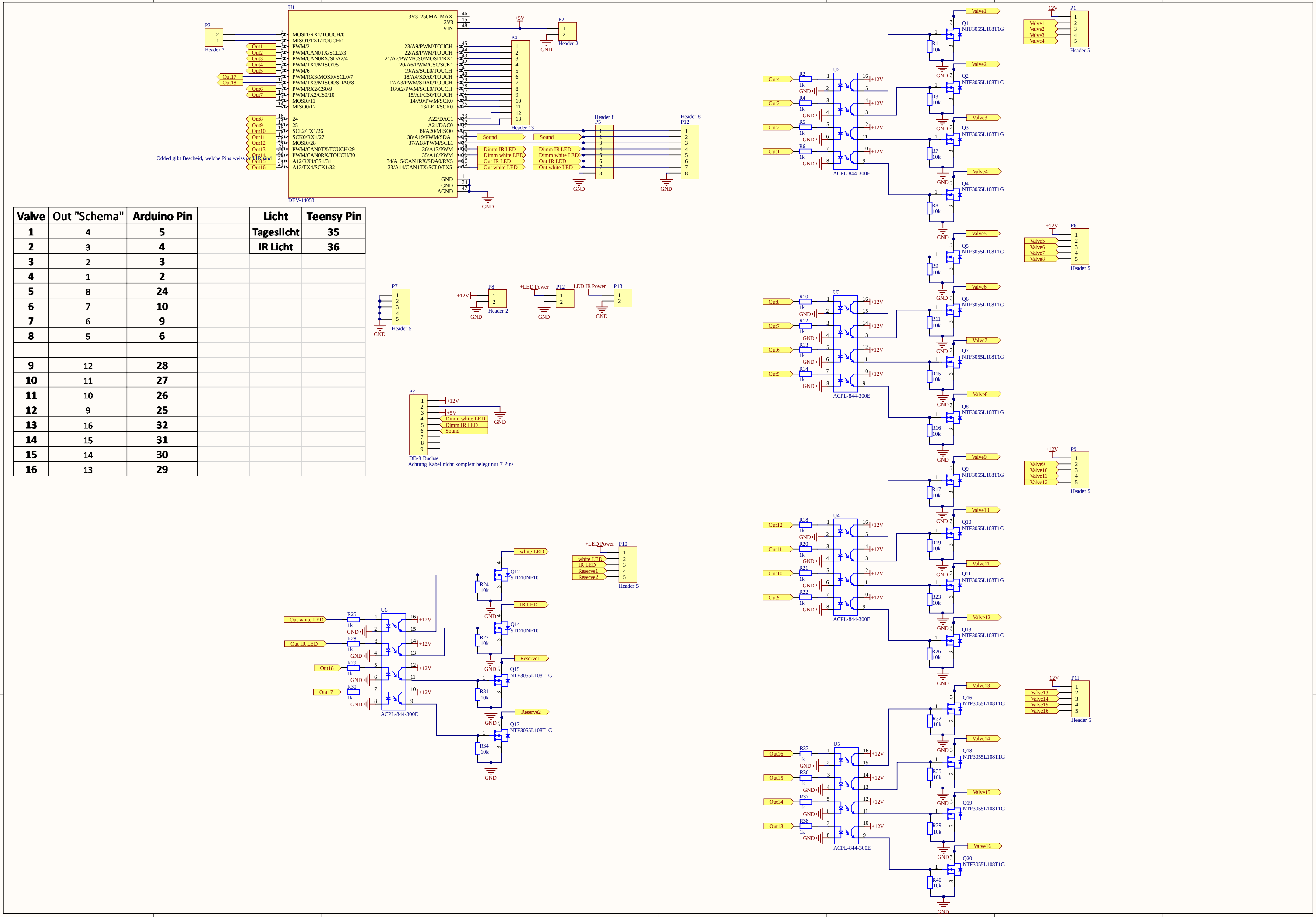

| Valve | Out "Schema" | Arduino Pin |
|---|---|---|
| 1 | 4 | 5 |
| 2 | 3 | 4 |
| 3 | 2 | 3 |
| 4 | 1 | 2 |
| 5 | 8 | 24 |
| 6 | 7 | 10 |
| 7 | 6 | 9 |
| 8 | 5 | 6 |
| 9 | 12 | 28 |
| 10 | 11 | 27 |
| 11 | 10 | 26 |
| 12 | 9 | 25 |
| 13 | 16 | 32 |
| 14 | 15 | 31 |
| 15 | 14 | 30 |
| 16 | 13 | 29 |

| Licht | Teensy Pin |
|---|---|
| Tageslicht | 35 |
| IR Licht | 36 |

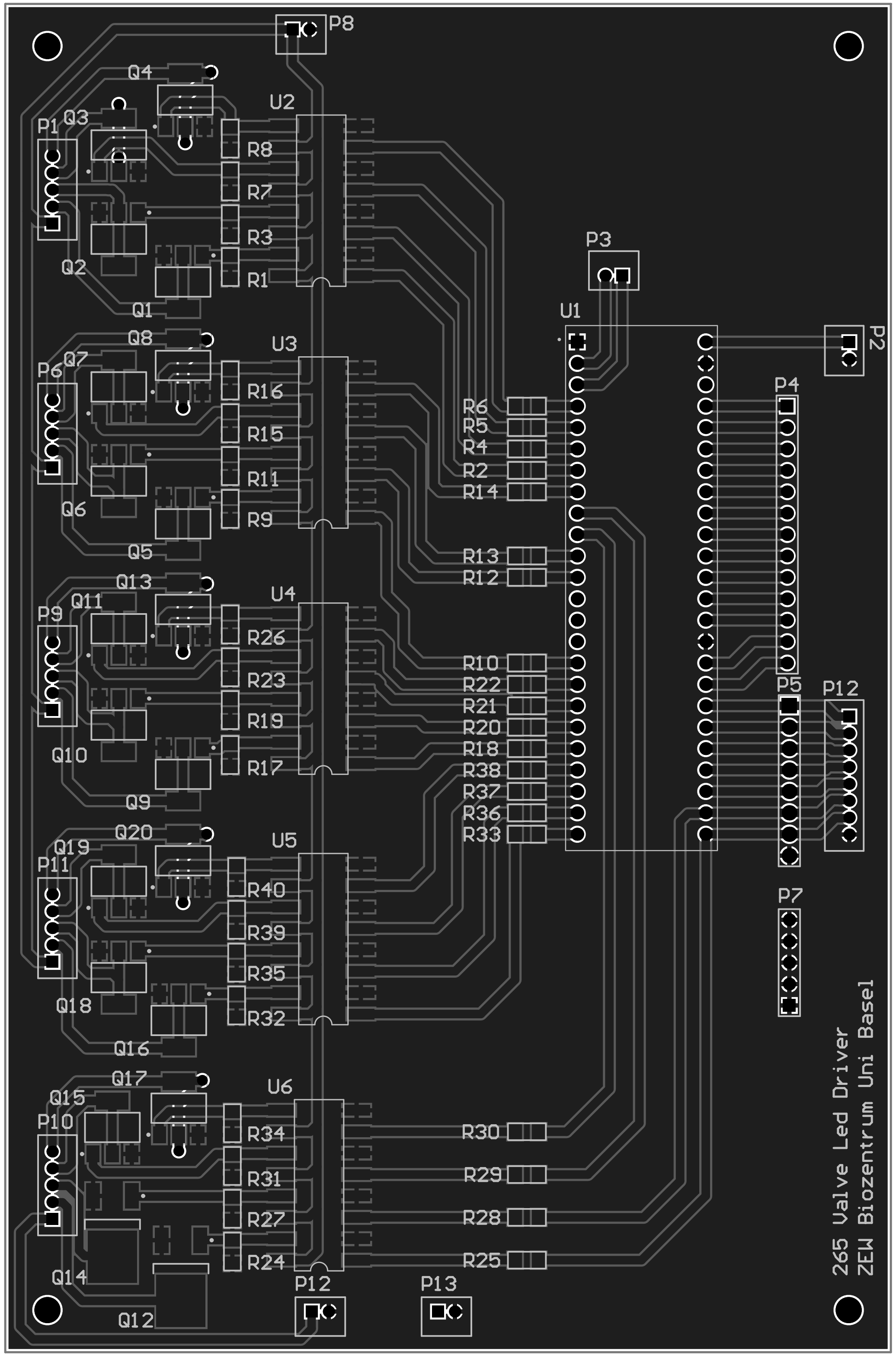

### PCB_NEW_PCB_2026-04-20.pdf

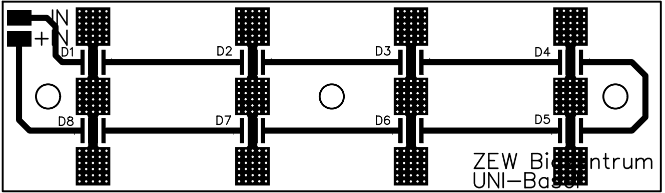
