## supplemental figures for "Odor preference maps to cohesive transcriptional domains in the olfactory bulb"

### Figure S1 I Validation and parameterization of the two-choice olfactory behavior assay

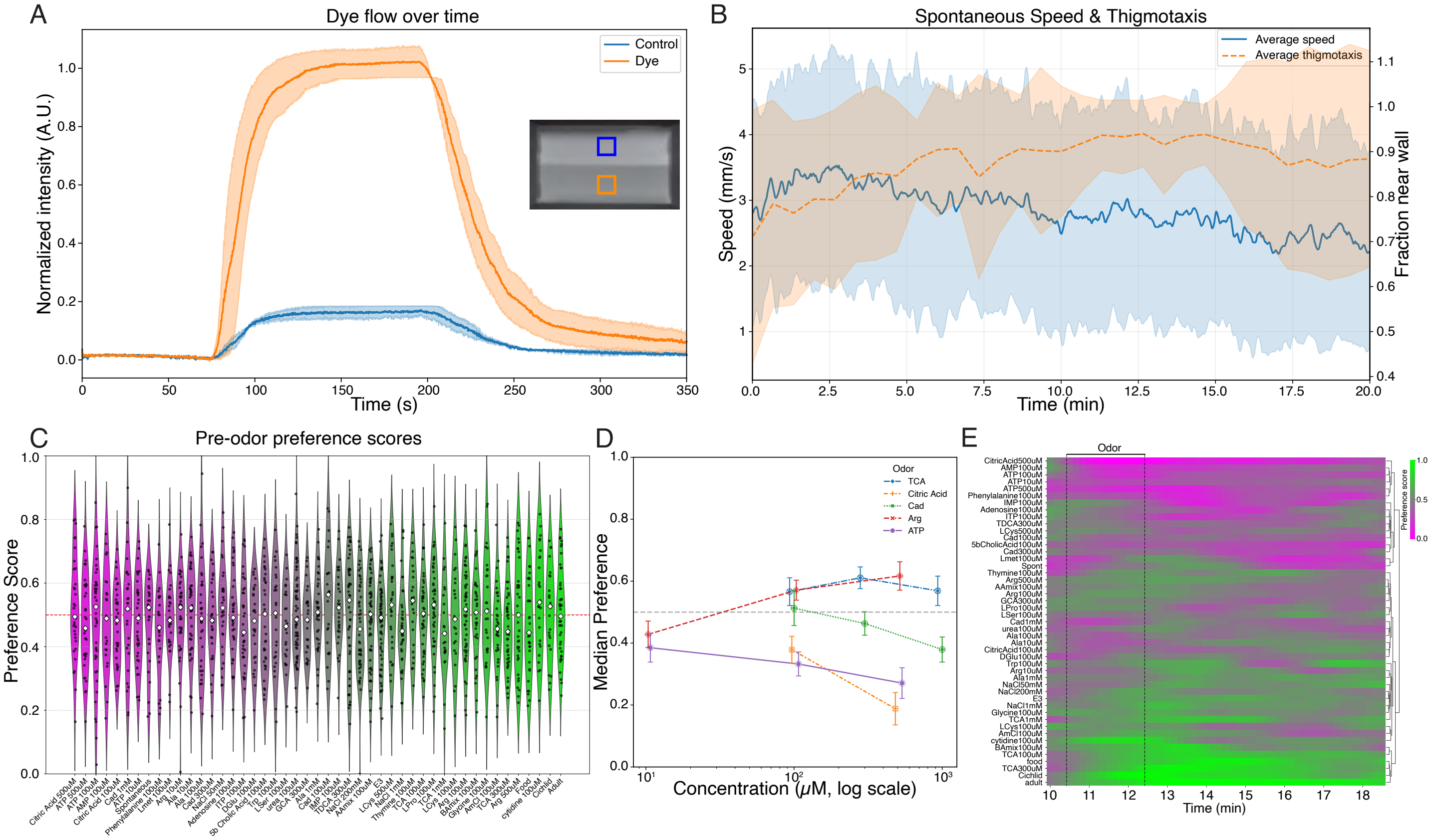

Figure S2 I Time-resolved larval position relative to the odor side

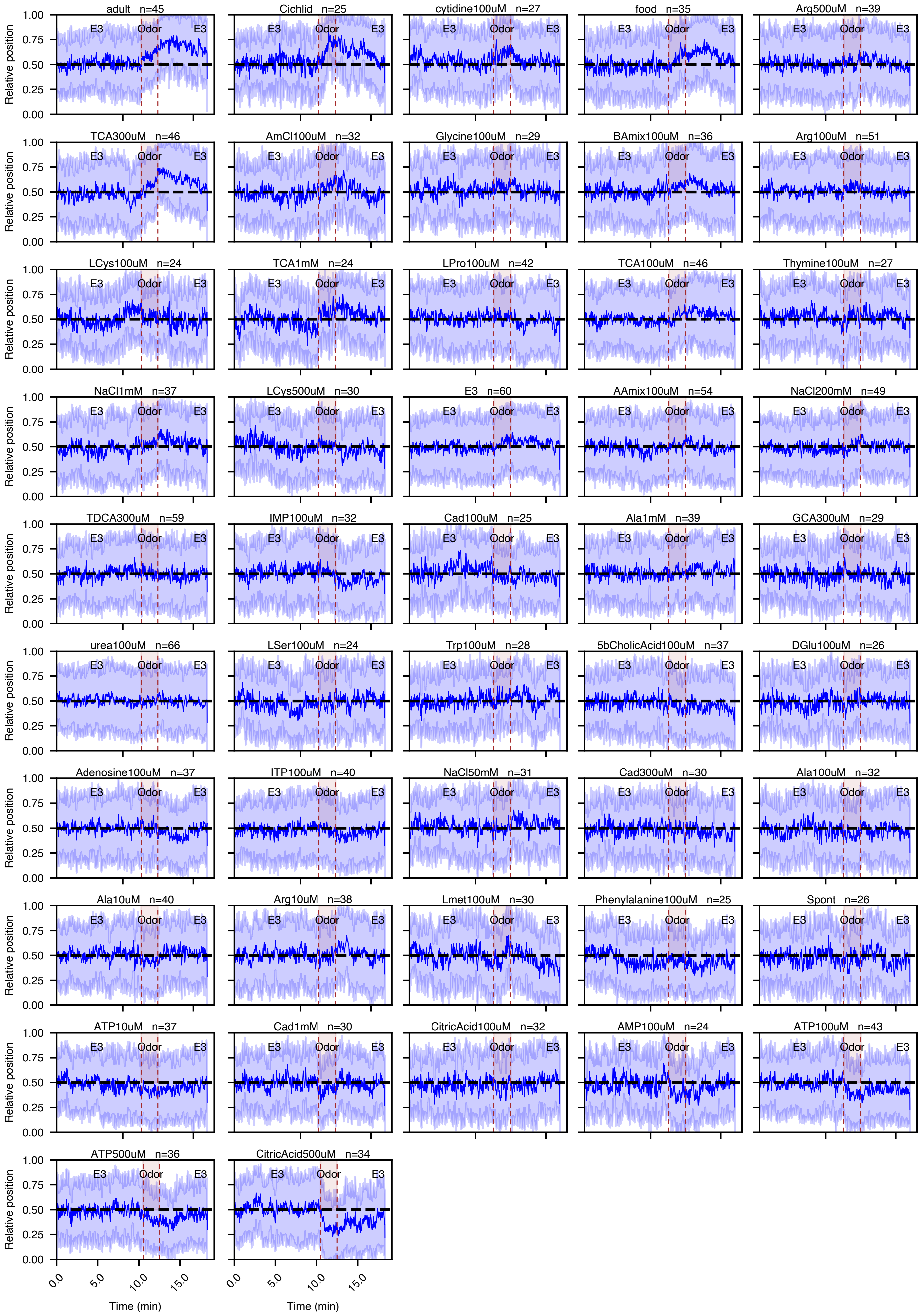

Figure S3 I Cross-modal correspondence of CaMPARI2-derived odor activity maps and activity pattern distribution

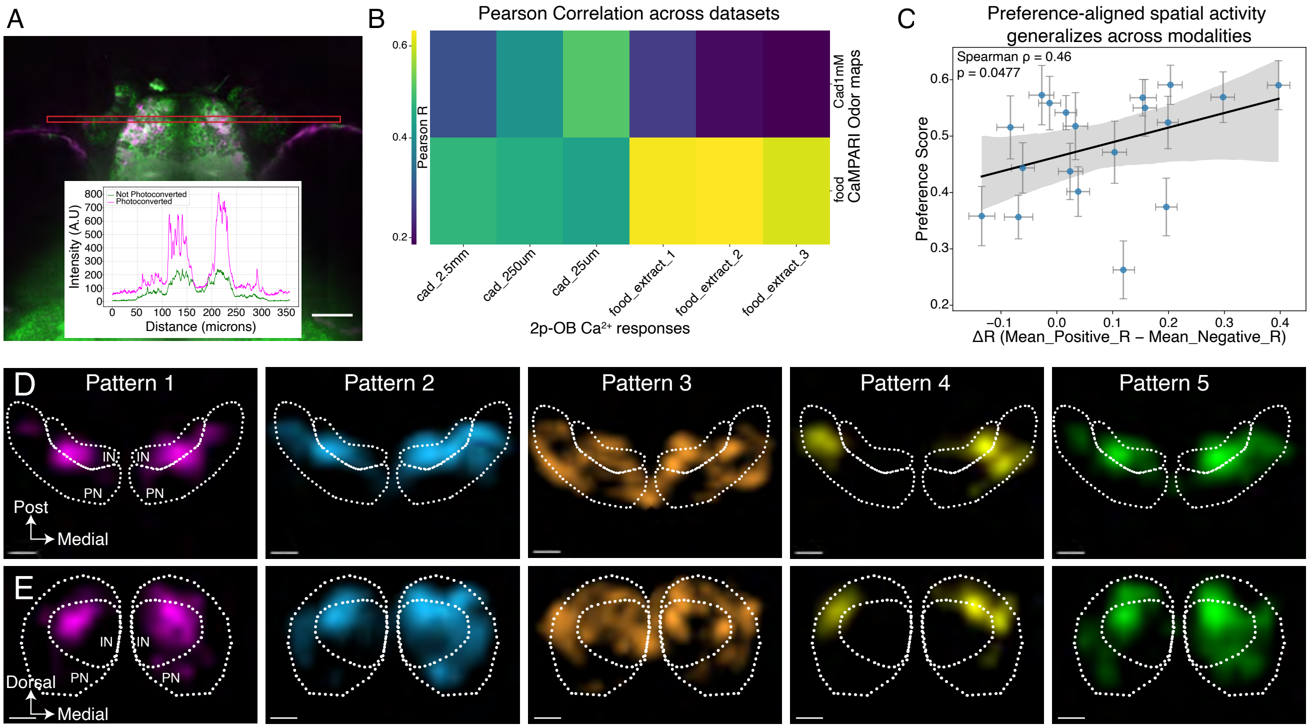

Figure S4 I Odorant-evoked spatial activity maps

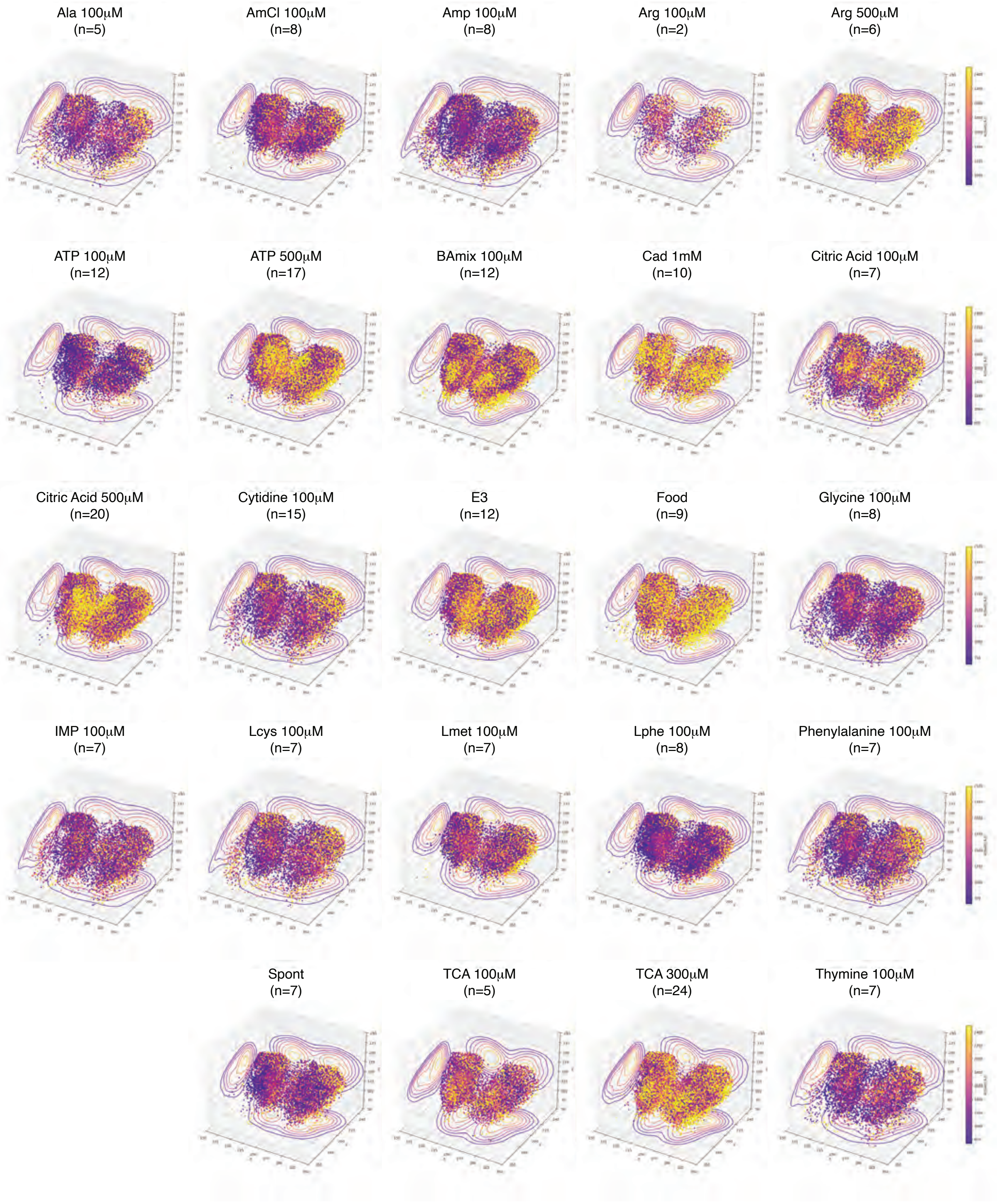

Figure S5 I Combinatorial transcriptional programs define neuronal subtypes and motivate spatial assignment to the olfactory bulb

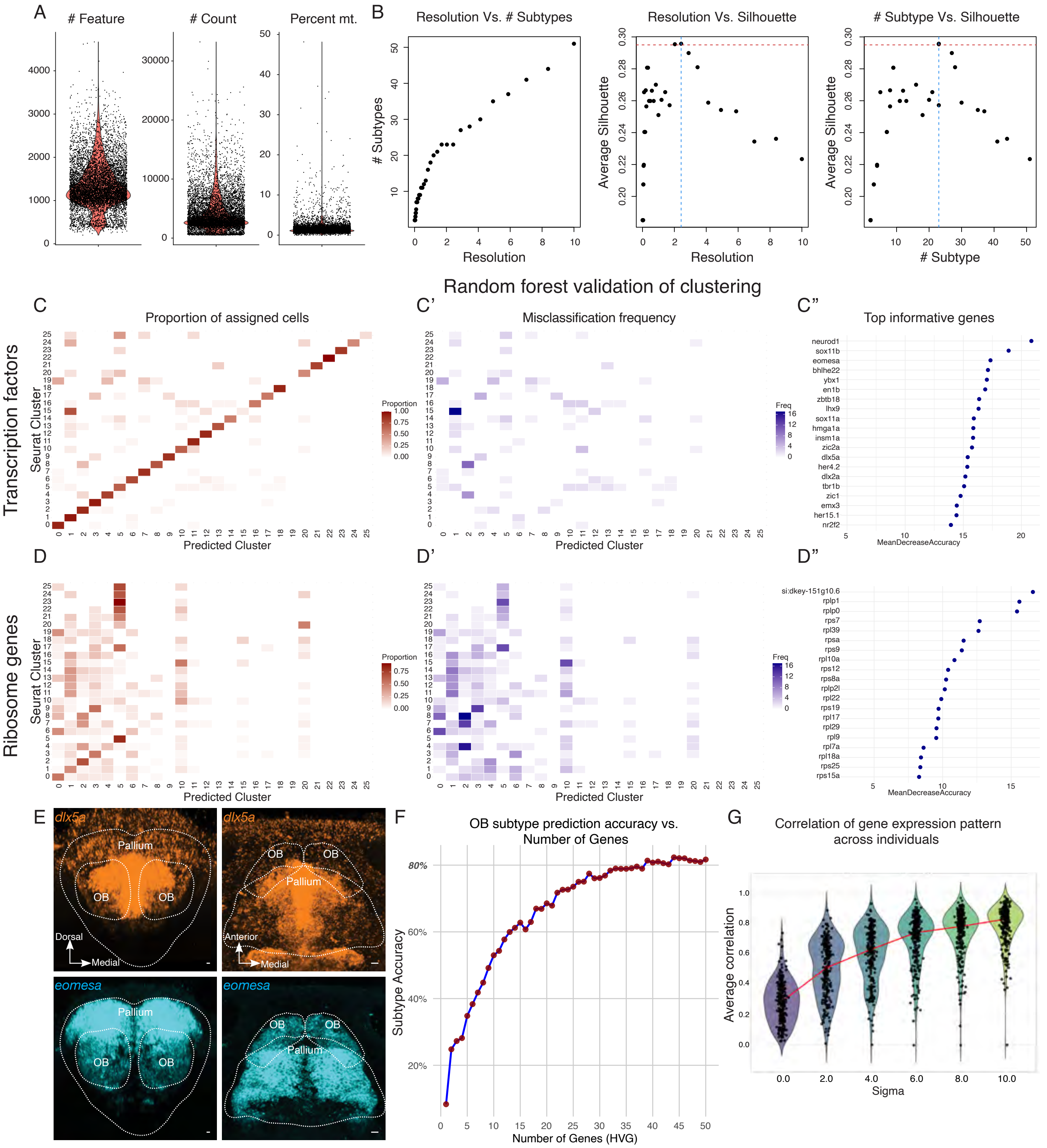

Figure S6 l Spatial transcriptomic maps of marker gene expression in the zebrafish forebrain (Brain1)

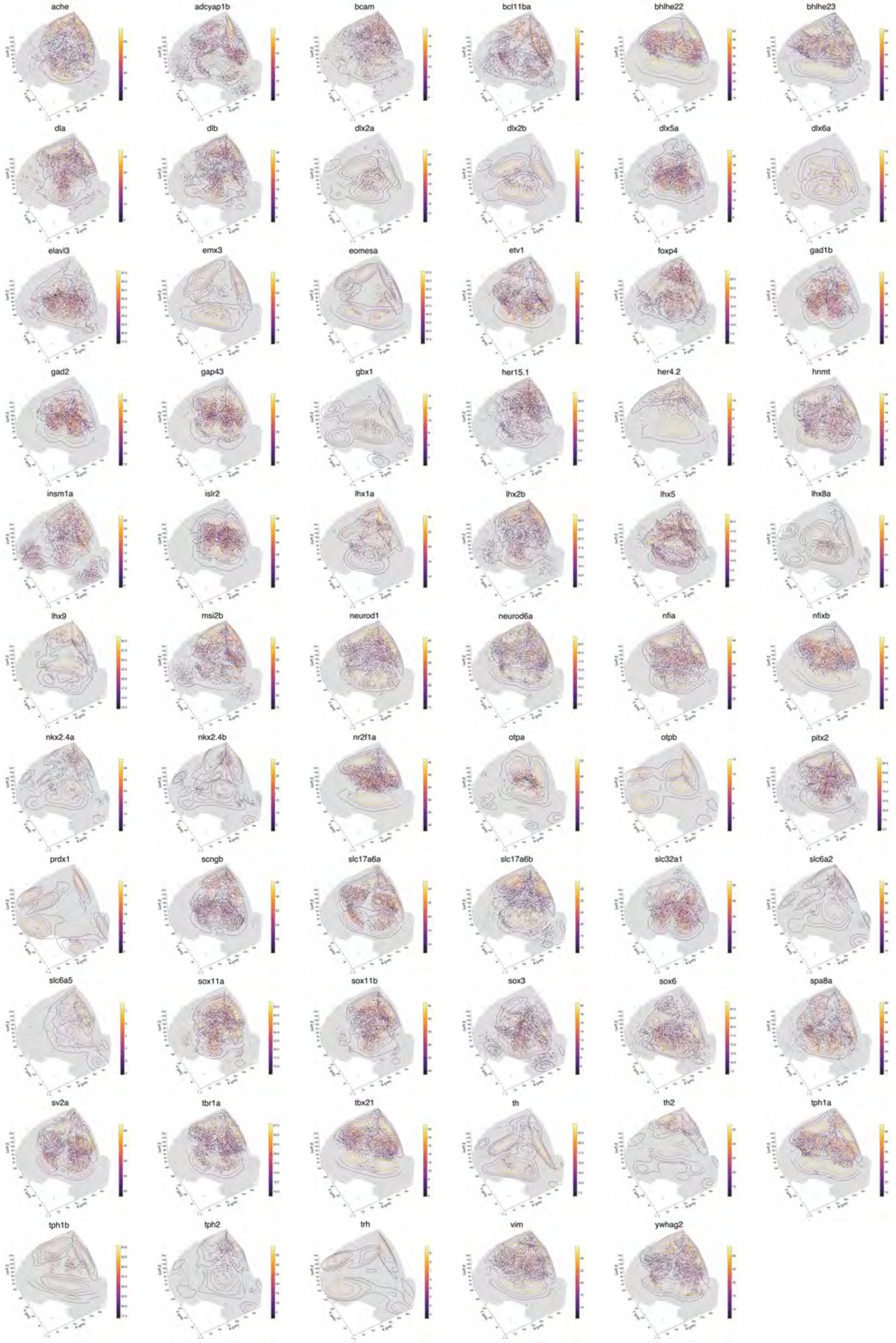

Figure S7 I Spatial transcriptomic maps of marker gene expression in the zebrafish forebrain (Brain2)

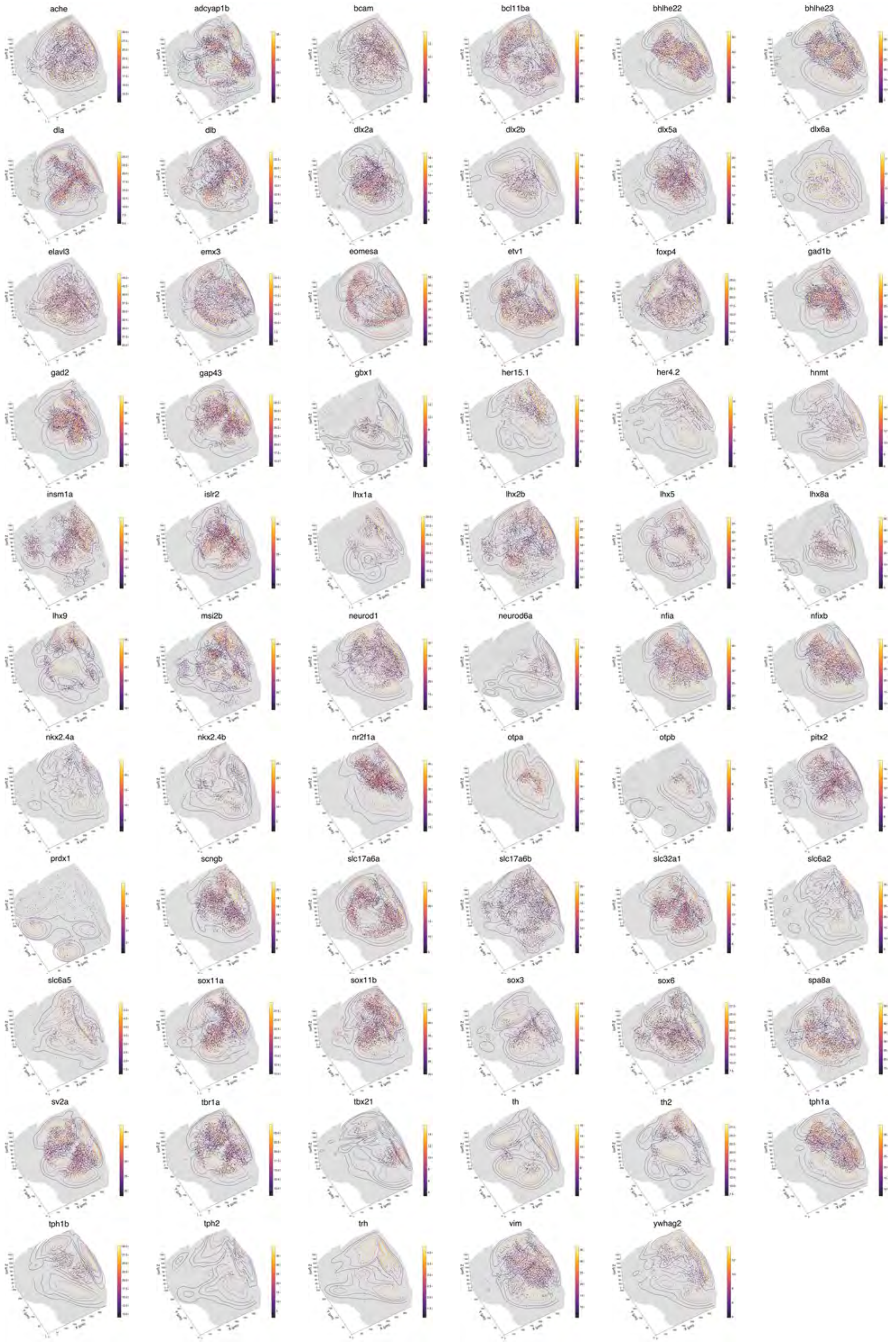

Figure S8 I Tangram imputation accuracy and spatial organization of transcriptionally defined OB subtypes

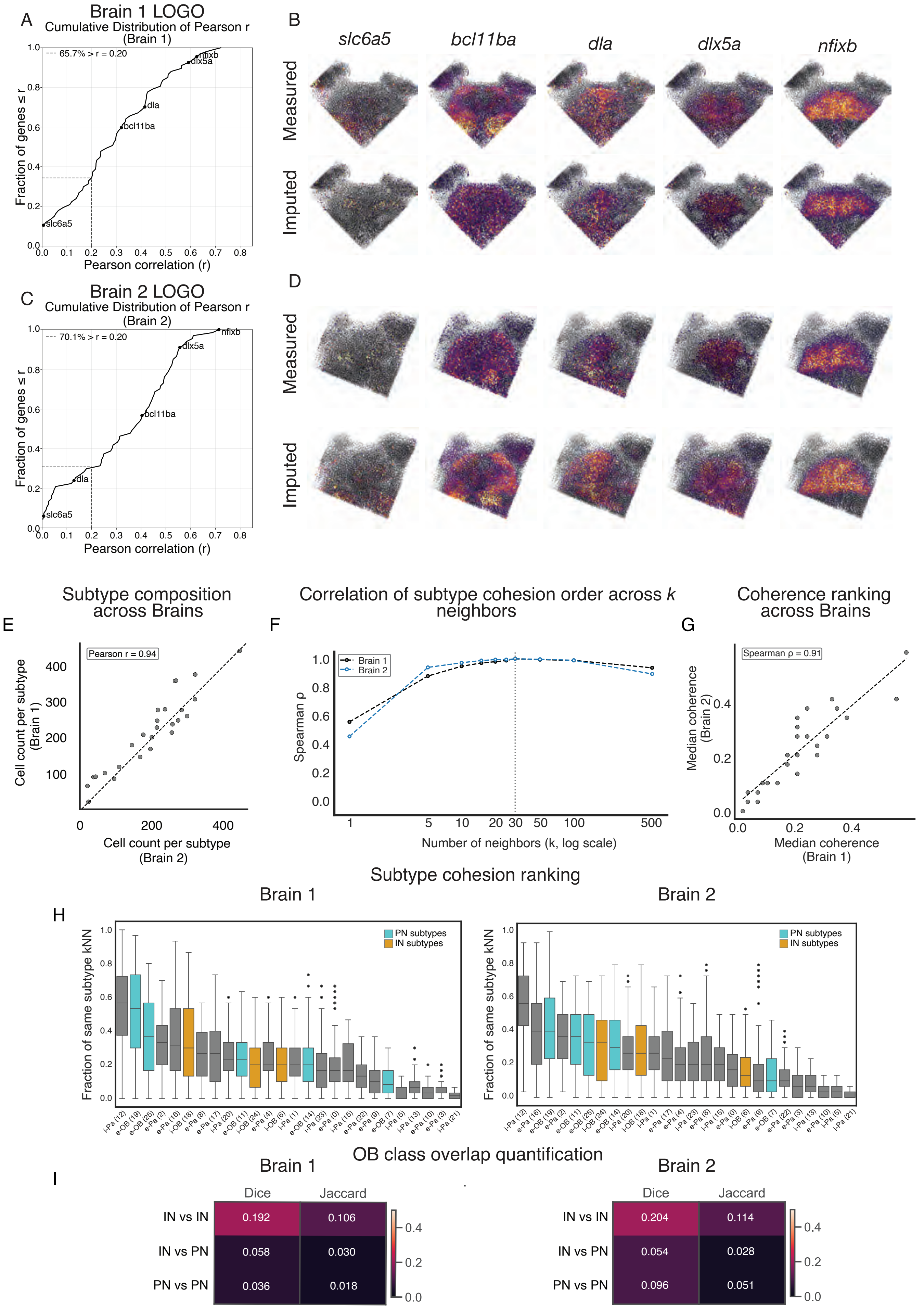


| Brain 1 | Dice | Jaccard |
|---|---|---|
| IN vs IN | 0.192 | 0.106 |
| IN vs PN | 0.058 | 0.030 |
| PN vs PN | 0.036 | 0.018 |

| Brain 2 | Dice | Jaccard |
|---|---|---|
| IN vs IN | 0.204 | 0.114 |
| IN vs PN | 0.054 | 0.028 |
| PN vs PN | 0.096 | 0.051 |

Figure S9 l Quality control and experimental controls for CaMPARI2-seq and two-photon ablation

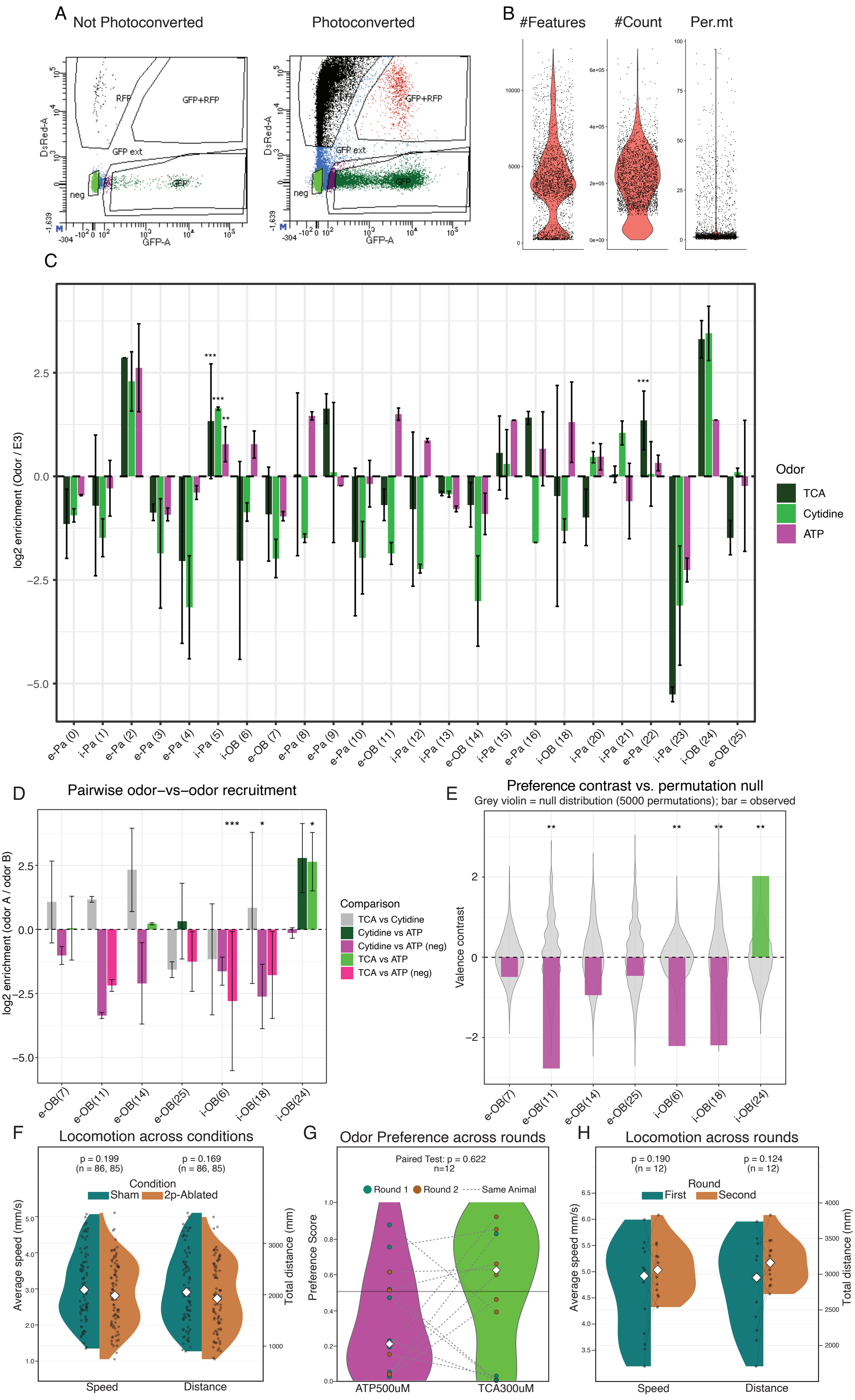
